# Rewriting regulatory DNA to dissect and reprogram gene expression

**DOI:** 10.1101/2023.12.20.572268

**Authors:** Gabriella E. Martyn, Michael T. Montgomery, Hank Jones, Katherine Guo, Benjamin R. Doughty, Johannes Linder, Ziwei Chen, Kelly Cochran, Kathryn A. Lawrence, Glen Munson, Anusri Pampari, Charles P. Fulco, David R. Kelley, Eric S. Lander, Anshul Kundaje, Jesse M. Engreitz

## Abstract

Regulatory DNA sequences within enhancers and promoters bind transcription factors to encode cell type-specific patterns of gene expression. However, the regulatory effects and programmability of such DNA sequences remain difficult to map or predict because we have lacked scalable methods to precisely edit regulatory DNA and quantify the effects in an endogenous genomic context. Here we present an approach to measure the quantitative effects of hundreds of designed DNA sequence variants on gene expression, by combining pooled CRISPR prime editing with RNA fluorescence *in situ* hybridization and cell sorting (Variant-FlowFISH). We apply this method to mutagenize and rewrite regulatory DNA sequences in an enhancer and the promoter of *PPIF* in two immune cell lines. Of 672 variant-cell type pairs, we identify 497 that affect *PPIF* expression. These variants appear to act through a variety of mechanisms including disruption or optimization of existing transcription factor binding sites, as well as creation of *de novo* sites. Disrupting a single endogenous transcription factor binding site often led to large changes in expression (up to –40% in the enhancer, and –50% in the promoter). The same variant often had different effects across cell types and states, demonstrating a highly tunable regulatory landscape. We use these data to benchmark performance of sequence-based predictive models of gene regulation, and find that certain types of variants are not accurately predicted by existing models. Finally, we computationally design 185 small sequence variants (≤10 bp) and optimize them for specific effects on expression *in silico*. 84% of these rationally designed edits showed the intended direction of effect, and some had dramatic effects on expression (–100% to +202%). Variant-FlowFISH thus provides a powerful tool to map the effects of variants and transcription factor binding sites on gene expression, test and improve computational models of gene regulation, and reprogram regulatory DNA.

## Introduction

Reading and writing the regulatory code of gene expression is a fundamental challenge in genome biology and medicine. This regulatory code, including collections of binding sites for transcription factors in *cis*-regulatory elements such as enhancers and promoters, specifies where and when genes turn on in different cell types in the body. Human genetics studies have now discovered hundreds of thousands of DNA variants in *cis*-regulatory elements that influence risk for common and rare diseases, each of which could point to new genes and cell types involved in pathogenesis^1–3^. Reprogramming *cis-*regulatory elements could enable treating diseases via cell type-specific modulation of gene regulation, as highlighted by the recent approval of an enhancer-targeting CRISPR therapy for sickle cell disease^4,5^.

To unlock these applications, we need an experimental approach to rapidly introduce many arbitrary sequence variants into a desired locus in the human genome and measure their quantitative effects on the expression of nearby genes. Such a method would accelerate our understanding of the rules and programmability of regulatory elements—for example, by enabling experiments to directly dissect transcription factor binding sites in an endogenous genomic context, to generate gold-standard data to train or evaluate predictive models of regulatory DNA^6–11^, and to iteratively test tools to reprogram gene expression for therapeutics.

For many years, the typical approach for studying such effects in an endogenous genomic context has been to study one variant or transcription factor binding site at a time. This approach involves introducing edits into a population of cells (*e.g.*, using homologous recombination), isolating many individual clones, genotyping to find homozygous edited cells, and then measuring the effects on gene expression. This process takes many months to characterize even a single variant, is subject to large clone-to-clone variation, and as such only a handful of sequence edits have been studied at a time (*e.g.*, ^12–14)^.

Existing high-throughput technologies have honed our understanding of this regulatory code, but have important limitations. Massively parallel reporter assays have revealed how transcription factor binding sites combine to drive enhancer and promoter activity in plasmids^15–22^, but do not properly model key aspects of genomic context. CRISPR nucleases and base editing have been coupled with single-cell or sorting-based readouts to identify and dissect regulatory elements^5,23–31^, but lack the flexibility to precisely reprogram regulatory sequences, depend on fortuitous positioning of editing sites to disrupt or introduce transcription factor binding sites, and often produce multiple possible edits per gRNA that lead to challenges in data interpretation^32^. The development of CRISPR prime editing, which provides the ability to precisely delete or insert designed sequences up to dozens of base pairs^33^, promises to address some of these challenges. However, the efficiency of prime editing remains limited, and pooled screening methods to date^34–36,99^ do not enable direct readouts of effects on gene expression (**Supplementary Table 1**).

To address these challenges, we developed an experimental method to measure the quantitative effects of hundreds of designed edits to endogenous regulatory DNA directly on gene expression. This method combines pooled prime editing—in which we introduce many programmed insertions or deletions into a population of cells—with RNA fluorescence *in situ* hybridization (RNA FISH) and flow sorting (Variant-FlowFISH), to directly measure effects on gene expression. To demonstrate this approach, we systematically dissect and reprogram the expression of *PPIF* (encoding cyclophilin D), a gene that influences genetic risk for autoimmune and inflammatory diseases^2^. Through a combination of tiling mutagenesis, transcription factor motif insertions, and computationally optimized sequence edits, we quantify the effects of 672 variants in the *PPIF* promoter and a distal enhancer that reveal a highly tunable and programmable regulatory landscape. By overcoming key barriers in the study of regulatory DNA, this method will provide insight into the sequence logic of gene regulation and enable developing genome editing reagents that achieve desired gene expression outcomes.

### Variant-FlowFISH enables pooled measurements of effects of sequence edits on gene expression

We developed a high-throughput technology called Variant-FlowFISH to study the effects of designed sequence edits on the expression of a gene of interest (**Fig. 1a**). Our approach starts with pooled CRISPR prime editing, in which we encode designed edits in a lentiviral pool of prime editing guide RNAs (pegRNAs) to mutagenize, replace, or rewrite regulatory DNA sequences at a selected locus. We transduce this lentiviral pegRNA pool into a population of cells expressing the CRISPR prime editor (PE2 system^33^, nCas9 fused to Moloney murine leukemia virus reverse transcriptase (MMLV) from an inducible promoter, and add doxycycline to activate the prime editing machinery. After a period of editing (here, 14+ days), we culture cells without doxycycline for at least 7 days to allow nCas9-MMLV to degrade, and thereby ensure that binding or active editing does not itself interfere with gene regulation at the targeted site. We then fluorescently label cells based on their expression of a gene of interest using RNA FISH, and use fluorescence-activated cell sorting (FACS) to sort cells into bins based on their expression levels of the target gene (FlowFISH)—allowing us to directly assay effects on mRNA expression while avoiding the need for antibodies or specialized reporters for a given gene of interest^37^. Finally, we extract genomic DNA from the cells, PCR-amplify the edited site, and use high-throughput sequencing to determine the frequency of edits in each of the expression bins (**Methods** & **Supplementary Fig. 1**). By directly sequencing the edited site, we can detect edits introduced even at low frequencies.

**Figure 1.**
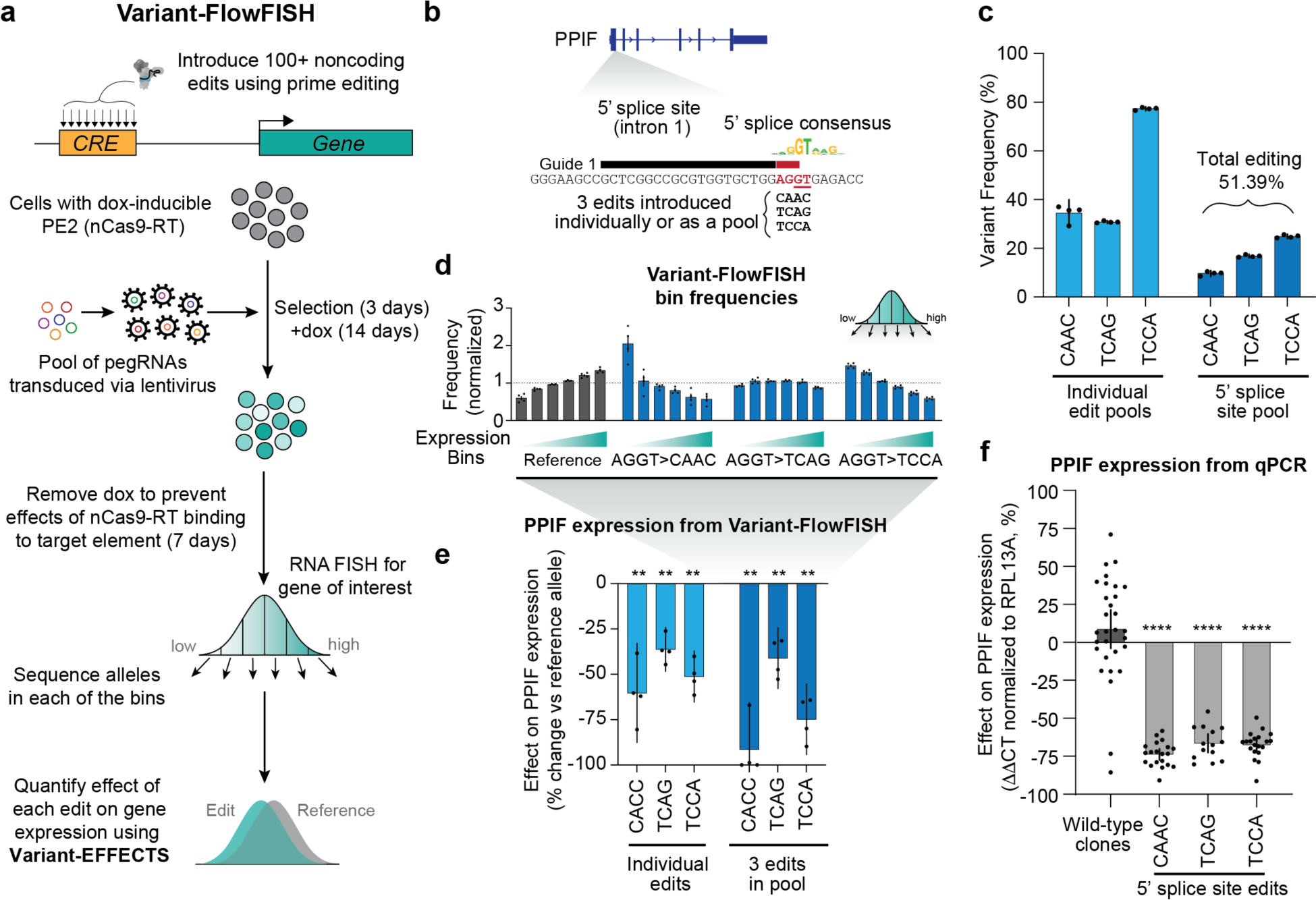
Variant FlowFISH combines prime editing with RNA-FlowFISH to investigate the effects of noncoding variants on expression of its target gene. **(a)** Overview of the Variant-FlowFISH pipeline. We introduce 100+ sequence variants targeting a *cis*-regulatory element (CRE) of interest into a pool of cells using CRISPR prime editing. Lentivirus is used to introduce pegRNAs into cells expressing PE2 prime editor from a doxycycline-inducible promoter. Successfully infected cells are selected with puromycin, and prime editing is activated with treatment with doxycycline. Cells are stained for an RNA of interest and sorted into bins based on expression (FlowFISH). We PCR amplify and sequence the edited site, measure the frequencies of each edit (allele) in each expression bin, and estimate the quantitative effect of the edit on gene expression using Variant-EFFECTS (see also **Supplementary Fig. 1**). **(b)** Prime editing strategy used to disrupt the 5’ splice site of the first intron of *PPIF*. Thick line: Location of the pegRNA spacer (black) and protospacer adjacent motif (PAM, red). Nucleotides to be replaced are highlighted in red, the critical ‘GT’ dinucleotide essential for splicing is underlined, and the 3 edit sequences are below in black. The 5’ splice site consensus motif is also shown^38^. **(c)** Frequency of each variant in cells after 13 days of prime editing activation with doxycycline treatment, prior to cell sorting, as measured by amplicon sequencing of the edited site. Dots: 2 technical FlowFISH replicates from each of 2 biological replicates (*n*=4). Bars: mean +/-95% confidence interval (c.i.). **(d)** Relative frequency of each allele (reference, and 3 edits) for the 3-pegRNA pool in each of 6 FlowFISH expression bins. Frequencies are normalized to the mean frequency of the reference allele across all 6 bins. Bars and dots as in **c**. **(e)** Effects of 5’ splice site edits on *PPIF* expression, as measured by Variant-FlowFISH (% effect versus the reference allele). Bars and dots as in **c**. **: *p* < 0.01, one-sample, two-tailed t-test. **(f)** Effects of 5’ splice site edits on *PPIF* expression, as measured by qPCR in clonalized cell lines homozygous for each edit. Dots: Clones for wild-type (*n*=30), AGGT>CACC (*n*=20), AGGT>TCAG (*n*=14), and AGGT>TCCA (*n*=20). Bars: mean effect +/- 95% c.i. ****: *p* < 0.0001, one-sample, two-tailed t-test.

We developed a mathematical approach and computational pipeline (Variant-EFFECTS: Variant-Estimation For Flow-sorting Effects in CRISPR Tiling Screens) to estimate the quantitative effect of each edit based on these frequency measurements, considering editing efficiency and cell ploidy. Because prime editing is not 100% efficient, we expect a mix of cells carrying homozygous and heterozygous edits, which will have different expression levels of the gene of interest. Accordingly, Variant-EFFECTS infers the effects of edits on gene expression by adjusting our previous maximum likelihood estimation procedure^37^ to account for a distribution of genotypes in the population of cells carrying 0, 1, or 2 alleles with the intended edit (see **Supplementary Fig. 1** and **2**). Notably, this estimation procedure assumes that, in diploid cells with two alleles of the targeted site, (i) the editing of each allele in a cell is independent of the other, which appears to fit well data for several individual variants with varying allele frequencies (*e.g.*, **Supplementary Fig. 3**), and (ii) a single cell does not receive two different edits, which holds for PE2 prime editing due to its precision in installing the intended edit^33^. Crucially, we found that methods that introduce frequent unintended edits, such as CRISPR homology-directed repair, base editing, or PE3 prime editing, confound Variant-FlowFISH or similar pooled screens because the presence of such edits can create artificial correlations that lead to both false positives and false negatives (**Supplementary Fig. 2**). Such artificial correlations represent a technical pitfall that Variant-FlowFISH circumvents by using PE2 prime editing, as opposed to the more efficient but less precise PE3 system,^33^ and by reading out the effects of variants directly by sequencing of the edited site, as opposed to reading out variants indirectly via sequencing of the pegRNA (**Supplementary Fig. 2**).

To demonstrate the Variant-FlowFISH protocol, we designed a proof-of-concept study to introduce sequence edits that should strongly reduce expression of a target gene of interest, *PPIF* (**Fig. 1b**). In macrophages, *PPIF* is involved in tuning mitochondrial membrane potential and pro-inflammatory signaling, and we have previously conducted *PPIF* FlowFISH screens in combination with CRISPR interference in several immune cell lines.^2^ We designed three pegRNAs with edits to disrupt the ‘GT’ splice donor at the first 5’ splice site (**Fig. 1b, Supplementary Table 2**), which we expected to lead to strong decreases in mRNA levels due to aberrant splicing and subsequent nonsense mediated decay. We transduced populations of THP-1 monocyte PE2 cells either with each pegRNA individually or with a pool of all three pegRNAs, sequenced the edited site, and observed editing rates of 34-77% per individual edit, or 51% total editing for the pool (**Fig. 1c**). We performed Variant-FlowFISH on each population of cells and quantified the frequency of variants across 6 expression bins. Compared to the reference sequence, all three edits showed higher frequencies in the low expression bins and lower frequencies in the high expression bins, indicating that they reduce *PPIF* expression (**Fig. 1d, Supplementary Fig. 3a**). With Variant-EFFECTS, we converted these measured frequencies to quantitative effect size estimates: single edits led to a 36-60% decrease in *PPIF* expression, with similar effects in the 3-pegRNA pool (**Fig. 1e, Supplementary Fig. 3c**). To compare these Variant-FlowFISH measurements to an orthogonal approach, we genotyped and derived homozygous clonal cell populations (**Supplementary Fig. 3**), performed qPCR on 84 clonal cell lines, and found that all 3 edits indeed decreased *PPIF* expression by similar amounts (**Fig. 1f**).

### Tiling mutagenesis of regulatory elements in the human genome

We next explored the utility of Variant-FlowFISH in mapping the functions of regulatory DNA sequences in their endogenous locations in the genome through high-throughput tiled mutagenesis. In particular, the identity, positions, and effect sizes of transcription factor binding sites in enhancers and promoters have been difficult to experimentally map. Accordingly, we selected the *PPIF* promoter and a distal enhancer of *PPIF* and designed experiments to systematically identify sequences important for regulating *PPIF* gene expression in THP-1 monocytic cells (**Fig. 2a**). At the *PPIF* promoter, we tiled edits across a 220-bp region spanning - 198 to +22 bp relative to the transcription start site (TSS). At the selected distal enhancer, located ∼60.5 Kb upstream of *PPIF*, we previously found that CRISPRi perturbations reduced *PPIF* expression by 37%^2^, and here tiled edits across a 175-bp region of interest (**Fig. 2a**).

**Figure 2.**
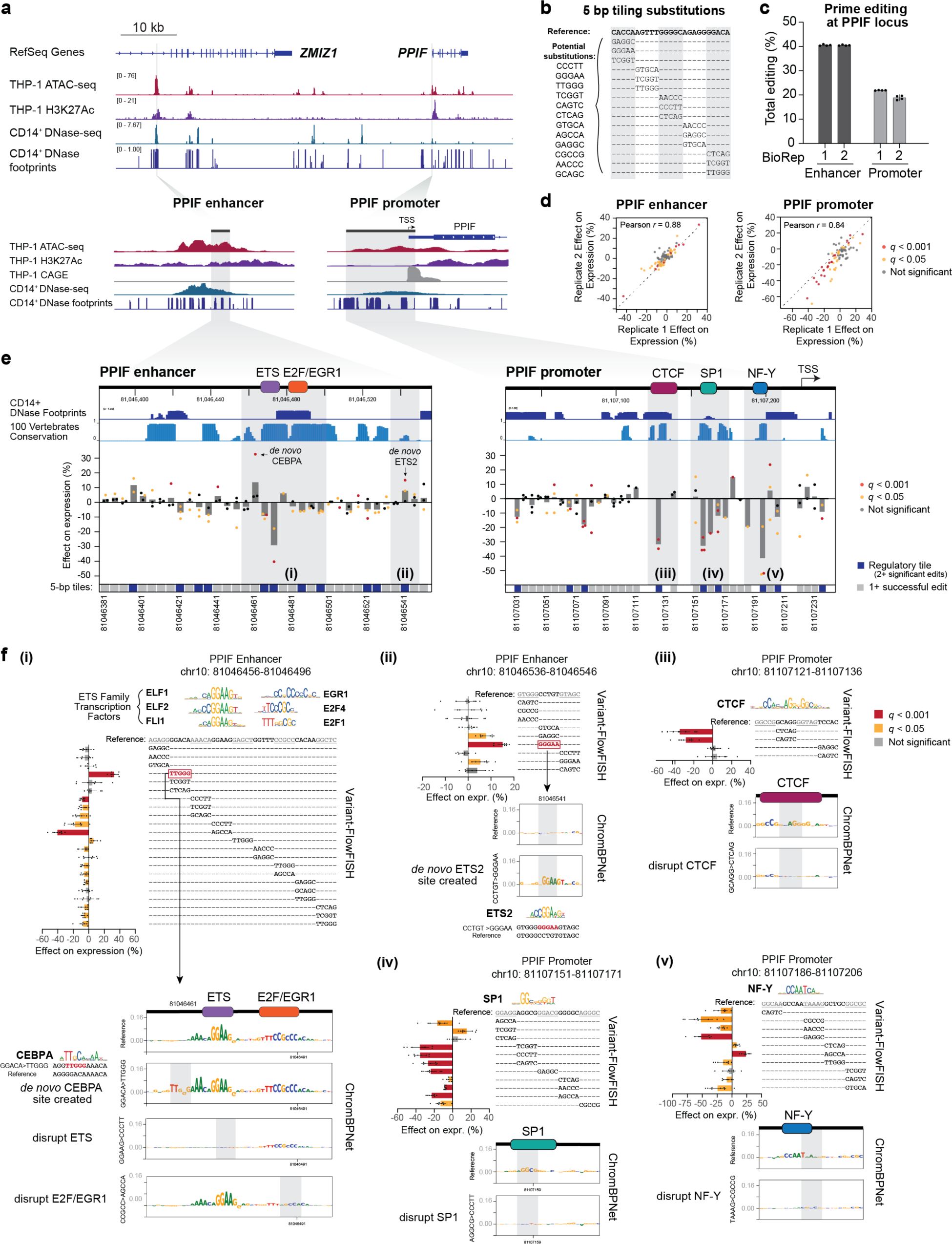
Tiling mutagenesis of an enhancer and promoter for *PPIF*. (**a**) Dissecting the regulation of *PPIF* via tiling mutagenesis of the promoter and a distal enhancer in THP-1 monocytic cells. Chromatin state signal tracks show data from THP-1 (ATAC-seq and H3K27ac) and the corresponding primary cell type CD14^+^ monocytes (DNase-seq and DNase footprints). Gray highlights show the regions for tiling mutagenesis. Cap Analysis of Gene Expression (CAGE) reads mark the TSS. Coordinates: *PPIF* locus (hg19 chr10:81,037,448-81,124,761), *PPIF* enhancer (chr10:81,045,489-81,047,143), *PPIF* promoter (chr10:81,106,967-81,107,535). (**b**) We conducted tiling mutagenesis in 5-bp windows across each regulatory element, and selected substitutions from a bank of 12 possible sequences selected for prime editing efficiency (Methods). (**c**) Total editing (% of sequencing reads, summed across all designed edits) for Variant-FlowFISH screens at the *PPIF* enhancer and *PPIF* promoter. Dots: technical FlowFISH replicates (*n*=4). Bar: mean +/-95% c.i. (**d**) Variant-FlowFISH measurements of variant effects on *PPIF* expression (%) are highly correlated between two biological replicates. Dots: all variants passing the frequency threshold for the enhancer and promoter tiling screens. Red: *q* < 0.001. Yellow: *q* < 0.05 (Benjamini-Hochberg corrected *p*-value, one-sample t-test). (**e**) Tiling mutagenesis data at the *PPIF* enhancer and promoter in THP-1. Dots: Effect of each 5-bp substitution on *PPIF* expression, as measured by Variant-FlowFISH (mean of 2 biological replicates x 4 technical replicates). Bars: Mean of 1-3 substitutions at each position. Variants with significant effects are highlighted in yellow (*q* < 0.05) and red (*q* < 0.001). Tracks at top show CD14^+^ DNase footprints and evolutionary conservation across 100 vertebrates (PhastCons). Bottom: Dark blue indicates “regulatory tiles” (positions with 2 or more significant variants with the same direction of effect), gray indicates a tested tile, and white indicates a tile with no edits of sufficiently high frequency. Gray highlights show regions of interest. Colored boxes at top: Positions of transcription factor binding sites identified by Variant-FlowFISH and motif analysis. Genomic coordinates: *PPIF* enhancer (chr10: 81,046,381-81,046,556) and *PPIF* promoter (chr10: 81,107,026-81,107,246). (**f**) Variant-FlowFISH data and ChromBPNet predictions at selected regulatory tiles (gray highlights in **e**). Barplots show effects on *PPIF* expression as measured by Variant-FlowFISH (bars: mean +/-95% c.i; dots: replicate experiments, *n*=6-8; yellow bars: *q* < 0.05; red bars: *q* < 0.001). Motifs (identified by FIMO from MEME Suite using the HOCOMOCO v11 database and JASPAR^56,57^) show potential transcription factors binding sites disrupted or created by 5-bp edits. Substitutions highlighted in red indicate the creation of a *de novo* binding site. ChromBPNet sequence interpretations (DeepSHAP) of the reference and edited sequences show the predicted contribution of each nucleotide for chromatin accessibility signal. Gray boxes within the ChromBPNet sequence interpretations highlight the position of selected 5-bp edits.

To identify endogenous regulatory motifs, we designed pegRNAs to introduce 5-bp substitutions tiled end-to-end across each element (**Fig. 2b**). At each 5-bp tile, we substituted three different sequences to account for the possibility that any one substitution could unintentionally create new regulatory sequences, such as novel transcription factor binding sites. We designed 105 pegRNAs for the *PPIF* enhancer and 132 pegRNAs for the *PPIF* promoter (**Supplementary Fig. 4, Supplementary Table 3,** see Methods for more details on pegRNA design) and transduced each pool separately into THP-1 cells. We observed a total editing rate of 40% and 20% for the 5-bp substitutions at the *PPIF* enhancer and promoter, respectively (**Fig. 2c**). Editing efficiency for individual variants varied, ranging from an estimated 0.001–100% per substitution (estimated as the proportion of cells with a variant divided by the proportion of the corresponding pegRNA in the plasmid library, **Supplementary Fig. 4f**). We focused on the 184 of 237 designed variants, across the *PPIF* enhancer and promoter, present at >0.01% frequency in the final pool of cells (**Supplementary Fig. 4e**), allowing us to assay >2,000 cells per variant per Variant-FlowFISH replicate in which we sort a total of 20 million cells. Aggregating across tiles, we obtained sufficient editing for at least one edit for 87% of targeted 5-bp tiles, including all 35 tiles at the *PPIF* enhancer and 34 of 44 tiles at the *PPIF* promoter (**Supplementary Fig. 4d**). This variability in editing efficiency is consistent with previous studies^33,39–41^, and may be explained by differences in transcription factor occupancy and/or features of the pegRNA (**Fig. 2a, Supplementary Fig. 4d,g**).

We performed Variant-FlowFISH for each pool of edited cells, and obtained data that were strongly correlated between biological replicates, both at the level of variant frequencies in each sorting bin (average Pearson *r* = 0.989-0.998 and 0.977-0.985 for the enhancer and promoter, respectively, **Supplementary Fig. 5a,b**) and at the level of effect sizes on gene expression from Variant-EFFECTS (average Pearson’s *r* = 0.88 and 0.84 for enhancer and promoter pools, respectively (**Fig. 2d**)). We tested whether each variant altered gene expression versus the reference allele by comparing across 2 biological replicates, each with 4 FlowFISH technical replicates (**Supplementary Fig. 1b**).

In total, we identified 93 variants with significant effects on *PPIF* expression (t-test with Benjamini-Hochberg corrected *p*-value (*q*) < 0.05; 49 variants for the enhancer and 44 at the promoter), located at 54 distinct 5-bp tiles, with effect sizes ranging from –52% to +32% (**Fig. 2e and Supplementary Fig 5c**). By aggregating results per tile, we defined 20 of 79 tiles as “regulatory tiles” based on ≥2 substitutions at that tile having significant effects on expression in the same direction (either increasing or decreasing *PPIF* expression; 11 tiles at the enhancer and 9 at the promoter) (**Fig 2e**).

Many of the regulatory tiles with the largest effect sizes on gene expression corresponded to DNase-seq footprints in CD14^+^ monocytes (a closely related cell type), to motif instances of transcription factors involved in regulation of mitochondrial genes like *PPIF*, and to sequences predicted to affect chromatin accessibility or gene expression by computational models such as Enformer^6^ and ChromBPNet^42^ (see below) (**Fig. 2e**):

For example, at the *PPIF* enhancer, Variant-FlowFISH identified a central region containing 4 regulatory tiles in a span of 35-bp where edits led to a significant decrease in *PPIF* gene expression (**Fig. 2f**(i)). The variants with the largest effect sizes fell within a predicted ETS family transcription factors motif: two variants that disrupted the core ‘GGAA’ of this motif led to an average -29% decrease in PPIF gene expression, and variants that disrupted the flanking ‘CA’ nucleotides immediately upstream led to an average -9% decrease. Adjacent to this ETS motif, substitutions disrupting predicted motif instances for E2F1, E2F4, and/or EGR1 led to an average -6% decrease in *PPIF* gene expression. Notably, ETS family factors, E2F factors, and EGR1 are transcriptional activators that are expressed in THP-1 cells (**Supplementary Fig. 6**) and are known to regulate genes that, like *PPIF*, are involved in mitochondrial function^43^. We trained a deep learning model (ChromBPNet^42^) to predict chromatin accessibility in THP-1 and annotated the predicted contribution of each base pair in this region using backpropagation with DeepLIFT/DeepSHAP^44,45^ (**Supplementary Fig. 7**). Disruptions to the ETS or E2F/EGR1 motifs were indeed predicted to decrease chromatin accessibility, consistent with the observed decrease in *PPIF* gene expression (**Fig. 2f**(i)). At the *PPIF* promoter, similar analysis identified predicted binding sites for CTCF (-35% decrease), SP1 (-36% decrease), and NF-Y (-52%) that were clustered in the region approximately –100 to -25 from the TSS (**Fig. 2f**(iii)-(v)).

In addition to effects of endogenous regulatory sequences, we identified cases where substitutions appeared to create *de novo* binding sites for transcriptional activators. For example, a substitution near the ETS motif instance in the enhancer increased *PPIF* gene expression by +32% and was predicted to create a *de novo* motif instance for CEBPA (**Fig. 2f**(i)). As another example, a substitution that increases *PPIF* gene expression by +15% was predicted to create a second ETS motif instance *de novo* (**Fig. 2f**(ii)).

Collectively, the distribution of functional sequences and effect sizes across the *PPIF* enhancer and promoter highlighted key properties of human regulatory elements:

i. The frequency and spacing of functional nucleotides appeared to match previous genetic studies of individual regulatory elements in animal models or plasmid-based reporter assays^15–22,46–54^. 28% of tested 5-bp tiles had consistent regulatory effects (2 or more edits in the same direction), and were interspersed with tiles that appeared to have subtle or no effect on expression (**Fig. 2e**).
ii. The effect sizes of disrupting or adding individual transcription factor binding sites at the enhancer were remarkably strong, and comparable to effects at the promoter (range: -40% to +32% at the enhancer and -52% to +23% at the promoter) (**Fig. 2e, Supplementary Figure 5c**). For example, removal of the central ETS motif instance resulted in an average -29% effect, and creation of an adjacent CEBPA motif instance resulted in a +32% increase in expression (**Fig 2f**(i)). Given that *PPIF* is already highly expressed (463 TPM), this represents a large increase in *PPIF* mRNA from small sequence changes in a distal enhancer.
iii. The magnitude of effects suggests that these transcription factor binding sites are likely to act super-additively with respect to linear gene expression, consistent with previous observations in other regulatory elements (*e.g.*, ^17,55^). For example, at the *PPIF* enhancer, disrupting just this one ETS motif instance in the enhancer (-29% decrease) leads to an effect similar in magnitude to CRISPRi repression of the entire enhancer element (-37% decrease^2^), and the sum of the effects of all regulatory tiles which decreased expression was 78% (**Fig 2e**). Similarly, at the *PPIF* promoter, the sum of the effects from disrupting the 4 individual putative transcription factor binding sites, CTCF (-35%), SP1 (-36%) and NF-Y (-52%), exceeds a 100% decrease in *PPIF* gene expression (**Fig. 2e**), consistent these sites acting super-additively with respect to gene expression.
iv. Across just 345 bp of tested DNA sequence, we identify 4 distinct sites corresponding to different transcriptional activators (ETS, E2F/EGR1, NF-Y and SP1; **Fig. 2e**) that have been reported to regulate genes that, like *PPIF*, are involved in mitochondrial metabolism^43^. This suggests that in THP-1 cells, *PPIF* may be programmed to respond to a large number of different transcription factor inputs, and indeed its expression pattern has been observed to be highly dynamic across time during THP-1 activation and across cell types^2^.

Altogether, these tiling mutagenesis experiments reveal a highly tunable regulatory landscape around *PPIF*, and illustrate the power of designed mutagenesis to identify the sequence architecture of regulatory elements in their endogenous contexts.

### Inserting transcription factor binding sites

Beyond studying endogenous regulatory sites, Variant-FlowFISH could enable the study of designed regulatory sequences—for example, to dissect the context-specific effects of transcription factor binding sites or reprogram genes to respond to particular signaling pathways. To explore this application, we selected a site in the *PPIF* promoter where prime editing was highly efficient (-58 bp relative to TSS, estimated 76% editing efficiency in the promoter tiling screen), and designed a library of 41 pegRNAs to insert 8-bp synthetic sequences that included predicted binding sites for transcription factors expressed in THP-1, or differentially expressed between THP-1 and another immune cell line (Jurkat T cells) (**Fig. 3a, Supplementary Tables 3,4 & Methods**).

**Figure 3.**
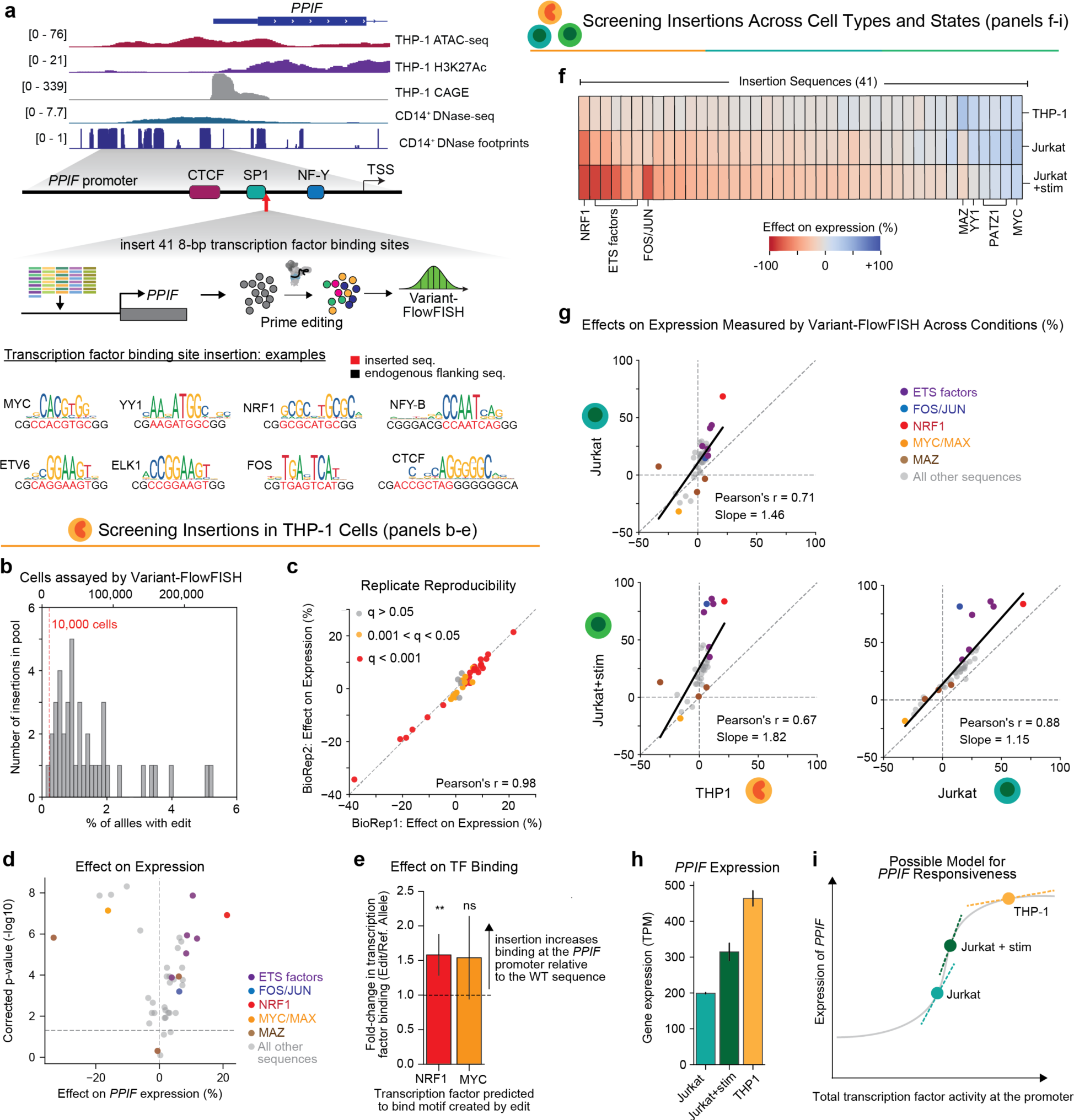
Inserting transcription factor binding sites at the *PPIF* promoter. (**a**) We inserted a library of 41 8-bp DNA sequences at a site 58 bp upstream of the *PPIF* TSS in THP-1 monocytes. (**b**) Histogram of frequencies of each 8-bp insertion in the edited pool of cells as a percentage of alleles (bottom x-axis) and the corresponding minimum number of cells assessed by Variant-FlowFISH (top x-axis). (**c**) Correlation of effect sizes on gene expression between 2 Variant-FlowFISH biological replicates. (**d**) Variant-FlowFISH measurements of variant effects on *PPIF* expression (%) in THP-1 cells are highly correlated between two biological replicates. Dots (*n*=41): all variants passing the frequency threshold. Red: *q* < 0.001. Yellow: *q* < 0.05, gray: *q* > 0.05 (Benjamini-Hochberg corrected *p*-value, one-sample t-test). (**e**) Change in the binding of MYC or NRF1 relative to the reference allele for 8-bp insertions that create binding motifs for these factors, as measured by ChIP followed by amplicon sequencing. Allele-specific fold change is calculated by comparing the frequencies of the edit and reference alleles in the ChIP sample versus whole-cell genomic DNA input (Methods). **: *p* = 0.007, one sided t-test. ns: not significant. (**f**) Effects of the 8-bp sequence insertion library were measured in three cellular conditions (rows) using Variant-FlowFISH: in THP-1 monocytes, Jurkat T cells, and Jurkat T cells stimulated with PMA and anti-CD3 antibody. Heatmap color: Effect of each insertion (columns) on *PPIF* expression, relative to the reference allele within each condition. (**g**) Pairwise comparison of effects of 8-bp insertion edits among the three cellular conditions. Black line: Linear regression line of best fit. Dots (*n*=41): all variants passing the frequency threshold. Dots are colored if the edit creates a predicted *de novo* motif instance of a transcription factor binding site (see legend). (**h**) *PPIF* expression measured by RNA-seq in wild-type cells in transcripts per million (TPM). (**i**) A simple model that could explain the differences in the magnitude of effects observed between cell types. Dose response curve shows a hypothetical relationship between total transcription factor input to the *PPIF* promoter (x-axis) and *PPIF* expression (y-axis). Points along this curve start with different levels of transcription factor activity at the promoter (*e.g.*, due to factors binding at the promoter or distal enhancers). Dotted tangent lines represent how gene expression might vary due to changes in total transcription factor activity from 8-bp sequence insertions, and their slopes are a theoretical representation of the effects observed across conditions (see Fig. 3g).

We introduced these 41 insertions in THP-1 monocytes and evaluated their effect on *PPIF* expression using Variant-FlowFISH. Pooled prime editing achieved an average frequency of 1.6% per edit (range: 0.2 -5.2%, all edits above frequency threshold of 0.01%), and >10,000 single cells were assayed for 40 out of 41 edits (**Fig. 3b**). At this coverage, we had >80% power to detect an effect size of 10% for all 40 edits at a cell coverage threshold of 10,000 cells, and 28 edits had >80% power to detect an effect size of 5% (**Supplementary Fig. 8)**. The observed effect sizes for each insertion were highly correlated among technical and biological replicates (average Pearson’s *r* = 0.93 and 0.96, respectively), and 37 edits (90%) significantly altered *PPIF* expression in THP-1 monocytes (**Fig. 3c,d**; **Supplementary Fig. 9**).

These 8-bp insertions induced a wide range of effect sizes on *PPIF* expression, including edits that either significantly increased or decreased gene expression relative to the wild type promoter (range: –33% to +21%, **Fig. 3d**). Notably, the effect sizes of the sequence insertions were uncorrelated with common heuristics for the global activities of corresponding transcription factors, including with transcription factor mRNA expression, enrichment of these motifs in ATAC-seq peaks in THP-1 cells, and the average predicted global effect of these motifs on chromatin accessibility (as computed by ChromBPNet and TF-MoDISco; (**Supplementary Fig. 10, Supplementary Table 5** & **Methods**)^42,58^.

The strongest activating edit corresponded to an insertion that is predicted to create a binding site for the cell-essential transcription factor NRF1 (**Fig. 3a**; +21% effect; *q* = 1.21×10^-7^), followed by binding sites for ETS family transcription factors (*e.g.*, ELK1, ETS1)^59^. Both NRF1 and ETS factors are known to be strong activators^55,60–62^ (see also **Fig. 2**). The sequence insertions that led to the strongest decreases in gene expression corresponded to motif instances for MAZ, YY1, and MYC/MAX (-33%, -19%, and -16%, respectively; *q* < 1.5×10^-6^ for all), all three of which have been reported to have either activating and repressive functions in different contexts^63–68^ (**Fig. 3d**). MAZ has recently been found to promote the insulating activity of CTCF, suggesting the possibility that this inserted MAZ site may affect *PPIF* expression in coordination with the CTCF binding site located 37-bp away^69^. Both YY1 and MYC have been reported to reduce expression by interfering with binding of or activation by SP1 and NF-Y^70,71^, and indeed our tiling mutagenesis screen indicated that the *PPIF* promoter contains motif instances for SP1 and NF-Y that have strong effects on *PPIF* expression (**Fig. 3a**, **Fig. 2e,f**). To verify that effects on *PPIF* expression were associated with increased recruitment of target transcription factors, we individually introduced two selected variants (for NRF1 and MYC/MAX motif instances) and performed chromatin immunoprecipitation (ChIP) paired with amplicon sequencing and allele-specific quantification for the corresponding transcription factors. Insertion of an NRF1 or MYC/MAX motif instance increased the *PPIF* promoter ChIP-seq signal of NRF1 or MYC respectively, by 1.5-fold relative to the reference allele (t-test, *p* = 0.007 and 0.056, respectively; **Fig. 3e**).

We next sought to compare the effects of these 8-bp insertions across cell types and states. We used the same 41-pegRNA pool to conduct Variant-FlowFISH screens for *PPIF* in Jurkat T-cells with and without stimulation (phorbol 12-myristate 13-acetate (PMA) + anti-CD3 antibody). We again obtained highly reproducible quantification of variant effect sizes (biological replicate average Pearson’s *r* = 0.98 and 0.96 for with and without stimulation, respectively), and found 38 variants to have significant effects on *PPIF* expression in both of these Jurkat conditions (**Supplementary Fig. 11**). Nearly all of the variants showed significantly different effects between at least two of the three conditions (≥36 variants between each pair of conditions, **Supplementary Fig. 12**).

Some of these differences in variant effects appeared to be explained by differences in the global activity of the corresponding transcription factor. For example, the sequence variant that showed the strongest differential effect was a motif instance for FOS/JUN, which led to a +81% increase in *PPIF* expression in stimulated Jurkat cells versus +15% in unstimulated Jurkat cells (**Figure 3f**). The activity of the FOS/JUN family transcription factors that compose the AP-1 complex are known to be central to the chromatin remodeling and gene activation that occurs upon T-cell activation^72^. Indeed, we found *JUNB* to be strongly up-regulated upon stimulation in Jurkat T cells (**Supplementary Fig. 13, Supplementary Table 4**) and the FOS/JUN motif was highly correlated with chromatin accessibility signals in stimulated Jurkat cells (by ChromBPNet and TF-MoDISco, **Supplementary Table 7**). Thus, this sequence insertion effectively reprograms the expression of *PPIF* in Jurkat cells to be >60% more responsive to T-cell activation via introduction of a FOS/JUN motif instance in the promoter.

Yet, for most tested insertion sequences, the differences in effects on *PPIF* expression between two conditions was largely uncorrelated with differences in either the expression of corresponding transcription factors or their global enrichments in chromatin accessible regions (**Supplementary Fig. 14, Supplementary Tables 4,5,6**). Instead, effect sizes across cell types appeared to be driven by a systematic difference in the responsiveness of *PPIF* that affected all tested variants. Specifically, across the 3 conditions, the effects of all variants were highly correlated, but with different slopes (Pearson’s *r =* 0.71, 0.67, 0.88 and regression line of best fit slope = 1.46, 1.82, 1.15 for Jurkat vs. THP-1, Jurkat+stim vs. THP-1 and Jurkat+stim vs. Jurkat respectively; **Fig. 3g**). For example, insertion of an NRF1 motif instance led to a +21%, +68%, and +84% effect on gene expression in THP-1, Jurkat, and stimulated Jurkat cells, respectively. We validated the effects of the MYC, NRF1, and FOS/JUN insertions via clonal isolation and qPCR, and confirmed these cell type-specific differences in effect sizes on *PPIF* expression (**Supplementary Fig. 15**).

Together, these data indicate that transcription factor binding sites have cell type-specific effects, but that these differences might reflect not only cell type-specific differences in transcription factor activity (such as for FOS/JUN) but also systematic differences that can modulate the amplitude of effects of many different transcription factor binding sites. While there are many possible explanations, one simple model that could explain such differences is a dose-response relationship between, for example, total input transcription factor activity and gene expression (**Fig. 3h,i**). In this model, the regulatory activity at the *PPIF* promoter would start at a different position along a curve in each condition. A cell type with very high baseline *PPIF* expression, like THP-1 (**Fig. 3h**), would already be closer to a saturation point and so additional TF binding sites would lead to smaller effects on expression compared to a cell type that started with lower *PPIF* expression (**Fig. 3i**).

### Benchmarking sequence models of variant effects

Deep learning models have been developed to predict gene regulatory signals directly from DNA sequence and thereby interpret the effects of noncoding variants^6–11^. These models have yielded insights into the regulatory logic that influences different aspects of gene regulation, but their quantitative predictions have been difficult to evaluate due to lack of gold-standard experimental data in an endogenous genomic context.

To explore using Variant-FlowFISH data to benchmark such models, we examined two types of models: Enformer^6^, which uses a hybrid convolutional-transformer architecture to analyze long DNA sequence contexts up to 196 Kb in length, and models based on BPNet^8^ (ChromBPNet^42^ and ProCapNet^73^), which use a convolutional neural network (CNN) architecture to examine short, local DNA sequence contexts up to ∼2 Kb (**Fig. 4a**). We considered variations of these models trained to predict different gene regulatory signals—gene expression with capped analysis of gene expression (CAGE) (for Enformer), transcription initiation rates with PRO-cap (for ProCapNet), and chromatin accessibility with ATAC-seq or DNase-seq (for Enformer and ChromBPNet)—that might capture distinct aspects of the regulatory logic of noncoding DNA. We note that, based on data availability, certain models were trained on data directly from the THP-1 monocytic cell line (for Enformer CAGE model and ChromBPNet), and some models were trained on the most closely matched available cell types (CD14^+^ monocytes for Enformer DNase-seq model, K562 for ProCapNet) (see **Methods**).

**Figure 4.**
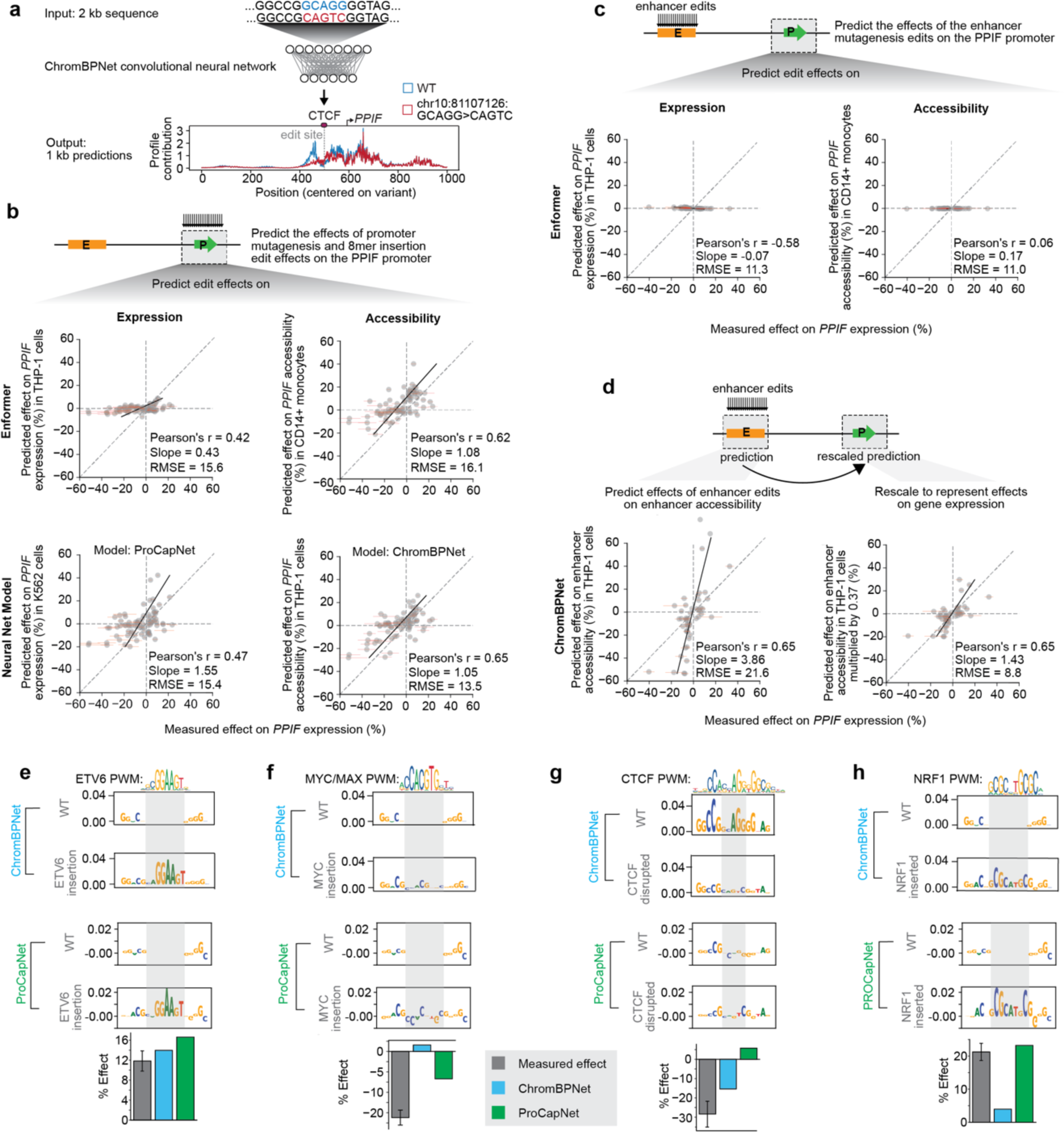
Benchmarking sequence-based predictive models of gene regulation. (**a**) Schematic of approach for calculating predicted effects of variants using ChromBPNet. ChromBPNet (or ProCapNet) takes as input 2 kb of DNA sequence and predicts base pair-resolution ATAC-seq (or PRO-Cap) profiles and counts. We calculate predicted effects as the difference in predicted counts between reference and edited 1 kb sequences, centered on the variant. Enformer (not shown) takes 196 kb of DNA sequence, and predicts CAGE or DNase signal in 128-bp bins. We calculate predicted effects as the difference in predicted signal between reference and edited 768-bp sequences (6 aggregated prediction bins), centered on the variant (for edits at the promoter) or the TSS (for predicting effects of edits at the enhancer on CAGE). See Methods for details on model predictions. (**b**) For promoter variants, comparison of measured effects on *PPIF* expression (Variant-FlowFISH) with predicted effects on either gene expression (left) or chromatin accessibility (right) at the promoter. Dots: *n*=82 variants at the *PPIF* promoter with significant effects on expression in THP-1 cells. Error bars: 95% c.i. for measured effect size. Black line: Linear regression line of best fit. Legend lists Pearson’s *r* correlation coefficient, slope from the linear regression, and root mean squared error (RMSE) of the predicted effects on expression (%). Predictions from Enformer (top row) and CNN models (ProCapNet and ChromBPNet, bottom) use data from THP-1 or the closest available cell type (Methods). (**c**) Similar to **b**, for *n*=50 edits with significant effects at the *PPIF* enhancer in THP-1 cells. Here, Enformer predicts effects of edits in the enhancer on CAGE or DNase-seq signals around the *PPIF* promoter (see Methods). (**d**) For enhancer variants with significant effects (n=50), we compared the measured effects on gene expression (Variant-FlowFISH) to predicted effects on chromatin accessibility at the enhancer (ChromBPNet, left). We then scaled these predicted effects on the enhancer by the measured effect of the enhancer on gene expression (37%), which we previously quantified using CRISPRi-FlowFISH, as a model for how effects on accessibility might affect gene expression (right). (**e-h**) DeepSHAP interpretations of base-resolution sequence contribution for ChromBPNet and ProCapNet predictions on (**e**) insertion of an ETV6 (ETS family) motif instance at the *PPIF* promoter, (**f**) insertion of a MYC/MAX motif instance at the *PPIF* promoter, (**g**) mutagenesis of an endogenous CTCF motif instance at the *PPIF* promoter, and (**h**) insertion of a NRF1 motif instance at the *PPIF* promoter. Transcription factor motif position weight matrices (PWMs) in e-h are from JASPAR^56^. Barplots show effects measured by Variant-FlowFISH (gray), effects predicted by ChromBPNet (light blue), and effects predicted by ProCapNet (green). Error bar: 95% c.i. of measured effect. For effects predicted by Enformer models, see Supplementary Table 8.

We examined to what extent these models can predict the effect sizes of DNA variants that significantly affected *PPIF* gene expression as measured by Variant-FlowFISH (**Fig. 4, Supplementary Table 8**). We evaluated performance using three metrics: the Pearson correlation between the measured and predicted effects, the slope of the line of best fit from linear regression (to assess whether predictions are calibrated with respect to the magnitude of the effects), and the root mean squared error (RMSE, to capture both correlation and calibration). Overall, performance differed considerably between models trained on expression versus accessibility, and on edits at the promoter versus edits at the enhancer. However, the best models obtained a correlation of up to 0.65 with slope close to 1 (**Fig. 4**):

For DNA variants introduced at the *PPIF* promoter (*n* = 82 significant variants in THP-1 combined across experiments), we first examined predicted effects on gene expression using Enformer (based on CAGE signals at the promoter in THP-1 cells), and ProCapNet (based on PRO-cap signals at the promoter in K562 cells; **Fig. 4b**). These predictions were positively correlated with measured effects (Pearson’s *r* = 0.42 and 0.47, respectively), although with slopes that indicated a degree of miscalibration (slope = 0.43 and 1.55, respectively). Interestingly, models that predicted effects on chromatin accessibility performed better, despite the fact that CAGE and PRO-cap are more direct readouts of gene expression: Enformer (DNase-seq in CD14^+^ monocytes) achieved a correlation of 0.62 with slope of 1.08, and ChromBPNet (ATAC-seq in THP-1 cells) achieved a correlation of 0.65 with slope of 1.05 (see **Discussion**).

We next assessed predictions for DNA variants introduced at the distal *PPIF* enhancer located 61 Kb upstream of the *PPIF* promoter (*n* = 50 significant variants, **Fig. 4c**). Only Enformer, by virtue of its large sequence context, is constructed to directly predict the effects of variants at this distance. However, Enformer performed poorly on this set of edits, as indicated by a low correlation with measured effects (Pearson’s *r* = -0.58 and 0.06 for CAGE and DNase-seq heads, respectively) and slopes close to 0 (**Fig. 4c**). Notably, performance was improved when we correlated measured effects on *PPIF* expression with Enformer predictions of effects on activity at the enhancer itself (Pearson’s *r* = 0.63 for CAGE and DNase-seq, **Supplementary Fig. 16**), indicating that Enformer learned the sequence logic for local effects but not the contribution of this distal enhancer for *PPIF* expression.

We considered an alternative approach to assessing the effects of enhancer variants on gene expression, based on the notion that variants in a distal enhancer should locally affect the activity of that enhancer, and that altered enhancer activity should be linearly related to gene expression based on the overall strength of the enhancer (**Fig. 4d, Supplementary Figure 16**). Accordingly, we used ChromBPNet and Enformer to predict effects of variants on accessibility at the *PPIF* enhancer, and multiplied by the measured effect of the entire enhancer on gene expression (37%), which we previously quantified using CRISPRi-FlowFISH^2^. These results yielded a significant improvement in the accuracy at predicting effect sizes of variants on gene expression. For example, this improved the slope of ChromBPNet predictions from 3.86 (predicted effect on chromatin accessibility at enhancer) to 1.43 (scaled effect to predict expression) (**Fig. 4d**; from slope = 2.36 to 0.87 for Enformer, **Supplementary Fig. 16**). This suggests that future modeling approaches could attempt to link distal variants to effects on gene expression through a two-step process: by (i) predicting effects of variants on enhancer activity and (ii) scaling by a prediction of the effect of the enhancer on gene expression.

To better understand the capabilities and limitations of current models for detecting effects of variants that influence expression through different mechanisms (*e.g.*, chromatin accessibility, transcription initiation), we examined variant predictions that were discordant with measurements or discordant across models. While the models recognized the effects of strong activating edits such as instances of ETS family motifs (**Fig. 4e**), other categories of edits appeared to be missed (**Fig. 4f**-**h**):

For example, insertion of a MYC/MAX binding site at the *PPIF* promoter in THP-1 cells decreased gene expression by 16% (**Fig. 3d,e**), but ProCapNet and Enformer underestimated this effect on expression and ChromBPNet predicted nearly no effect on accessibility (**Fig. 4f**). The lack of a predicted effect of MYC/MAX on accessibility is consistent with previous studies in reprogramming systems that have shown that MYC binding relies entirely on other pioneer transcription factors such as OCT4 and SOX2 to open chromatin even when MYC is overexpressed^74^, and we have previously observed that ChromBPNet does not detect an impact of MYC on accessibility in these reprogramming systems^75^. However, these inaccuracies in predictions of gene expression may also be due to difficulties in modeling MYC/MAX’s dual role as an activator or repressor depending on sequence context^67,75^.

The predicted direction of effect for edits to CTCF motif instances were often discordant between expression and accessibility models. For example, disruption of a CTCF binding site at the *PPIF* promoter decreased gene expression in THP-1 cells by 28% (**Fig. 2e**). ChromBPNet predicted a comparable effect on accessibility (-15%), but ProCapNet predicted an opposing positive effect on local transcription initiation rate (+6%; **Fig. 4g**). This may be because these models can learn the local effect of CTCF binding on accessibility, but not the regulatory logic of how CTCF binding would influence long-range 3D contacts to affect gene expression.

In some cases, sequence edits appeared to affect the logic of transcriptional initiation, rather than enhancer/promoter activity, in that they involved transcription factors known to affect transcriptional initiation and were more accurately predicted by ProCapNet than by any other model we tested. For example, insertion of an NRF1 motif was measured to have an +21% effect, which was accurately predicted by the the ProCapNet model of transcription initiation (predicted effect = +23%) but not by the CAGE model of steady-state gene expression (predicted effect = +3.3%) or models of chromatin accessibility (predicted effect = +4.0% and +7.1% for ChromBPNet ATAC and Enfomer DNase models, respectively) (**Fig. 4h, Supplementary Table 8**). This is consistent with prior work suggesting that NRF1 may influence transcriptional initiation based on the positional bias of its motif relative to transcription start sites^10^.

Together, these analyses illustrate how quantitative, gold-standard sequence perturbation data from Variant-FlowFISH can be applied to evaluate and identify limitations of sequence-based predictive models of different molecular readouts of gene regulation. Future model development could improve predictions for variants in distal enhancers, and combining the predictions of models trained on different epigenomic assays could help to improve interpretation of variants that affect gene regulation through distinct mechanisms.

### Reprogramming gene expression using designed sequence edits

The results of our synthetic DNA sequence insertion experiments led us to consider how reprogramming of noncoding regulatory DNA could be optimized to yield precise and tunable changes to cell type-specific gene expression, with potential applications in synthetic biology and gene therapy. In particular, while the experiments above yielded effects ranging from -53% to +86%, we tested whether we could achieve even larger changes in expression with CRISPR prime editing by rationally designing edits with predictive models.

We developed a computational approach to optimize sequence edits that are both tractable with prime editing and predicted to create desired gene expression outcomes. We first select a target site with efficient prime editing, randomly initialize a sequence insertion at that site, and then optimize the sequence of the edit over 1,000 steps using simulated annealing with a standard Metropolis acceptance criterion^76,77^. Each step involves insertion, deletion, or substitution of 1 nucleotide in the edit, or a 1 bp shift of the edit insertion site. We predict the effect of the edit after each step using a deep learning model (here, the Enformer CAGE heads, which were available for both THP-1 and Jurkat cells). Using the predicted increase in expression fold-change in each cell type (edited sequence/WT sequence), we made incremental 1 bp changes to the current edit to optimize for either maximal or minimal target gene expression. To design for efficient prime editing, we limit the insertion site offset to +/- 2 bp from the original position, limit deletion of endogenous sequences flanking the insertion site to 4 bp, and limit the size of the edit (inserted + deleted base pairs) to 10 bp. Note that this restricted action space (randomly perturbing a small sequence within a larger sequence context) differs from most previous uses of simulated annealing for rational sequence design, which typically involved optimizing larger sequences (100-200 bp) of fixed length^76–80^.

We used this approach to design a total of 185 sequence edits (inserted sequence size range = 4-10 bp) at 5 high-efficiency pegRNA editing sites at the *PPIF* promoter (21-166 bp upstream of the TSS) (**Fig. 5a, Supplementary Table 3** & **Methods)**. For both THP-1 and Jurkat cells, we designed edits that were optimized to (i) maximally increase gene expression, (ii) maximally decrease gene expression, (iii) have no effect on gene expression, or (iv) maximize the fold-change in expression versus the other cell type (**Fig. 5a, Supplementary Table 9** & **Methods**).

**Figure 5.**
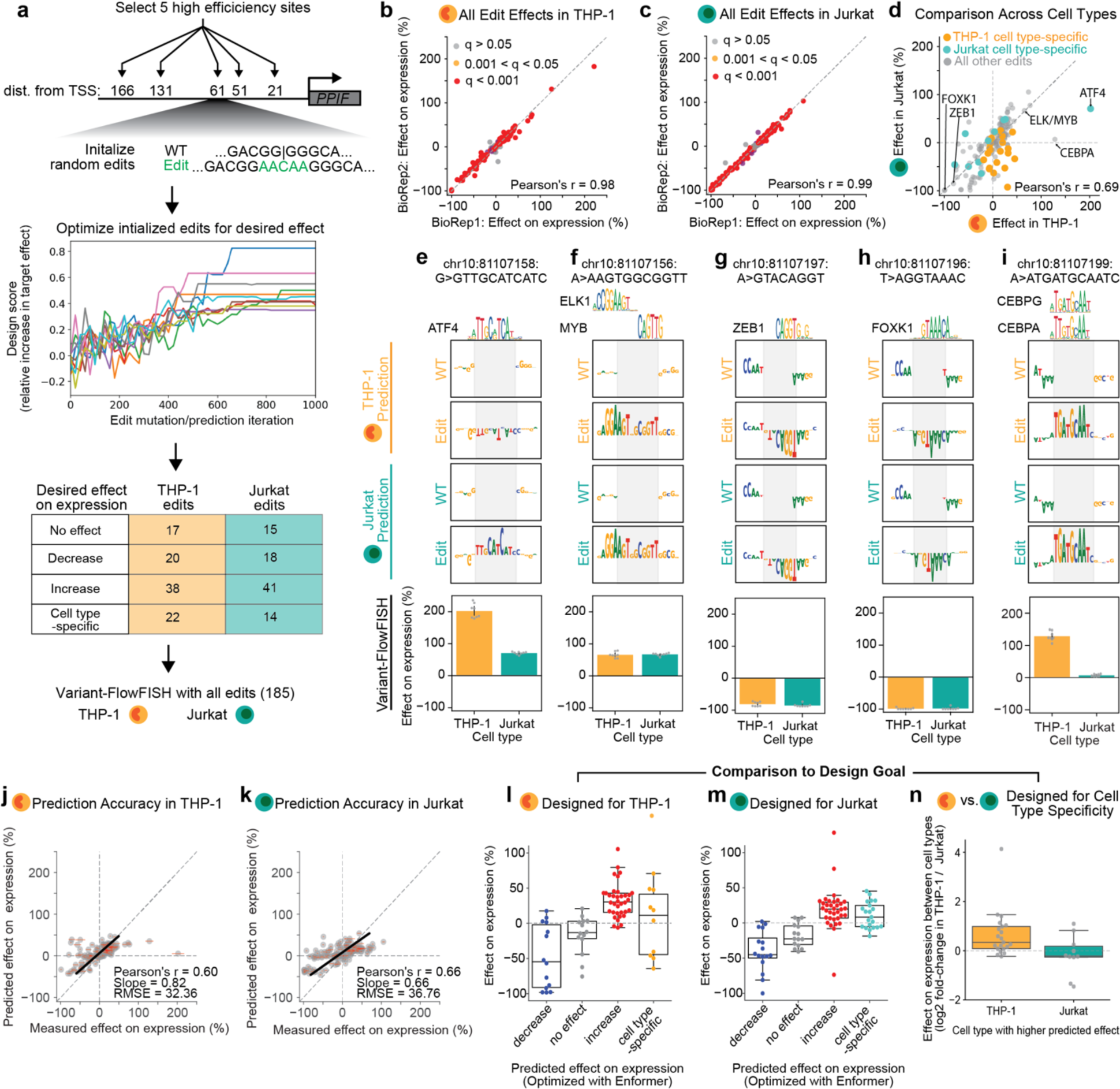
Designed sequence edits reprogram *PPIF* gene expression. (**a**) Overview of the design framework and Variant-FlowFISH screening of edits designed with Enformer. We selected 5 sites in the *PPIF* promoter that had high editing efficiency in the promoter tiling mutagenesis experiment and initialized random sequence edits ≤10 bp. We then used Simulated Annealing with a standard Metropolis acceptance criterion to optimize these sequences for specific effects on expression via 1,000 iterations of 1-bp sequence changes. We used the predicted difference between the wild-type and edited *PPIF* promoter CAGE signal from Enformer as the fitness predictor. We designed edits with different predicted outcomes on expression for both THP-1 and Jurkat cells, and combined these edits into a single pool of 185 pegRNAs to test in both cell types. (**b**) Variant-FlowFISH measurements of variant effects on *PPIF* expression (%) in THP-1 cells are highly correlated between two biological replicates. Dots (*n*=164): all variants passing the frequency threshold. Red: *q* < 0.001. Yellow: *q* < 0.05, gray: *q* > 0.05 (Benjamini-Hochberg corrected *p*-value, one-sample t-test). (**c**) Similar to **b**, but for Variant-FlowFISH measurements of variant effects on *PPIF* expression (%) in Jurkat cells. (**d**) Comparison of effects between THP-1 cells and Jurkat cells for all edits. Edits designed to increase expression specifically in THP-1 or Jurkat (cell type-specific designs) are colored orange and green, respectively. Selected edits that introduce predicted motif instances are annotated with the name of the corresponding transcription factor. (**e-i**) *In silico* mutagenesis from Enformer for select edits annotated in (d) was performed using THP-1 and Jurkat CAGE heads, revealing motif instances for the transcription factors (**e**) ATF4, (**f**) ELK/MYB, (**g**) ZEB1, (**h**) FOXK1, and (**i**) CEBPG/CEBPA predicted to be created by sequence edits. Edited sequences are highlighted in gray, including inserted base pairs (Edit) and deleted base pairs (WT). Transcription factor motif PWMs in e-i are from JASPAR^56^. Barplots (bottom) are the effects of each select edit in THP-1 and Jurkat cells. Each dot is a Variant-FlowFISH replicate and the error bar represents the 95% confidence interval of the mean. In **i**, Enformer interpretation appears to better match the motif for CEBPG, but CEBPG is similarly expressed between both cell types (THP-1 and Jurkat) and therefore is less likely than CEBPA (differentially expressed between cell types) to explain the differential effect of this edit on expression. (**j**) Comparison of measured effects on *PPIF* expression (Variant-FlowFISH) with Enformer predicted effects on gene expression (CAGE) in THP-1 cells. Dots: *n*=164 variants at the *PPIF* promoter with significant effects on expression in THP-1 cells. Error bars: 95% c.i. for measured effect size. Black line: Linear regression line of best fit. Legend lists Pearson’s r correlation coefficient, slope from the linear regression and room mean squared error (RMSE) of the predicted effects on expression (%). (**k**) Similar to **j**, for Jurkat cells. (**l**) Measured effects of edits designed for THP-1 cells to decrease expression (red), have no effect on expression (gray), increase expression (blue), or increase expression relative to Jurkat cells (orange). Boxplots show median, interquartile range, and whiskers show the rest of the distribution, except for points that are “outliers” from the interquartile range. (**m**) Measured effects of edits designed for Jurkat cells to decrease expression (red), have no effect on expression (gray), increase expression (blue), or increase expression relative to THP-1 cells (green). (**n**) Log2 fold-change of measured effects on *PPIF* expression between cell types (THP-1/Jurkat) for edits designed to increase expression in one cell type versus the other.

We introduced a library of pegRNAs encoding these 185 edits into each cell type, and performed Variant-FlowFISH. Of the 185 edits in the pool, 164 (88%) were observed at a frequency of greater than 0.01% in both cell types (range: 0.01-1.4% in THP-1 and 0.01-0.94% in Jurkat) (**Supplementary Figure 17**). The effect sizes of these 164 edits were highly correlated across replicates in both cell types (Pearson’s r = 0.99; **Fig. 5b,c, Supplementary Figure 18**). The effects of 144 and 150 edits in this screen had statistically significant effects in THP-1 and Jurkat cells, respectively (**Fig. 5b,c, Supplementary Figure 19, Supplementary Table 9**).

Overall, the measured effects of variants were correlated with the predicted effects by Enformer (Pearson’s *r =* 0.60 and 0.66 for THP-1 and Jurkat, respectively, **Fig. 5j,k**), and in total 84% of variants designed to increase or decrease gene expression significantly affected *PPIF* expression in the intended direction, although to varying degrees (**Fig. 5l,m**). In THP-1 cells, the groups of edits designed to increase, decrease, or have no effect on expression had average measured effects of +20%, -40%, and -18%, respectively **(Fig. 5l**). Similarly, in Jurkat cells, the groups of edits designed to increase, decrease, or have no effect on expression had average measured effects of +32%, -47%, and -16%, respectively (**Fig. 5m**). On average, edits designed to increase *PPIF* expression in THP-1 relative to Jurkat had a 1.6-fold greater effect in THP-1, and edits designed to increase expression in Jurkat relative to THP-1 had a 1.1 fold-change greater effect in Jurkat (**Fig. 5n**).

The range of effect sizes was striking: We identified edits that silenced *PPIF* expression (estimated -100% effect), as well as edits that increased *PPIF* expression by as much as +202% (**Fig. 5d**). For example, we inserted 10 bp at a position -60 bp relative to the TSS, and observed a +202% and +71% effect on gene expression in THP-1 and Jurkat cells, respectively (**Fig. 5e**). This edit was predicted to create a motif instance for the activating transcription factor ATF4, which is highly expressed in both of these cell types (TPM > 200). Another 10-bp insertion, located -62 bp from the TSS, was predicted to create tandem motif instances for ELK1 and MYB, which increased *PPIF* expression by approximately +65% in both cell types (**Fig. 5f**). Edits with strongest negative effects on expression appeared to create motif instances for known repressors. For example, an 8-bp insertion paired with a 1-bp deletion of endogenous sequence at -22 bp from the TSS created a motif instance for ZEB1, which decreased expression by more than -80% in both cell types (**Fig. 5g**). Similarly, an insertion of 8-bp paired with a 1-bp deletion of endogenous sequence at -23 bp from the TSS created a motif instance for FOXK1, which effectively silenced *PPIF* in both cell types (-99% and -98% effect in THP-1 and Jurkat cells, respectively; **Fig. 5h**). We also identified edits in our screen that had strong effects in only one cell type (**Fig. 5d,k**). For example, one 10-bp insertion increased expression 2-fold specifically in THP-1 cells (effects of +128% and +7% in THP-1 and Jurkat, respectively) (**Fig. 5d,i**). This edit was predicted to create a motif instance for CEBPA, which indeed was differentially expressed (TPM = 68 and 1 in THP-1 and Jurkat, respectively) and which was predicted to strongly affect chromatin accessibility in THP-1 but not Jurkat (TF-MoDISCo seqlets = 35,932 and 0 for THP-1 and Jurkat, respectively).

Altogether, these results demonstrate that *PPIF* expression is highly tunable with even small ≤10-bp changes its promoter sequence, and illustrate how Variant-FlowFISH can be combined with computational design strategies to rationally program gene expression with prime editing. The diverse ranges of effect sizes highlight how an iterative combination of both computational modeling and Variant-FlowFISH experiments will be required to identify edits with the desired effect.

## Discussion

Here we introduced Variant-FlowFISH to measure the quantitative effects of hundreds of designed DNA edits on gene expression, and applied it to map and reprogram gene regulatory sequences for a key gene involved in mitochondrial metabolism and immune disease: *PPIF*. Variant-FlowFISH achieves high power to detect subtle effects on gene expression, is highly reproducible, enables quantitative measurements of hundreds of variants in parallel, and is generally applicable to study effects across multiple cell types and states. We demonstrate how this technology will enable systematic mapping of endogenous regulatory sequences, learning the context-specific regulatory logic of transcription factor binding sites, advancing predictive models, and developing tools to reprogram gene expression.

Variant-FlowFISH will enable large-scale mutagenesis of human regulatory elements *in situ.* At the *PPIF* locus, our tiling mutagenesis experiments provide the most comprehensive view to date of the effects of regulatory sequences on gene expression in an endogenous context, and highlight 4 aspects of the architecture of regulatory elements in the human genome. First, functional nucleotides are sparse: combining results from both the enhancer and promoter, only 28% of tested 5-bp tiles appeared to correspond to endogenous regulatory sequences (**Fig. 2e**). Second, endogenous transcription factor binding sites had large effect sizes and appeared to act super-additively with respect to gene expression. Third, from tiling just two of the many regulatory elements of *PPIF*, we found 4 likely binding sites corresponding to transcription factors with reported effect on mitochondrial function, showing how the regulatory landscape of *PPIF* integrates diverse signaling inputs. Finally, our analysis comparing effects of variants with predictive models suggests that the motifs identified in the *PPIF* enhancer and promoter may involve factors that contribute to gene activation through different mechanisms, including via effects on transcriptional initiation (NRF1) or long-range 3D contacts (CTCF). Qualitatively, these observations are consistent with a long body of work to individually dissect regulatory elements in *Drosophila*, mouse, and other model organisms^46–54^; with mutagenesis experiments performed in plasmid-based reporters in human cells^15–22^; and with predictive maps derived via *in silico* interpretation or mutagenesis (^6,79,81^, *e.g.*, **Supplementary Fig. 7**). Variant-FlowFISH will enable accelerating such studies to dissect regulatory elements directly in the human genome.

Variant-FlowFISH will enable new types of studies to understand how transcription factor binding sites encode cell type-specific patterns of gene expression. Here, we compared the effects of inserting transcription factor binding sites into the *PPIF* promoter between three cell types and states, and expected that differences in effects would be determined by the cell type-specific activities of the corresponding transcription factors. Instead, the effects of variants were highly correlated and differed between cell types by a constant scaling factor (**Fig. 3g**), an effect that we validated for several individual edits (**Supplementary Fig. 14)**. This suggests that cellular context can affect the contributions of genomic variation and transcription factor binding sites to gene expression in multiple ways—not only via the cell type-specific activity of the cognate transcription factor, but also possibly via the cell type-specific responsiveness of the promoter. Further studies will be required to identify the mechanisms underlying such differences.

Because noncoding DNA sequences have highly context-specific effects and CRISPR variant editing will not be possible in many cell types *in vivo* in the human body, development of accurate computational models for predicting effects of variants on gene expression will be essential for complete dissection of human gene regulatory sequences. Previous work in other domains such as 3D protein structure prediction and enhancer-gene regulatory interactions has highlighted how an important step in the development of such models will be the collection of sufficiently large gold-standard datasets^82,83^. The dataset we collected here represents, to our knowledge, the largest describing the effects of isogenic sequence variants on quantitative gene expression in an endogenous genomic context, and so we explored benchmarking the performance of recent and new predictive models of variant effects. We considered Enformer^6^, ChromBPNet^42^, and ProCapNet^73^, which have vastly different input sequence contexts and model capacity, and are trained on different readouts of gene regulation. We find that model predictions were reasonably well correlated with Variant-FlowFISH data (up to a Pearson’s *r* = 0.65 for promoter variants), with notable differences between models and datasets (**Fig. 4**). First, although the long-range and local models used very different amounts of sequence context (165-Kb for Enformer, 2-Kb for ChromBPNet and ProCapNet), the local models performed as well if not slightly better at predicting effects on both expression and accessibility (**Fig. 4b, Supplementary Fig. 16**). Second, chromatin accessibility models (Enformer DNase-seq head, ChromBPNet) outperformed the corresponding expression models for both model architectures, similar to previous observations predicting effects in plasmid reporter assays^6^ (**Fig. 4b**). Third, none of the models appear to correctly interpret the effects of edits to the distal enhancers without explicit external calibration, either due to the limited sequence context of the local models (for ChromBPNet) or because the long-range models do not properly learn the importance of this distal sequence (for Enformer, **Fig. 4c**; for other benchmarks versus CRISPRi enhancer perturbation data, see also ^83,84^). Our results suggest an alternative route to capture long-range effects of variants, by first predicting the local effects of variants on enhancer accessibility or activity and then propagating those effects to gene expression via an enhancer-gene linking model. Finally, our CRISPR data also suggest that combining insights from models trained on different aspects of gene regulation, such as chromatin accessibility, transcriptional initiation, and long-range 3D contacts, may be required to capture the effects of noncoding variants that act through different mechanisms.

The ability to screen more complex, designed edits to regulatory DNA could enable new approaches to reprogram gene expression for treating disease. The recent approval of the first CRISPR therapeutic, targeting a cell type-specific gene regulatory sequence within *BCL11A* for sickle cell disease, suggests broad potential for genome editing approaches that tune gene expression^4^. Here we demonstrate the ability to use Variant-FlowFISH, together with predictive modeling, to reprogram gene expression through designed edits to a promoter. Small edits (4-10 bp) can turn *PPIF* expression either 3-fold up or nearly completely off, and engineer an inducible response to a stimulus. Because *PPIF* is known to regulate cell type-specific processes in disease, including inflammatory signaling in macrophages^2,85^ and reperfusion injury in the liver and heart^86,87^, future work will characterize how tuning the expression of *PPIF* could reprogram cell-type specific responses. Many other diseases may benefit from therapeutic approaches to quantitatively tune gene expression in a cell type-specific fashion, including genetic diseases of haploinsufficiency^88,89^ and complex diseases in which noncoding variants implicate specific regulatory elements in disease risk^5^. The series of experiments we present here—involving first mapping the endogenous regulatory motifs within regulatory elements, identifying efficient prime editing sites, and then iteratively applying computational modeling and Variant-FlowFISH— provides a generalizable strategy to develop genome editing reagents that create the desired change in gene expression.

We note several methodological considerations, limitations, and potential improvements to guide future applications and development:

i. Scale: We anticipate that Variant-FlowFISH will enable studying up to thousands of genomic variants at a time. Here, we successfully test up to 185 variants in a single pooled experiment, with scale limited by the editing efficiency of CRISPR prime editing (here, total 20-40% editing per pool), desired power for small effect sizes (*e.g.*, >80% power for >10% effects), and the throughput of RNA FlowFISH (∼20 million cells per experimental replicate). Scale could be adjusted with changes to any of these three variables.
ii. Completeness: We aimed to comprehensively mutagenize two regulatory elements, but in practice 13% of targeted 5-bp tiles did not show sufficient editing to assess their effects. This could be due to limitations in PAM positioning, pegRNA sequence composition, or steric hindrance from transcription factors bound to these positions (**Supplementary Fig. 5**)^33,39–41^. Our strategy would particularly benefit from advances in the efficiency and/or predictability of CRISPR prime editing^39,40,90^. Future adaptations should consider that, due the possibility of artificial correlations between variants in diploid cells, new CRISPR editors will need to be both efficient at introducing the intended edit as well as precise in avoiding unintended edits (**Supplementary Fig. 2**).
iii. Applicability to other genes and phenotypes: RNA FlowFISH is generally applicable to study many genes (*e.g.*, as in ^37,91^) without requiring special reagents such as antibodies or GFP-engineered cell lines. However, the same experimental design and Variant-EFFECTS analysis should be applicable to enable studying variant effects on other molecular or cellular phenotypes where suitable fluorescence sorting strategies are available.

In summary, our study provides a technology and systematic strategy for dissecting and reprogramming the sequences within regulatory elements that tune cell type-specific gene expression. We anticipate that this tool will be generally applicable to characterizing the effects of genomic variants associated with disease, learning principles of gene regulation, and developing genome editing therapies that rationally reprogram gene regulation.

## Methods

### Generating doxycycline-inducible prime editing (PE2) cell lines

We generated inducible PE2 prime editing cell line by transducing THP-1 and Jurkat cells with a construct expressing rTA linked to IRES to a neomycin resistance cassette expressed from an EF1ɑ promoter and selecting with G418. We generated an inducible PE2 lentiviral plasmid (TRE3G-PE2-P2A-BFP) by performing gibson assembly with the TRE3G promoter and P2A-BFP-WPRE amplified from TRE-KRAB-dCas9-IRES-BFP (addgene #85449) combined with nCas9(H840A)-MMLV(RT) amplified from pLenti-Synapsin-hChR2(H134R)-EYFP-WPRE (Addgene# 20945) and 7.3Kb (lentiviral backbone) and 5.6Kb (nCas9-partial RT without promoter) fragments digested (with XbaI, ClaI, AgeI) and purified from pLenti-Synapsin-hChR2(H134R)-EYFP-WPRE. We then transduced these rTA-expressing cells with TRE-KRAB-dCas9-IRES-BFP and selected for cells expressing BFP by FACS, yielding polyclonal PE2-BFP inducible cell lines.

### Tissue culture and stimulations

We maintained THP-1 and Jurkat cell (ATCC) density between 100K and 1M per mL in RMPI-1640 (Thermo Fisher Scientific) with 10% heat-inactivated FBS (Thermo Fisher Scientific), 4 mM L-glutamine, and 100 units/mL streptomycin and 100 mg/mL penicillin. We maintained HEK293T cells (ATCC) between 20-90% confluence in DMEM with 1 mM sodium pyruvate, 25 mM glucose (Thermo Fisher Scientific) and 10% heat-inactivated FBS. For stimulation of Jurkat cells, we added 5 ug/mL anti-human anti-CD3 (Biolegend-317315) and 100 ng/mL phorbol 12-myristate 13-acetate (PMA, Sigma Aldrich #P1585) for 4 hours at a cell density of 800K to 1M cells/mL then immediately proceeded with fixation (Variant-FlowFISH) or lysis (qPCR).

### pegRNA design

#### 1) For *PPIF* 5’ splice site edits and prime-editing optimization at the HEK3 target site

We designed small pools of prime-editing guide RNAs (pegRNAs) to target the first *PPIF* 5’ splice site, or the previously described HEK3 locus^33^, to optimize prime editing efficiency and demonstrate the application of the Variant-FlowFISH technology. Briefly, the gRNA spacer for the *PPIF* 5’ splice site was designed using the online tool PrimeDesign^92^ and we chose to use a previously validated gRNA spacer at the HEK3 locus^33^. We used the original scaffold RNA sequence^33^ (sequence shown in **Supplementary Fig. 20**), the reverse transcription template (RTT) length was between 14-17 bp long and the primer-binding site (PBS) length was between 11-13 bp long. The desired edits, ranging from single-nucleotide variants to substitutions 4 bp in length were introduced into the RT template. Immediately following the pegRNA was a transcription termination signal (string of 7 T nucleotides). Oligos were designed as two fragments which could be PCR amplified off each other. See **Supplementary Table 2** for more details on the design of these pegRNAs and further details are below as to how these individual pegRNAs were cloned.

#### 2) For Variant-FlowFISH screens

To design edits for mutagenesis of a target region, we randomly selected three 5-bp mutagenesis substitutions from a list of 12 pre-designed substitutions (AACCC, AGCCA, CAGTC, CCCTT, CGCCG, CTCAG, GAGGC, GCAGC, GGGAA, GTGCA, TCGGT, TTGGG) to be introduce at each 5-bp window tiling across the target region: the *PPIF* enhancer target region is located at chr10: 81,046,381-81,046,556 and the *PPIF* promoter target region is chr10: 81,107,026-81,107,246, relative to the TSS (as determined by RefSeq^93^). *PPIF* enhancer (chr10: 81,046,381-81,046,556) and *PPIF* promoter (chr10: 81,107,026-81,107,246) To design edits for insertion of transcription factor motif instances in the *PPIF* promoter, we identified factors that are (i) expressed in one or more tested conditions (THP-1, cells, Jurkat cells, or Jurkat cells stimulated with PMA+anti-CD3), (ii) have DNA binding motifs (from JASPAR and HOCOMOCO databases^94^ characterized by short, non-degenerate consensus sequences, (iii) and have been observed to affect gene expression in plasmid-based reporter assays^55,60,61^. To comply with the design of this initial 8 bp sequence insertion pilot screen, we designed insertions for specific transcription factors using the center 8 bp consensus sequence from the binding profiles. Design of edits using deep-learning is addressed later in the methods (see “Design and optimization of sequence edits with Enformer”). The pegRNA sequences for all of the Variant-FlowFISH screens are described in **Supplementary Table 3**.

To select the gRNA spacer component of each pegRNA, we generated a list of all possible NGG gRNA spacer sequences in the region and performed exhaustive evaluation of all potential off-target sites in the human genome (up to five mismatches) as previously described^95^. We selected the closest gRNA to the edit (maximum nick to edit distance of 50 bp) with a specificity score >50 that lacked homopolymer stretches of more than seven identical nucleotides in a row, and we included a leading “G” at the beginning of the selected gRNA if it was not already included. We designed pegRNAs to have a set PBS length of 11 bp and extended the RTT length 10 bp beyond the edit, with a maximum total reverse transcriptase template length of 50 bp (**Supplementary Fig. 4**). We filtered out pegRNAs with a RTT or PBS GC content of less than 30%, as well as pegRNAs containing homopolymer stretches of more than four “T” nucleotides in a row (encoded as “U” in the RNA) that could prematurely terminate Polymerase II transcription of the pegRNA. We used the optimized 86-bp flip + extension scaffold for all of our pegRNAs, which we and others have found to yield high editing efficiency (by 1.46-fold) relative to the standard scaffold (see **Supplementary Fig. 20**, which includes the optimized scaffold sequence).^39,90,96^

### Cloning pegRNA libraries or pegRNAs individually

For tiling mutagenesis experiments, we cloned pegRNA libraries with PCR tags (ordered from Agilent Technologies) into the lentiviral vector SgOpti (Addgene #85681). We first amplified the subpool of pegRNA oligonucleotides for each experiment using primers against the subpool-specific PCR tags. Next we performed a secondary PCR to add homology arms for Gibson assembly into SgOpti (**Supplementary Table 10**). If cloning pegRNAs individually, we start the cloning protocol by PCR amplifying the two oligo fragments (Fragment 1 and 2) and then performing this secondary PCR step. Each of these PCR steps was followed by a 1.5X Ampure XP SPRI purification. We prepared the SgOpti backbone by digesting with BsmBI, ClaI and EcoRI (New England Biolabs), followed by purification with 1X Ampure XP SPRI. Gibson assembly was performed for 1 hour at 50°C using Gibson Assembly Master Mix (New England Biolabs), 500 ng backbone, and 70 ng of purified PCR-amplified pegRNA pool in a final volume of 40 uL. We purified the gibson assembly with 0.7X Ampure XP SPRI and eluted in 15 uL, then electroporated 10 uL into Endura competent cells (Lucigen #602422) and then expanded the cells in liquid culture for 18 hours at 30°C and purified library plasmid with the Nucleobond Xtra Midi EF kit (Machery-Nagel #740420.50). To sequence-validate pegRNAs and ascertain their relative abundance for use in Variant-FlowFISH analysis, we PCR amplified and sequenced the pegRNAs from each purified plasmid library. The same protocol was used for cloning the pegRNA libraries for designed sequence edits, with the exception of first modifying the SgOpti vector to include the trimmed evopreQ1 (tevopreQ1) RNA pseudoknot motif^39^ 3’ to the BsmBI restriction enzyme site. This yielded assembled pegRNAs with the tevopreQ1 sequence encoded immediately 3’ to the PBS.

### Lentivirus production

We plated 550,000 HEK293T cells on 6-well plates and 24 hours later we transfected the cells with 900 ng psPAX2 packaging vector (Addgene #12260), 360 ng pMD2.g VSV-G envelope vector (Addgene #12259), and 1.2 ug of purified plasmid library using 5.8 uL of X-tremeGENE HP^TM^ DNA Transfection Reagent (Millipore Sigma #06366236001). We harvested viral supernatant 24 hours later with 0.45 uM filtration.

### Lentiviral infection, selection, and doxycycline-induced prime editing

We transduced THP-1 or Jurkat cells harboring doxycycline-inducible PE2 at a multiplicity of infection (MOI) of 2 at a coverage of ≥270,000 transduced cells per pegRNA. We transduce at such high coverage to account for the observed prime editing efficiency in our experiments, wherein ≤40% of cells are edited in each experiment. This equates to an equivalent “edited cell” coverage of approximately 70,000 cells or more per edit. Because not every pegRNA is efficient at introducing the desired edit, preliminary experiments showed that the percentage of total editing increased by increasing the MOI (0.2 vs. 2) and by also increasing the amount of time the PE2 machinery is induced by a doxycycline treatment (**Supplementary Figure 21**). Transduction was performed by adding lentivirus and polybrene (Millipore Sigma #TR-1003-G) to a final concentration 8 ug/mL in the cell suspension, which we then centrifuged at 1,200xg for 40 minutes at 37°C. We performed 72h of puromycin selection (1.5 ug/mL and 1.0 ug/mL for THP-1 and Jurkat respectively), after which we resuspended cells in fresh media containing 1.0 ug/mL doxycycline to induce prime editing and 0.3 ug/mL puromycin for maintenance selection. We maintained cells in this PE2-induction media for 13-21 days at their initial infection coverage, then removed doxycycline and continued to maintain cells for 7-10 days before proceeding to downstream assays to ensure that doxycycline-induced PE2 was degraded and absent from the cells.

### PrimeFlow RNA assay

We used the PrimeFlow RNA Assay Kit (#88-18005-210) according to the manufacturer’s instructions, with some modifications. Specifically, we used 12-15 million cells per reaction (12 reactions and 1 control reaction per biological replicate) and performed the fixation and permeabilization steps in one tube for all reactions from a single biological replicate (>156M cells for 13 reactions). We split reactions into individual 1.5 mL tubes prior to the target probe hybridization step, during which the control reaction was incubated with 100 uL of Target Probe Diluent without target probes. The control reaction is used to assess background fluorescence signal from non-specific binding of the label probes (see **Supplementary Fig. 22**). Target probes are as follows: gene of interest (*PPIF;* Thermo Fisher Scientific Type ‘Type 1’ probe set #VA1-30000735) and positive control housekeeping gene (*RPL13A;* Thermo Fisher Scientific ‘Type 4’ probe set *#*VA4-13187). Each sample (including the control/background staining reaction) was stained with the fluorescent Label Probes (Alexa Fluor 647 and 488 for *PPIF* and *RPL13A,* respectively) and we added a total of five washes with 40°C wash buffer following the final wash described in the manufacturer’s protocol to remove as much unbound label probes as possible before proceeding to cell sorting.

### Cell sorting

We diluted the samples to a concentration of approximately 3 x 10^7^ cells per mL in PBS with 0.5% BSA and filtered through a 35-um filter (Corning, no 352235) by centrifugation at 800g for 5 minutes at 4°C. We aliquoted approximately 2.5 x 10^5^ cells from the unstained samples for use as total FACS input controls. For tiling mutagenesis and designed sequence edit experiments, we sorted approximately 20 million cells for each FlowFISH replicate into 4 bins based on the fluorescence intensity of the target gene using the MA900 Multi-Application Cell Sorter (Sony). For splice site disruption and motif insertion experiments, we sorted up to 6 million cells for each FlowFISH replicate into 6 bins based on the fluorescence intensity of the target gene using the Influx Cell Sorter (BD Biosciences). To control for differences in staining efficiency of each cell, we normalized the fluorescence associated with the gene of interest (*PPIF*) to that of a housekeeping control gene (*RPL13A*). Specifically, we used color compensation to subtract a portion of each cell’s AF647 signal (*PPIF*) based on the intensity of its AF488 (*RPL13A*) signal. This portion is selected such that the mean of the AF488 signal in the top and bottom 25% of cells based on AF647 was approximately 10%. For the 4-bin sorting experiments, we set the gates for each bin on the compensation signal to capture 25% of the cells (full distribution sort). For the 6-bin sorting experiments, we set the gates for each bin on the compensation signal to capture 10% of the cells (60% distribution sort) according to the percentiles (1) 0-10%, (2) 10-20%, (3) 35-45%, (4) 55-65%, (5) 80-90% and (6) 90-100% (see **Supplementary Figure 22** for an example of the gating and sorting strategy). We recommend using the four-bins sorting strategy to capture the entire distribution of cells and to maximize cell recovery from sorting.

### Genomic DNA extraction and amplicon sequencing

We collected the sorted cells by centrifugation at 800g for 5 minutes and 4°C, resuspended the pellet in 70 uL of lysis buffer per million cells (50 mM Tris-HCl, p 8.1, 10mM EDTA, 1% SDS) and incubated at 65° for 10 min for reverse cross-linking before cooling to 37°C. Once cooled, we added 2 uL of RNase Cocktail (Invitrogen, no AM2286), mixed well and incubated at 37°C for 30 min. Finally, we added 10 uL Proteinase K Solution (Invitrogen, no 4333793), mixed well and incubated at 37°C for 2 h followed by incubation at 95°C for 20 min. We purified genomic DNA using a 0.7X clean-up with Agencourt XP (SPRI) beads (Beckman Coulter).

Following extraction, we split clean genomic DNA into PCR replicates (each containing gDNA from approximately 250,000-500,000 cells) and amplified the genomic region of interest with Q5 High-Fidelity 2X Master Mix (New England Biolabs, no M0492L) and 0.5 uM of each amplification primer (**Supplementary Table 11**). We carried out genomic amplicon PCR with the following thermocycling parameters: 98°C for 1 min and then 28 cycles of 98°C for 10 s, annealing for 20 s (variable), 72°C for 20 s, followed by a final 72°C extension for 2 min. We purified PCR products using a 1.0X SPRI bead clean-up and used 3 uL of the clean product as input for a secondary sample indexing PCR. Specifically, each 10 uL reaction contained 3 uL of sample, 5 uL of Q5 High-Fidelity 2X Master Mix, and 0.5 uM of each indexing primer containing Illumina adapters: CAAGCAGAAGACGGCATACGAGATNNNNNNNNGTGACTGGAGTTCAGACGTGTGCTCTTC CGATCT; AATGATACGGCGACCACCGAGATCTACACNNNNTTTCCCTACACGACGCTCTTCCGATCT.

We carried out indexing PCR with the following thermocycling parameters: 98°C for 1 min and then 10 cycles of 98°C for 10 s, 60°C for 20 s, 72°C for 20 s, followed by a final 72°C extension for 2 min. We pooled equal volumes of indexing PCR product together and performed two consecutive 1.0X SPRI bead clean-ups, then measured DNA concentration using the Qubit 1X dsDNA HS Assay Kit (Invitrogen, no Q33231). We sequenced DNA amplicon libraries on an Illumina Nextseq 550 System using a custom Illumina Index 2 sequencing primer: TCGGAAGAGCGTCGTGTAGGGAAA.

### Analysis of Variant-FlowFISH screens (Variant-EFFECTS)

To determine the effects of each edit on fluorescence, we developed an analysis framework that takes as input sequencing reads for each edit across gene expression bins as well as other experimental metadata and infers quantitative effects of each edit versus the reference allele (**Supplementary Fig. 1**).

First, we demultiplexed raw sequencing reads using the bcl2fastq program (Illumina) and used CRISPResso2^97^ to align reads with a minimum average quality score of 20 (phred 33) to a reference amplicon sequence (CRISPResso -r1 {input.read1} -r2 {input.read2} -o results/crispresso/ --amplicon_seq {params.amplicon_seq} --amplicon_name {params.amplicon_id} --name {wildcards.SampleID} --quantification_window_coordinates 27-201 --exclude_bp_from_left 0 --exclude_bp_from_right 0 --plot_window_size 0 -- min_frequency_alleles_around_cut_to_plot 0.05 -q 20 -s 0). For the filtered and now aligned reads in each sample, we now need to count the number of unedited allele reads as well as the number of reads containing each edit.

This operation is performed by searching for each variant using a file that contains a “Mapping Sequence” column: a sequence containing the expected variant as well as 3-5-bp buffer on each side of the variant. If a read exactly contains a variant’s mapping sequence, we add it to the variant’s read count. The length of buffer sequence was determined to be an adequate number of bases to resolve any combination of variant and position within the amplicon (10 bp plus the length of the introduced edit for all experiments except for rationally designed edits, which use a fixed mapping sequence length of 16 bp).

Quantifying the unedited allele reads necessitates an alternative approach due to the fact that the entire amplicon now must be assessed for edits. An important consideration here is the increased likelihood of observing PCR or sequencing error when considering the entire amplicon. To account for this, we use the corresponding unedited cell line and estimate the background average number of sequencing or PCR errors for a given amplicon. We use this number as a threshold for the maximum number of errors a read can contain when compared to the reference sequence and still be counted as an unedited read. We discard the reads that fall below this threshold and don’t contain a designed variant.

We then aggregate the read counts across PCR replicates to obtain a total count for unedited reference and designed edit sequences in each FACS bin. We normalized these aggregated counts so that the sum of reference and variant alleles in each sample equates to the total number of cells sorted into that FACS bin (*i.e.*, to account for differences between the number of sequencing reads and the number of cells in a given bin).

We then used the limited-memory Broyden–Fletcher–Goldfarb–Shanno algorithm maximum-likelihood method in the R stats4 package to fit the read counts in each fluorescence bin to the log-normal distribution that has the highest likelihood for having produced the observed counts in the bins. To calculate effect sizes, we determined the percent difference between the mean of the log-normal fit for each edit and the mean of the log-normal fit for the wild type reference sequence.

We applied two scaling factors to these effect sizes: one to account for background signal in the RNA FISH assay and one to account for heterozygous editing events (genotype frequencies) during prime editing.

The first adjustment is needed to account for background signal in the RNA FlowFISH assay, which we assume combines additively with real signal due from the target gene of interest. To correct for this background, we used the difference between CRISPRi-FlowFISH TSS guide knockdown and qPCR effect size to generate a linear scaling factor applied to our effect sizes (1.48 and 1.33 for THP-1 and Jurkat, respectively), which accounts for FlowFISH signal from non-specific RNA FISH probe binding (this is based on our observation that promoter CRISPRi typically shows 80-90% knockdown by qPCR).

The second adjustment accounts for the frequencies of genotypes in the cell population. This is needed because, for a given desired edit, there will be some distribution of allelic editing events. Some fraction of our cells will be edited in a heterozygous fashion where only 1 allele contains the edit, leading to an underestimation of the effect of a given variant in those specific cells. With this in mind, we estimate the frequencies of each genotype for each edit and use this to correct the estimated effect sizes.

To estimate genotype frequencies, we first calculate the editing efficiency (*p*) of each designed edit by dividing the observed frequency in the population (FACS input population) by the frequency of the corresponding pegRNA in the plasmid library (from sequence-verification of plasmid constructs). For plasmid libraries that were generated by arrayed pegRNA cloning and equimolar plasmid pooling (PPIF 5’ splice site and 8-mer insertion experiments), we assumed that pegRNAs were present at equal frequencies in the pool. Then, using this estimated editing efficiency (*p*), we compute the frequency of each genotype assuming that the two chromosomes in a diploid cell are edited independently: homozygous edited (frequency = *p*^2^), heterozygous edited (frequency = 2*p*(1-*p*), and homozygous wild-type (frequency = (1*-p*)^2^). Expression of the target gene in cells of each genotype are related to the sum of effects on each of their two chromosomes, such that the expression level of a cell is 1+*x* for homozygous edited cells, 1+(*x*/2) for heterozygous edited cells, and 1 for homozygous wild-type cells, where *x* is the true effect size of the edit (*x*). We assume that expression levels for all three genotypes are normally distributed and have the same variance in log space during the FACS sort.

With these assumptions, consider a simplified case in which a screen contains only 1 pegRNA. In this case, the (uncorrected) effect inferred from the MLE procedure for the edited allele (*y*) will be related to the true effect size (*x*; positive *x* indicates that edit increases gene expression) and editing efficiency (*p*) by the following equation:

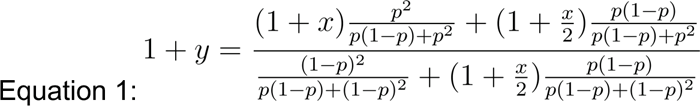

The equation states that the uncorrected estimate of the effect of a variant from Variant-FlowFISH (left) is equal to the expression in homozygous edited cells (1+x) multiplied by the the proportion of the cells that are homozygous edited plus the expression in heterozygous cells (1+*x*/2) multiplied by half of the proportion of cells that are heterozygous edited, divided by the expression of homozygous wild-type cells (1) multiplied by the proportion of cells that are wild type (unedited) plus the expression of heterozygous cells multiplied by the other half of the heterozygously edited cells (right). This is essentially a weighted sum of expression levels of cells carrying a given allele (edit or wild-type).

We then solve for *x* (the change in effect size from reference), and obtain:

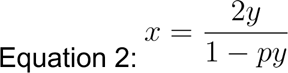

This equation specifies the correction needed in Variant-FlowFISH experiments involving a single edit (*e.g.*, individual edits in **Fig. 1e**).

Now, consider the case of a pool with multiple pegRNAs introducing different edits. In this case, the naive estimate of the expression level of the reference allele (denominator of Equation 1) depends not only on the effect size (*x*) of a single edit, but of all edits in the pool of cells (*e.g.*, this is graphically depicted in **Supplementary Fig. 2** part 8). In a pooled experiment where many cells are unedited (due to low editing efficiency), and/or where the average effects of edits in the pool is close to zero (*e.g.*, similar numbers of edits that increase or decrease expression), then the denominator of Equation 1 is approximately 1. Solving this simplified equation for *x*, we obtain:

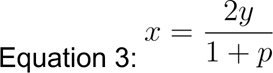

As such, in pooled experiments, we adjust each uncorrected effect *y* for its editing efficiency *p* using Equation 3. Equation 3 also provides some intuition that if there is almost no editing (p∼=0), the actual homozygous effect should be two times the measured effect. If almost all the alleles are edited (p∼=1), the homozygous effect should be very close to the measured effect.

### Setting variant frequency filters by assessing reproducibility as a function of frequency

To determine the relationship between edit frequency and effect size, we assessed the reproducibility of variant frequencies across PCR replicates from all FACS bins in an experiment. We found that the variant frequencies were highly reproducible across PCR replicates for variants above a frequency threshold of 0.0001, but that their correlations decreased substantially below this threshold (**Supplementary Figure 6**). PCR and sequencing error result in false positives in variant read counts, introducing noise into the estimations of effects. The variants we introduce contain at least 3 nucleotides distinguishable from the reference sequence which enables enhanced resolvability of variants from technical error and tracks with our capacity to assess variants at frequencies as low as 0.0001. Thus, we filter out variants that have an average bin frequency below 0.0001 (0.01% of sequencing reads) prior to effect size calculation for variants in a given FlowFISH sample (**Supplementary Figure 5**), and we only report effect sizes for variants in our mutagenesis screens that have at least 6 FlowFISH replicate measurements (at least two per biological replicate) that passed this additional filter.

### Single edit induction and clonal cell line selection

To generate cell lines for qPCR and ChIP experiments, we performed single-pegRNA lentiviral transduction and doxycycline-induced PE2 expression to produce polyclonal populations of cells with possible genotypes: homozygous WT, heterozygous WT/edit, edit/edit. To generate monoclonal cell lines that were homozygous for either the reference allele or the target edit, we sorted single cells from these populations into 96-well plates using FACS and expanded them. Genomic DNA was extracted from 250,000 cells for each clonal cell line in 25 uL QuickExtract^TM^ DNA Extraction Solution (BioSearch Technologies #QE09050) by thermocycling at 65°C for 6 minutes followed by 98°C for 2 minutes. The target locus was amplified and sequence as described above (*Genomic DNA extraction and amplicon sequencing)* to screen for monoclonal cell lines that were wild type or homozygous for the target edit.

### RNA extraction and quantitative RT-PCR

Total RNA was extracted from homozygous clonal cell lines with RNeasy Mini Kit (Qiagen # 74106) with DNase treatment according to the manufacturer’s protocol. Approximately 100 ng total RNA and 250 nM primer for *PPIF* (Fwd: AGAACTTCAGAGCCCTGTGC, Rev: CATTGTGGTTGGTGAAGTCG) or *RPL13A* (Fwd: CTCAAGGTGTTTGACGGCATCC, Rev: TACTTCCAGCCAACCTCGTGAG) were used to perform Quantitative RT-PCR using the Brilliant II SYBR Green QRT-PCR Master Mix (Agilent # 600825) with the following program: 45°C for 45 minutes for cDNA generation, 95°C for 10 minutes, followed by 45 cycles of 95°C for 30 seconds, 58°C for 45 seconds, and 72°C for 45 seconds for fluorescence detection. Effect on *PPIF* expression for each clonal edited cell line was calculated as the percent change of housekeeping-normalized CT from the geometric mean of the housekeeping-normalized CT for wild type clones (2^-(edited clone PPIF–RPL13A)- (geometric mean of PPIF–RPL13A from WT clones)^).

### Chromatin immunoprecipitation and sequencing

ChIP was performed on polyclonal single-edit THP-1 cell cultures as previously described^14^. Briefly, 1×10^8^ cells were cross-linked with 1% formaldehyde for 10 minutes at 25°C and quenched with glycine at a final concentration of 125 mM. Cross-linked cells were lysed and sonicated for 20 minutes (cycles of 30 seconds on/off) at 4°C to generate 300-600 bp chromatin fragments. An aliquot of total fragmented chromatin was saved as an input control and the remaining chromatin was split into two tubes (representing 5×10^7^ cells each) for antibody pulldown and incubated overnight at 4°C with 15 ug of NRF1 (Abcam #ab55744), or MYC (Abcam #ab56) antibody. Chromatin cross-linking was then reversed and DNA was eluted at 65°C overnight, purified and eluted in 40 uL of molecular biology grade water.

To generate *PPIF* promoter amplicon sequencing libraries from fragmented chromatin, we optimized PCR primers to amplify 114-bp of the *PPIF* promoter (Fwd primer target: AGGGGTAGTCCACGGACAG, Rev primer target: GCCTTTATTGGCTTGCCCAG) containing the edit site and performed PCR with the following thermocycling parameters: 98°C for 1 min and then 30 cycles of 98°C for 10 seconds, 68°C for 20 seconds, and 72°C for 20 s, followed by a final 72°C extension for 2 min. We purified PCR products using a 1.8X SPRI bead clean-up, and followed the protocol above for the remaining steps of the library preparation and sequencing (see *Genomic DNA extraction and amplicon sequencing)*.

For quantification of edited and wild type alleles in antibody-immunoprecipitated chromatin and total chromatin control (whole cell extract), we demultiplexed raw sequencing reads using the bcl2fastq program (Illumina) and filtered for sequencing reads that mapped unambiguously to either the wild-type or edited *PPIF* promoter sequences by running CRISPResso2 in HDR mode^97^. We used these reads to calculate allele-specific fold-change for transcription factor binding between edited and unedited alleles. Allele-specific fold-change = [(% of reads with edit in immunoprecipitation % of reads with insertion in whole cell genomic DNA extract) (% of unedited wild type reads in immunoprecipitation % unedited wild type reads in whole cell genomic DNA extract)].

### Enformer predictions and sequence attributions

The Enformer model^6^ was downloaded from ‘https://tfhub.dev/deepmind/enformer/1’. Effects of Variant-FlowFISH edits were predicted using two different methods: In the first ‘variant-centered’ approach, the 196 kb input window was centered on the site of the insertion or substitution. For 8-bp insertion edits specifically, the variant sequence was constructed by keeping the left-to-centermost portion of the input (98 kb) fixed and inserting the variant at the center, which pushed the center-to-rightmost endogenous sequence 8-bp to the right (left-justified, rightmost 8-bp trimmed off variant sequence to maintain prediction window size). Substitution variants were constructed by replacing the reference sequence with the corresponding substitution sequence at the same centered position.

In the second, ‘TSS-centered’ approach, the reference and variant sequences were constructed identical to the method described above, but the final sequences were re-centered such that the input window was centered on the annotated TSS of the *PPIF* gene instead of on the variant. This prediction configuration was used for predicting effects of *PPIF* enhancer edits on the distal *PPIF* promoter. For both of these approaches, each variant was scored by predicting the output coverage within a subset of cell type-matched assay tracks for both the reference and variant sequences (FANTOM IDs: CNhs10724 and CNhs11253 for THP-1 and Jurkat CAGE tracks, respectively. ENCODE IDs ENCFF659BVQ and ENCFF622JVD for monocyte (representing THP-1) and T cell (representing Jurkat) DNase tracks, respectively, aggregating the total signal within a local window centered on the variant or TSS and computing the change between the aggregated variant and reference sequence values. The local window size was chosen to be 768 bp, or 6 aggregated 128-bp Enformer output bins. The predicted effects were averaged across six additional shifts (the input sequence is shifted by a small offset and re-scored by the model), and both the forward and reverse-complement were scored per shift (14x total redundancy).

Sequence attributions were generated using in-silico saturation mutagenesis (ISM). Briefly, the reference nucleotide at every sequence position within a small window around the target edit site were systematically exchanged for every other possible nucleotide, and the log fold-change in predicted signal within the 768 bp output window was recorded for each substitution. The final ISM score is visualized per position as the average effect size across the 3 variant alleles. Only reverse-complement ensembling with no additional shifts was used in order to speed up computation.

### Design and optimization of sequence edits with Enformer

Using the CAGE prediction tracks for THP-1 and Jurkat cell types from the Enformer sequence-based deep learning model (see above), we applied simulated annealing with a standard Metropolis acceptance criterion^76,77^ to optimize short sequence edits at five high efficiency prime editing loci (chr10:81107053, 81107088,, 81107158, 81107168, 81107198) in the *PPIF* promoter. In brief, starting from a randomly initialized sequence insertion (≤ 10bp) at the targetable loci, we performed 1,000 randomly sampled alterations to the sequence edit (either of: (a) inserting one new nucleotide, (b) deleting one nucleotide, (c) shifting the insertion site by one position, or (d) substituting one existing nucleotide for another letter). The total edited sequence was restricted from growing beyond 10 bp. We always accepted edit alterations that increased the objective function we sought to optimize. Importantly, with a probability that decreases smoothly and monotonically as a function of the number of alterations lapsed (until the fixed budget of 1,000 alterations has been made), we accept ‘bad’ alterations that decrease the objective function value. By accepting bad changes with a small probability, we allow the optimization to escape local minima. After ∼700 iterations, this probability is near-zero, and the design method effectively becomes a standard hill-climbing algorithm that greedily accepts only good changes. By annealing slowly, we find short sequence edits that reach near-global optimal impact on promoter function as predicted by Enformer. In the application described here, we optimized the sequence edits for maximal CAGE fold change between the edited and the original PPIF promoter sequence (estimated from the three Enformer output bins overlapping the TSS), for THP-1 and Jurkat cell types, respectively. We also optimized edits to maximize the absolute fold change between the THP-1 and Jurkat effects (e.g., THP-1- or Jurkat-specific designs).

### ChromBPNet predictions, DeepSHAP sequence attributions and TF-MoDISco analysis

We downloaded ATAC-seq datasets for THP-1 cells (GSM4706091, GSM4706092), Jurkat cells (GSM4706084,GSM4706085) and stimulated Jurkat cells (GSM4706088,GSM4706089) from GEO. We processed the ATAC-seq datasets using the ENCODE ATAC-seq pipeline (https://github.com/ENCODE-DCC/atac-seq-pipeline), which aligns reads to the GRCh38/hg38 Reference Genome using the Bowtie2 aligner and identifies peaks using the MACS2 peak caller. We used the tagalign file pooled across replicates and the optimal overlap peak sets output from the ENCODE ATAC-seq pipeline to train individual, single-task ChromBPNet models for THP-1, Jurkat, and Jurkat under stimulation. The training was conducted using the ChromBPNet^42^ repository (https://github.com/kundajelab/chrombpnet/tree/master, v0.1.3). We sampled non-peak regions with GC content matching that of the peak regions as negative samples for training. All models were trained using reverse complement augmentation and five-fold cross-validation (CV).

ChromBPNet only uses the “variant-centered” approach for predicting the effects of Variant-FlowFISH edits. The 2114-bp input window was centered on the site of the insertion or substitution as described for Enformer above. Sequence contribution scores for ChromBPNet were generated using the DeepSHAP/DeepLIFT algorithm^44,45^. DeepLIFT generates base pair resolution sequence contributions by back propagating a score, which is derived from comparing the activations of all neurons for the input sequence with those from reference sequences. We created 20 dinucleotide-shuffled versions of the input sequences as reference sequences and then utilized the DeepSHAP implementation of DeepLIFT (https://github.com/kundajelab/shap) to generate sequence contribution scores. The predicted effects for Variant-FlowFISH edits and sequence contributions were averaged across five folds to obtain consolidated scores.

We then used TF-MoDISco^58^ to distill sequence motifs from the DeepSHAP contribution score profiles (averaged over 5 folds) over all peak region sequences in each cell type. TF-MoDISco is a motif-discovery method that identifies high importance regions from sequence contribution scores as seqlets and then aligns and clusters those seqlets into distinct, consolidated motif patterns. We computed sequence contribution scores centered around the ATAC-seq peaks for each cell type using its corresponding ChromBPNet model. We then applied TF-MoDISco, setting the parameters to 1 million seqlets and a window size of 500 bp to extract motifs from the sequence contribution scores for each cell type. For each motif identified by TF-MoDISco, we reported the top three matches to existing motifs using TOMTOM from the MEME Suite^57^. To compare the motifs learned between Jurkat and THP-1, we extracted seqlets from their respective MoDISco objects and reclustered them using *SimilarPatternsCollapser* function from TF-MoDISco. We averaged the contribution scores from the seqlets to construct the contribution weight matrix (CWM) for each motif pattern. The contribution score for each motif pattern was calculated by taking the maximum absolute value per position of each CWM.

### ProCapNet predictions and DeepSHAP sequence attributions

ProCapNet is a deep learning model that maps DNA sequence to base-resolution nascent transcription initiation profiles from PRO-cap experiments. The ProCapNet architecture matches that of BPNet, previously described in Avsec et al. 2021, with the following modifications. First, no control track was used; second, ProCapNet’s sequence input is 2114 bp and the output is 1kb wide; third, ProCapNet is adapted to account for the strand asymmetry of transcription: the counts tasks predicts the log of the number of PRO-cap reads summed over both strands, and the profile task applies the final softmax function across the predictions for both strands as one array; fourth, ProCapNet implements a mappability-aware training scheme, masking bases are not uniquely mappable by PRO-cap-length sequencing reads (according to 36-mer hg38 UMAP tracks from Karimzadeh et al. 2018)^98^ from being incorporated into the profile task loss function; and finally, all post-training ProCapNet predictions are the average of the model output for the forward and reverse strand sequences.

ProCapNet was trained on ENCODE processed K562 PRO-cap data (ENCSR261KBX filtered BAM files and peaks). TSS events were identified as R2 5’ ends. During training, K562 DNase hypersensitive sites (DHSs; ENCSR000EKS) not overlapping PRO-cap peaks were randomly sampled as negative examples, using a 1:7 ratio of DHSs to peaks within batches. Training was done using random 5’-3’ jittering up to 200 bp, reverse complement augmentation, and seven-fold cross-validation (CV) split by chromosome. The code repository is available at https://github.com/kundajelab/nascent_RNA_models, and further details will be available in the manuscript^73^.

Variant effect prediction and DeepSHAP sequence interpretation were performed with ProCapNet using the same strategy as ChromBPNet, with the PyTorch Captum implementation of DeepSHAP.

### Data Availability

DNA sequencing data for Variant-FlowFISH experiments can be found in the Impact of Genomic Variation on Function (IGVF) Data Portal (https://data.igvf.org) under accession numbers: IGVFDS5031MNRR and IGVFDS3423PSID for *PPIF* enhancer and promoter tiling experiments, respectively; IGVFDS5056OAGR, IGVFDS9162RHFP, and IGVFDS1863TJOE for *PPIF* promoter 8-bp insertion experiments in THP-1, Jurkat, and Jurkat cells stimulated with PMA + anti-CD3 antibody, respectively; IGVFDS4359OODY and IGVFDS6299IJHG for *PPIF* promoter rationally (Enformer) designed edits in THP-1 and Jurkat cells, respectively.

### Code and Model Availability

Variant-FlowFISH data processing and analysis: https://github.com/EngreitzLab/variant-flowfish

ChromBPNet code repository: https://github.com/kundajelab/chrombpnet

ProCapNet code respository: https://github.com/kundajelab/nascent_RNA_models

Enformer code repository: https://github.com/google-deepmind/deepmind-research/tree/master/enformer

ChromBPNet and ProCapNet models trained for this work: 10.5281/zenodo.10403551

Code for computational approach to rational design with simulated annealing and Enformer: 10.5281/zenodo.10403551

## Acknowledgements

We thank Ricardo Zermeno for technical support with flow sorting, and Ian Whaling, Meenakshi Kagda, and Idan Gabdank for assistance sharing the data via the IGVF Data Portal. We thank Greg Newby, Jonathan Levy, Andrew Anzalone, Luke Koblan, David Liu, Nir Hacohen, Michael Kane, Mark Osborn, John Ray, Carl de Boer, Michael Bassik, Gaelen Hess, and Ioannis Karakikes for reagents and technical discussions. This work was supported by the NIH-NHGRI Impact of Genomic Variation on Function Consortium (UM1HG011972 to J.M.E. and U01HG012069 to A.K.); an NIH-NHGRI Pathway to Independence Award (K99HG009917 and R00HG009917 to J.M.E.); the Novo Nordisk Foundation (NNF21SA0072102 to J.M.E.); Gordon and Betty Moore and the BASE Research Initiative at the Lucile Packard Children’s Hospital at Stanford University (J.M.E.); the Broad Institute (E.S.L.); The Walter V. and Idun Berry Postdoctoral Fellowship Program, Stanford University (G.E.M); and a Stanford Graduate Fellowship (Z.C.). This material is based upon work supported by the National Science Foundation Graduate Research Fellowship Program under Grant No. DGE-1656518 (to B.R.D and M.T.M). Any opinions, findings, and conclusions or recommendations expressed in this material are those of the authors and do not necessarily reflect the views of the National Science Foundation.

## Author Contributions

G.E.M. and M.T.M. contributed equally as co-first authors, and have agreed that either author can be listed first in reporting this study. G.E.M., M.T.M., and J.M.E. co-led the design, implementation, and writing of the study. G.E.M., M.T.M., H.J., K.G., and J.M.E. developed, optimized, and validated Variant-FlowFISH with CRISPR prime editing. B.R.D., H.J., G.M., C.P.F., E.S.L., and J.M.E. contributed to developing experimental and analysis protocols for FlowFISH. B.R.D., K.G., K.A.L., and J.M.E. developed the mathematical approach to correcting for artificial correlations in diploid cells. K.G., H.J., B.R.D., and J.M.E. developed and implemented the computational pipeline to estimate variant effects with maximum likelihood estimation. G.M. and J.M.E. engineered stable prime editing cell lines. G.E.M. led execution and analysis of *PPIF* 5’ splice site and tiling mutagenesis experiments. H.J., M.T.M., K.G. and J.M.E. contributed to tiling mutagenesis experiments and analysis. M.T.M. led design, execution, and analysis of promoter insertion and rationally designed edit experiments. K.G., G.E.M., and J.M.E. contributed to promoter insertion experiments and analysis. J.L. and J.M.E contributed to design and analysis of Enformer rationally designed edit experiments. M.T.M. led analysis of benchmarking computational models using Variant-FlowFISH data. Z.C., A.P., and A.K. conducted analysis with the ChromBPNet model. K.C. and A.K. conducted analysis with the ProCapNet model. J.L. and D.R.K. contributed analysis with the Enformer model. J.L. developed the computational method to optimize sequence edits for desired gene expression outcomes. G.E.M., M.T.M., and J.M.E. wrote the manuscript with input from all authors. J.M.E., A.K., E.S.L., and D.R.K. supervised aspects of the study.

## Supplementary Tables

**Supplementary Table 1.**
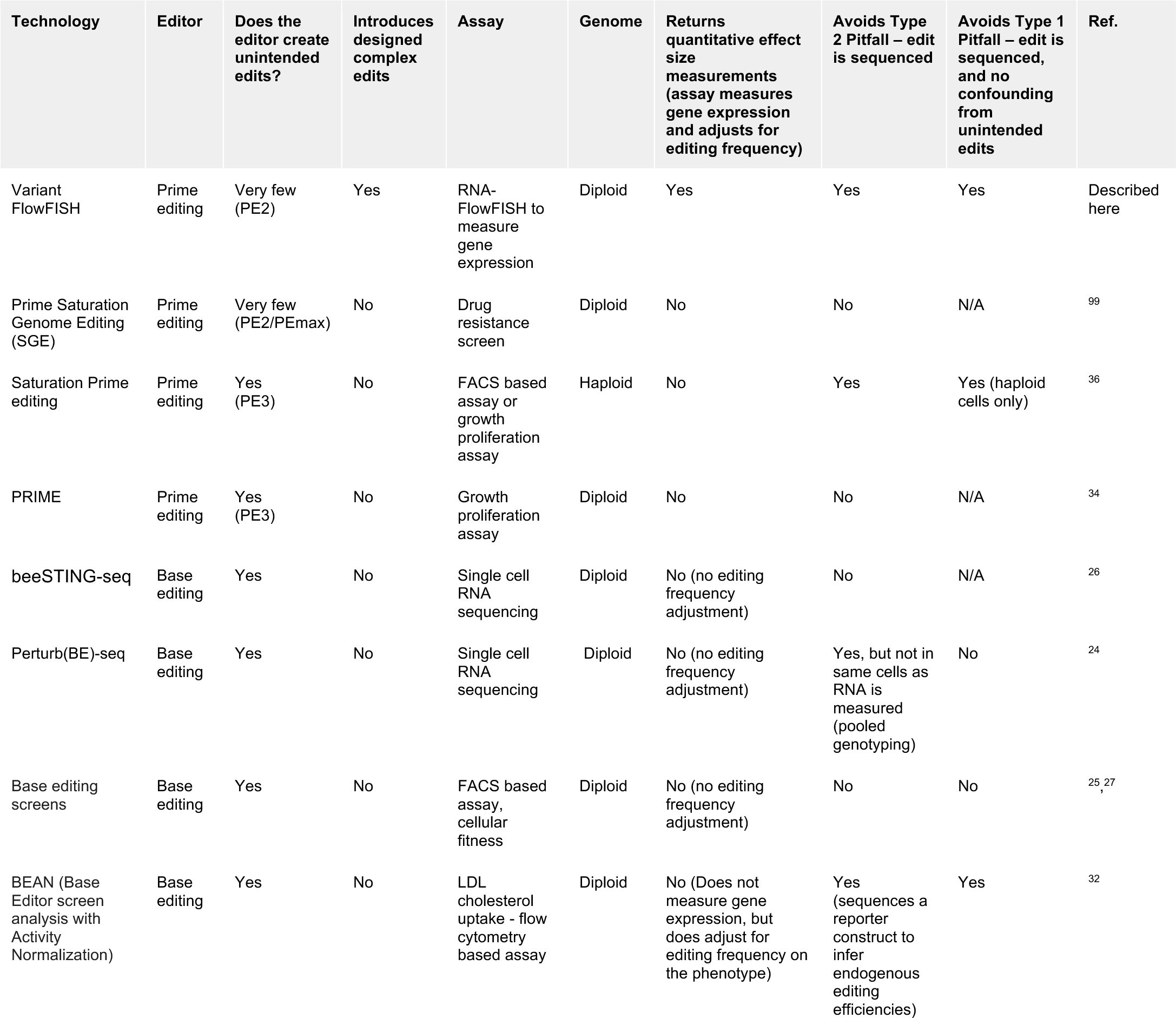
Comparison of Variant-FlowFISH to previous screening methods for measuring the effects of sequence edits on gene expression or other phenotypes.

| Technology | Editor | Does the editor create unintended edits? | Introduces designed complex edits | Assay | Genome | Returns quantitative effect size measurements (assay measures gene expression and adjusts for editing frequency) | Avoids Type 2 Pitfall – edit is sequenced | Avoids Type 1 Pitfall – edit is sequenced, and no confounding from unintended edits | Ref. |
| --- | --- | --- | --- | --- | --- | --- | --- | --- | --- |
| Variant FlowFISH | Prime editing | Very few (PE2) | Yes | RNA-FlowFISH to measure gene expression | Diploid | Yes | Yes | Yes | Described here |
| Prime Saturation Genome Editing (SGE) | Prime editing | Very few (PE2/PEmax) | No | Drug resistance screen | Diploid | No | No | N/A | 99 |
| Saturation Prime editing | Prime editing | Yes (PE3) | No | FACS based assay or growth proliferation assay | Haploid | No | Yes | Yes (haploid cells only) | 36 |
| PRIME | Prime editing | Yes (PE3) | No | Growth proliferation assay | Diploid | No | No | N/A | 34 |
| beeSTING-seq | Base editing | Yes | No | Single cell RNA sequencing | Diploid | No (no editing frequency adjustment) | No | N/A | 26 |
| Perturb(BE)-seq | Base editing | Yes | No | Single cell RNA sequencing | Diploid | No (no editing frequency adjustment) | Yes, but not in same cells as RNA is measured (pooled genotyping) | No | 24 |
| Base editing screens | Base editing | Yes | No | FACS based assay, cellular fitness | Diploid | No (no editing frequency adjustment) | No | No | 25, 27 |
| BEAN (Base Editor screen analysis with Activity Normalization) | Base editing | Yes | No | LDL cholesterol uptake - flow cytometry based assay | Diploid | No (Does not measure gene expression, but does adjust for editing frequency on the phenotype) | Yes (sequences a reporter construct to infer endogenous editing efficiencies) | Yes | 32 |

**Supplementary Table 2: Target edits and pegRNA sequences for the PPIF 5’ splice site and HEK3 locus**.

pegRNA designs for the PPIF 5’ splice site edits and the HEK3 locus experiment, for prime-editing optimization. Each row represents one designed edit and includes position and sequence information for the edit and the corresponding pegRNA. The pegRNA designs here used the Scaffold 1 RNA sequence (shown in **Supplementary Fig. 20**).

**Supplementary Table 3: Target edits and pegRNA sequences for the Variant-FlowFISH screens**.

pegRNA designs for all screening experiments. Each row represents one designed edit and includes position and sequence information for the edit and the corresponding pegRNA. The Flip + Extension Scaffold RNA is used here for all of the pegRNA designs (sequence shown in Supplementary Fig. 20).

**Supplementary Table 4: Expression of transcription factors selected for 8mer insertion screen**.

Transcription factors that 8-bp insertion edits were designed to create motif instances for and expression in THP-1, Jurkat, and Jurkat cells stimulated with Phorbol 12-myristate 13-acetate (PMA) and anti-CD3 antibody. Transcription factor gene expression is reported as the mean transcripts per million (TPM) from two RNA-sequencing technical replicates.

**Supplementary Table 5: Motifs of high predicted contribution to chromatin accessibility in THP-1 cells**.

TF-MoDISco was used to cluster and consolidate sequence motifs with high ChromBPNet contribution scores that were enriched in ATAC-seq peaks in THP-1 cells. Motifs discovered by TF-MoDISco (pattern column) are listed with their abundance in accessible peaks (seqlets), their motif pattern (modisco_cwm_fwd; modisco correlation weight matrix), and the top three matches (match, match logo, q value) to transcription factor motifs using TOMTOM from the MEME Suite.

**Supplementary Table 6: Motifs of high predicted contribution to chromatin accessibility in Jurkat cells.**

TF-MoDISco was used to cluster and consolidate sequence motifs with high ChromBPNet contribution scores that were enriched in ATAC-seq peaks in Jurkat cells. Columns are the same as in Supplementary Table 4.

**Supplementary Table 7: Motifs of high predicted contribution to chromatin accessibility in stimulated Jurkat cells.**

TF-MoDISco was used to cluster and consolidate sequence motifs with high ChromBPNet contribution scores that were enriched in ATAC-seq peaks from Jurkat cells stimulated with PMA + anti-CD3 antibody. Columns are the same as in Supplementary Table 4 and 5.

**Supplementary Table 8: Prediction effects from sequence-based deep learning models.**

Listing of effects predicted by Borzoi, Enformer, ProCapNet and ChromBPNet and the corresponding effects on expression measured by Variant-FlowFISH in THP-1 cells for *PPIF* tiling and 8mer insertion edits that passed quality control filters.

**Supplementary Table 9: Summary of designs and measured effects for rationally designed sequence ed**its.

Listing of edits, design metadata, effects on expression predicted by Enformer, transcription factor motif instances predicted to be introduced by edits, and effects on expression measured by Variant-FlowFISH.

**Supplementary Table 10: Primer sequences used for cloning pegRNA libraries.**

This table lists the primers used to amplify the subpools of oligos and add the homology arms for Gibson Assembly into the lentiviral vector SgOpti.

**Supplementary Table 11: Amplicon sequencing primers.**

Summary of the amplicon sequencing primers used in this study. Each primer has an adaptor for the barcoding of PCR samples for the Illumina sequencing platform. The adaptors are highlighted in blue for the forward primer and orange for the reverse primer.

## Supplementary Figures

**Supplementary Figure 1.**
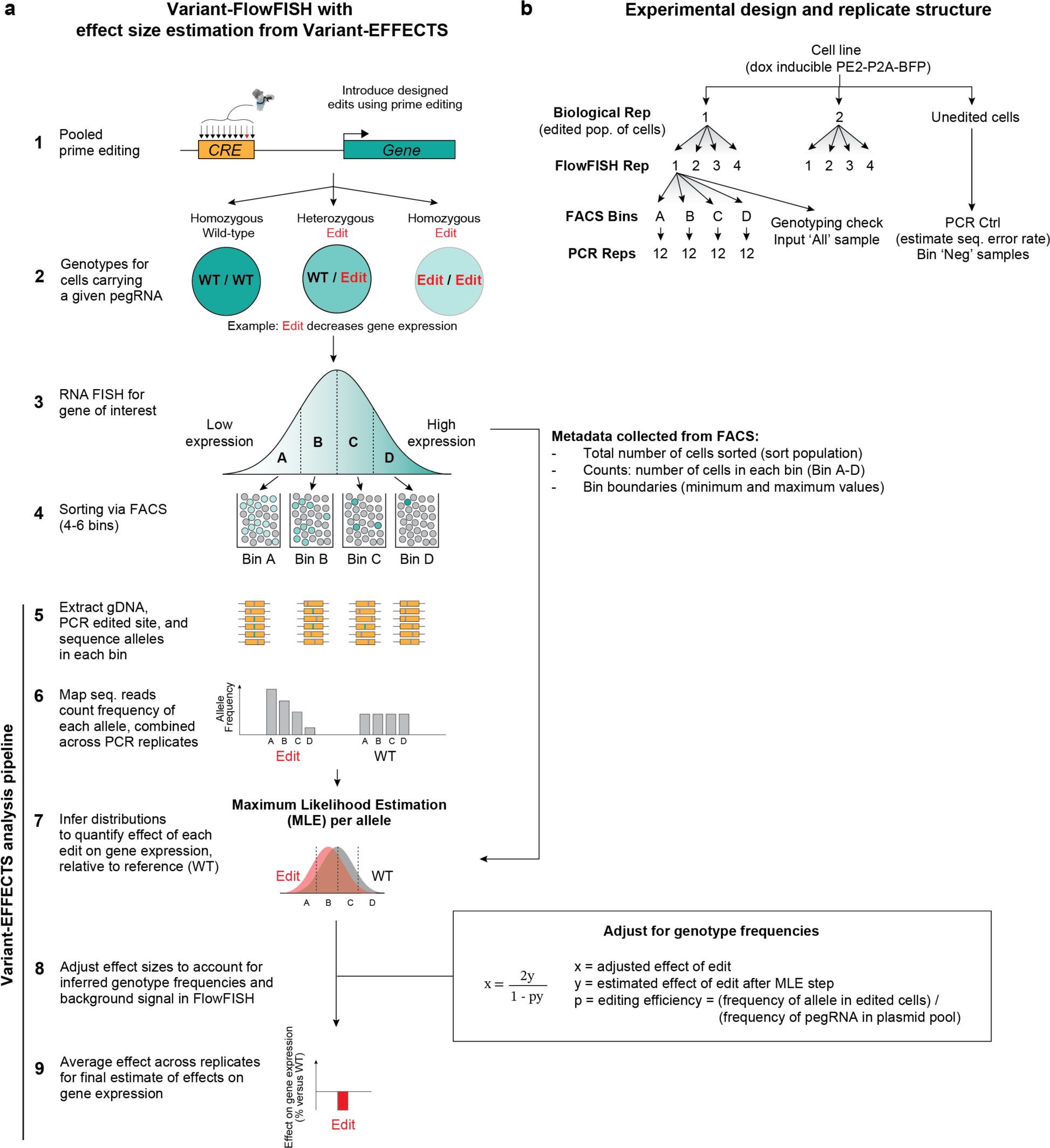
Overview of experimental design of Variant-FlowFISH experiments and computational analysis with Variant-EFFECTS. **(a)** Schematic representing how the data collected from the FACS sort and subsequent sequencing of alleles from cells collected in each of the FACS bins is used to determine the effect of each of the variants on the expression of a target gene. (1) Cells are transduced with a lentiviral pool of pegRNAs designed to introduce edits into a target site of interest. (2) Each pegRNA can create cells carrying 3 genotypes: homozygous wild-type (WT), heterozygous edited, and homozygous edited. In this example, the edit decreases gene expression (light green color), and each of the three genotypes will result in levels of gene expression corresponding to the sum of effects of each allele. (3,4) We conducted RNA FlowFISH and sort cells into expression bins (here: A = low expression, D = high expression). (5) We extract gDNA from cells in each bin, PCR amplify the edited site, and deeply sequence the amplicon using Illumina sequencing. (6) We match sequencing reads to an index of designed edits to quantify the frequency of each allele, including all edits and the wild-type allele, and sum counts across PCR replicates (see panel **b**). (7) We use these allele frequencies to infer the mean and variance of the distribution of each allele in gene expression space using maximum likelihood estimation (MLE, see Methods). The MLE step uses as input the allele frequencies from Step 6 as well as FACS metadata collected from Step 3. (8) The distributions of alleles are affected by the editing frequency, because heterozygous vs homozygous edited cells will have different expression levels (see Methods). We apply a correction step to account for this (box inset, right). We also adjust for background FlowFISH signal (see Methods). (9) We average the effect sizes across technical and biological replicates for the final estimate of effects on gene expression. (**b**) Experimental setup for a representative FlowFISH experiment. Biological replicates are populations of cells independently infected with a lentiviral pegRNA pool. Cells are split into “FlowFISH technical replicates” just before FlowFISH, and sorted into 4-6 expression bins; an “input” unsorted sample (referred to as input ‘All’) is taken to measure allele frequencies in the entire population of cells. Cells from sorted bins are split into “PCR replicates” for library preparation and sequencing, such that each PCR replicate includes ∼250,000 cells per PCR reaction. Unedited cells are also sequenced (“PCR Ctrl”) to estimate the error rate due to Illumina sequencing (referred to as Bin negative or ‘Neg’).

**Supplementary Figure 2.**
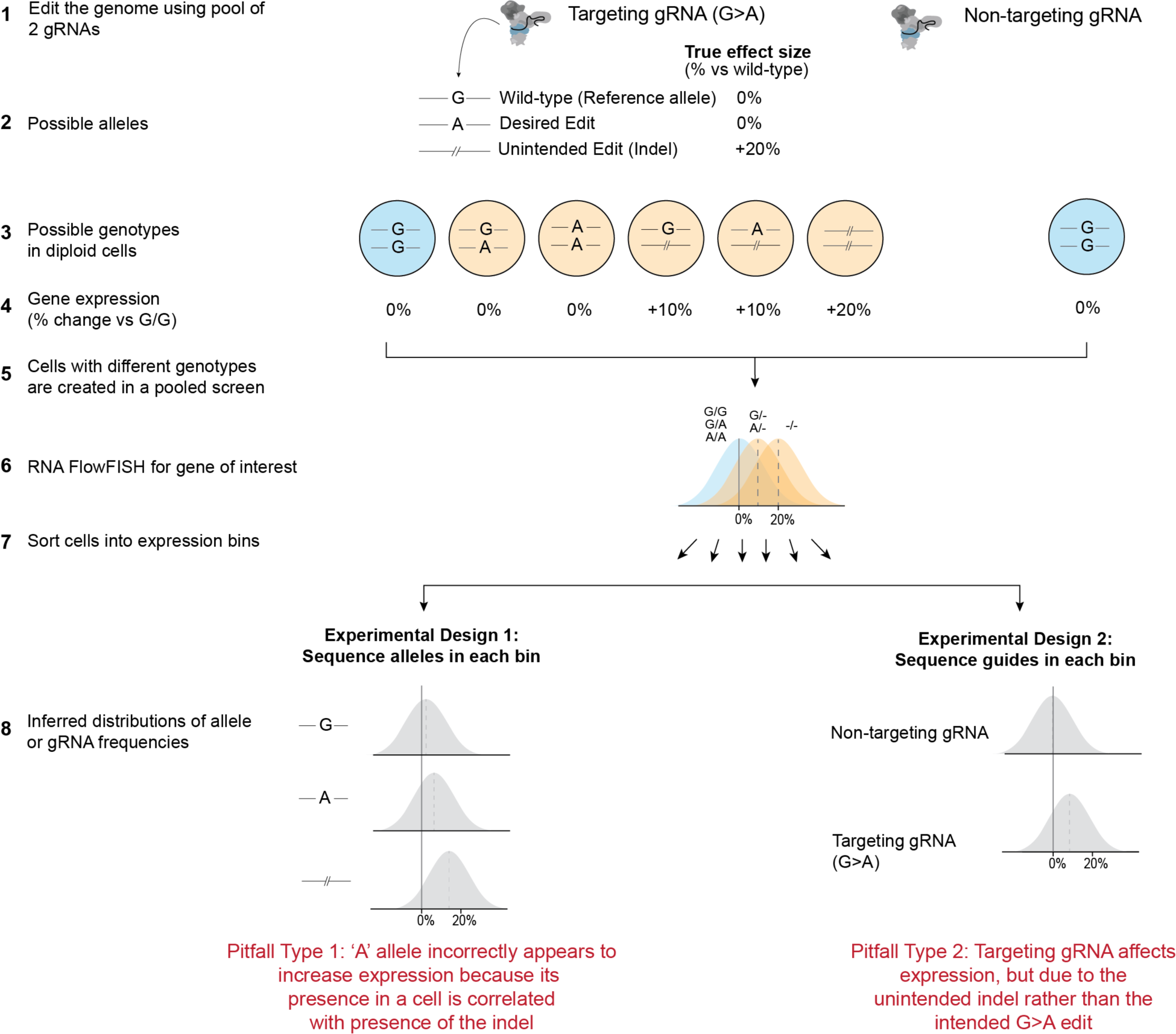
Unintended edits can confound interpretation of CRISPR screens aiming to measure effects of genetic variants. CRISPR editors that introduce multiple edits at high frequencies—such as CRISPR homology-directed repair, base editing, and PE3 prime editing—can confound the results of CRISPR screens aiming to measure the effects of specific variants. (1) This figure presents a hypothetical example of a pooled screen that uses two gRNAs: one to introduce a G>A variant at a particular position, and one non-targeting gRNA that does not lead to editing at this site. (2) In this hypothetical example, the G>A variant has no effect on gene expression, but the gRNA introducing this variant also creates an unintended indel that leads to a +20% effect on gene expression. (3,4) In diploid cells that receive the targeting gRNA, 6 possible genotypes are produced, with the levels of gene expression indicated (average of the effects of the two alleles in the cell). In cells that receive the non-targeting gRNA, all cells remain wild-type (G/G) at the targeted site. (5) These cells with different genotypes are created in a pool in the screen. (6,7) The cells are sorted into bins based on their expression of interest. The blue and orange distributions represent the RNA levels measured for cells with different genotypes. (8) At this point, two alternative experimental designs are to sequence the alleles in each bin (as done in Variant-FlowFISH; left) or to sequence the gRNAs present in each bin (right). In the first case, we obtain the marginal frequency distributions of each of the 3 alleles across expression bins (dotted line: mean of distribution, *i.e.* a naive estimate of the effect size of each allele when compared to the wild-type G allele). Note that the A allele will appear shifted toward higher expression because its presence in cells is correlated with the presence of an indel; that is, some cells carrying the ‘A’ allele, which has no effect, will also carry an indel on the other homolog, which does have an effect on expression. The G allele is shifted as well, but to a lesser extent because some ‘G’ alleles come from wild-type cells (G/G) carrying the non-targeting gRNA. Thus, a naive analysis that does not account for this correlation will mistakenly conclude that the ‘A’ allele affects gene expression, when in fact it has no effect. In the second case (sequencing guides in each bin), the non-targeting gRNA will appear to have zero mean effect and the targeting gRNA will have a positive effect. However, the effect of this gRNA is not due to the intended edit (‘A’ allele), but rather due to the fact that this gRNA also unintentionally introduces a functional indel. As such, a naive analysis that does not account for this correlation will mistakenly conclude that the ‘A’ allele affects gene expression. Variant-FlowFISH avoids both of these pitfalls by using the PE2 system (which has a very low rate of unintended edits). Notes:

- While the example here is drawn to illustrate a sorting-based screen, the same challenge applies to screening formats such as Perturb-seq screens that read out expression and the identity of the gRNA (similar to Experimental Design 2).
- The degree of confounding depends on multiple factors including the frequencies of each allele, the strength of their effect sizes, the covariance of the alleles in diploid cells, and the distribution of other effect sizes in the pool.

**Supplementary Figure 3.**
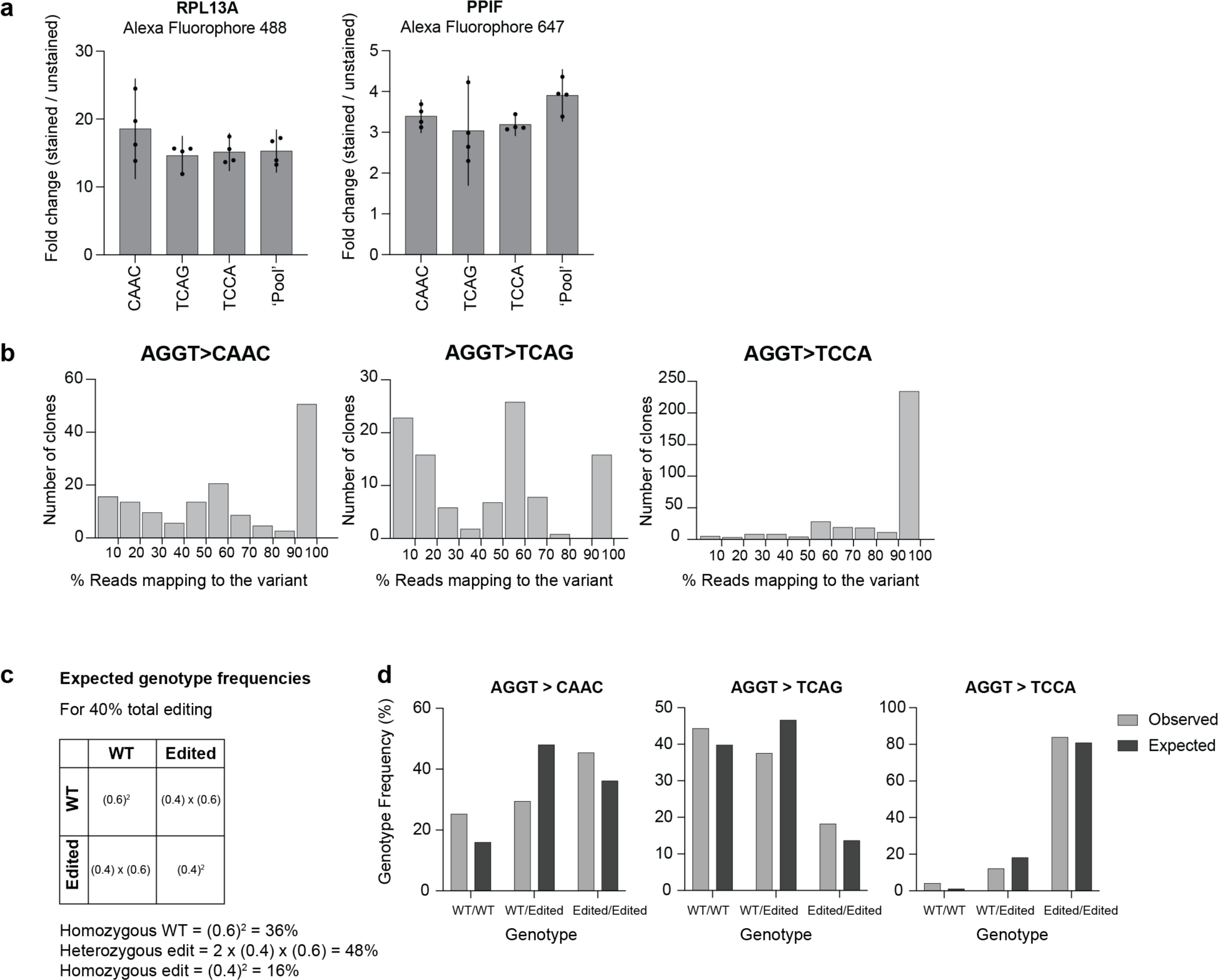
PPIF 5’ splice site RNA FlowFISH fold changes and genotyping PPIF 5’ splice site clonal populations. (**a**) Fold changes (stained / unstained) as determined via FACS for the control gene RPL13A (Alexa Fluorophore 488) and PPIF gene (Alexa Fluorophore 647). A fold change greater than 2 is acceptable for the RNA FlowFISH assay. (**b**) Clonal populations were derived for each of the PPIF 5’ splice site edits: AGGT>CAAC, TCAG and TCCA. Homozygous edited clones were defined as containing 90-100% of the reads mapping to the intended ‘edited’ sequence; homozygous reference clones were defined as 0-10%. Plots represent the following number of clones analyzed: AGGT>CAAC (n=149), TCAG (n=105) and TCCA (n=358). Note that the frequency of “clones” with an intermediate level of editing (not 0%, 50%, or 100%) indicates that some of these “clones” may represent expansions of several single cells. (**c**) An example calculation showing how the ‘expected’ editing rate can be calculated for a given variant. (**d**) Plots comparing the observed vs.‘expected editing rate for each of the 5’ splice site variants. Genotypes are as indicated: WT/WT (Homozygous WT), WT/Edited (Heterozygous Edited), Edited/Edited (Homozygous Edited).

**Supplementary Figure 4.**
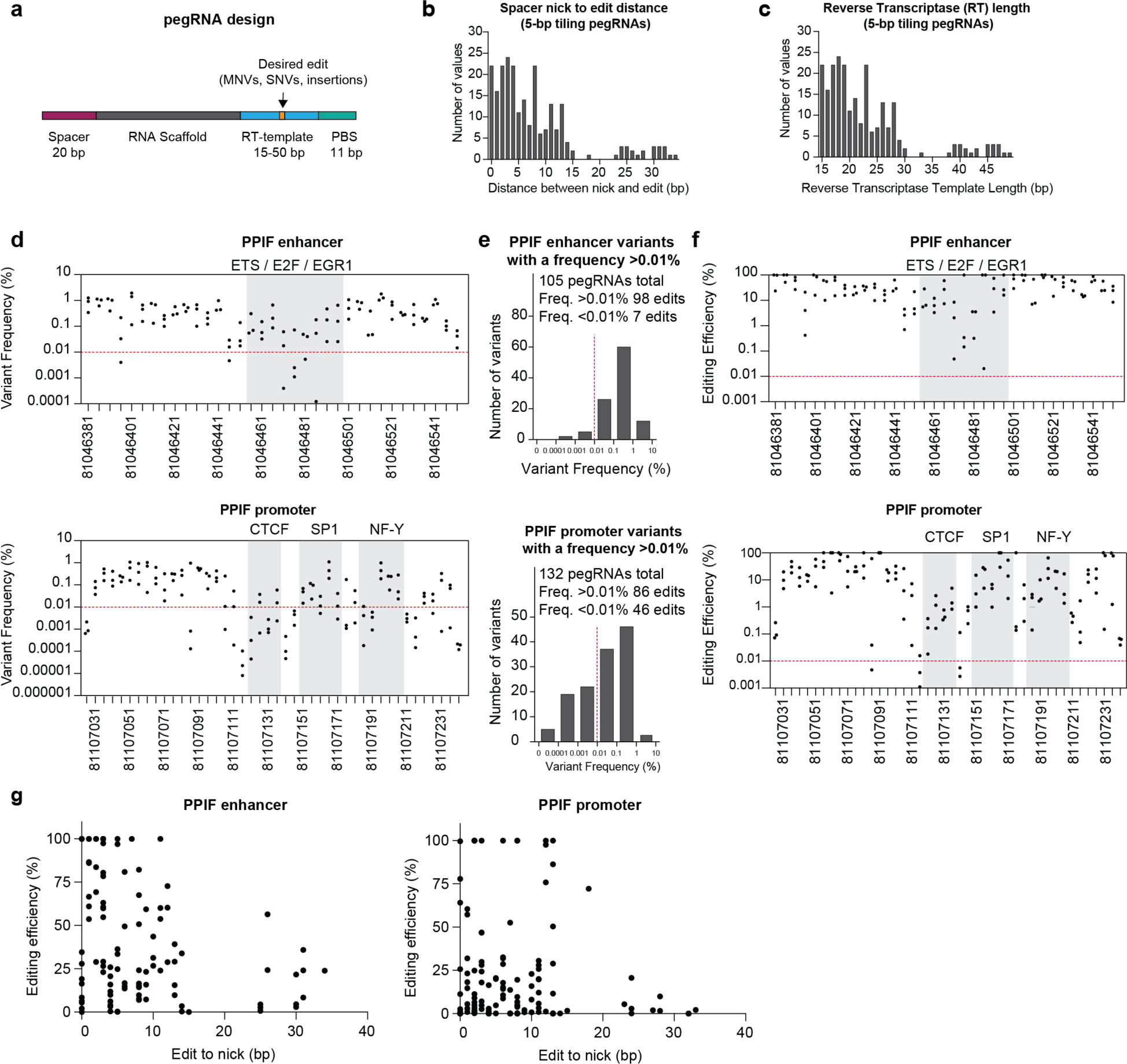
The design of the prime editing guide RNA (pegRNAs) and the frequency of variants for the prime editing tiling mutagenesis experiments at the PPIF enhancer and promoter. (**a**) A schematic of the pegRNA design with the various lengths of the spacer, reverse-transcription (RT) template and primer binding site (PBS) indicated. (**b**) A frequency histogram of the distance between the ‘nick and edit’ (bp) for each of the pegRNAs designed at the PPIF locus: enhancer and the promoter. (**c**) A frequency histogram for the reverse transcriptase (RT) template length (bp) for each of the pegRNAs designed to target the PPIF locus. (**d**) The frequency of variants (%) mapped to the PPIF enhancer and promoter. Dots represent the average variant frequency for each of the 5-bp substitutions across FlowFISH replicates (*n*=8). A dotted red line is shown at a frequency of 0.01%. Binding sites for transcription factors at the PPIF enhancer and promoter are indicated by gray shading. (**e**) Frequency histograms for the PPIF enhancer and promoter highlight the number of variants which are at a frequency >0.01%. (**f**) The editing efficiency of each of the 5-bp tiling variants mapped back to the genomic coordinates for the PPIF enhancer and promoter. Dots represent each of the three 5-bp substitutions per targeted tile, across FlowFISH replicates (*n*=8). A red line indicates an efficiency of 0.01%. Binding sites for transcription factors at the PPIF enhancer and promoter are indicated by gray shading. (**g**) Editing efficiency (%) vs. the distance of the edit position relative to ‘nick’ (bp) for the pegRNA. Dots represent each 5-bp substitution.

**Supplementary Figure 5.**
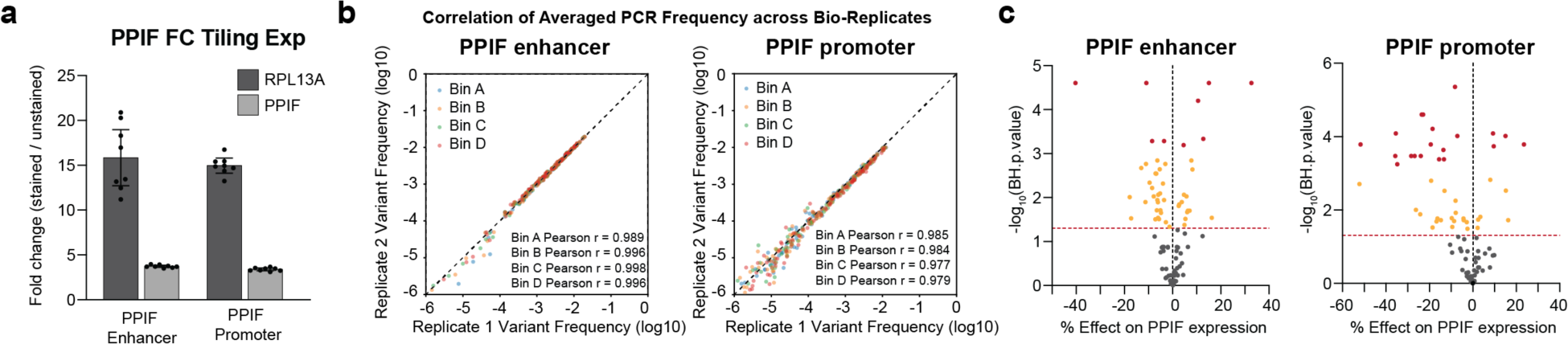
Fold-change and correlation plots highlighting the technical details of the Variant-FlowFISH assays at the PPIF loci and volcano plots representing the effect sizes of each of the variants. (**a**) A bar plot representing the fold change (FC; stained/unstained) for each of the 5-bp tiling mutagenesis Variant FlowFISH experiments at the PPIF locus. Bars: mean +/- 95% c.i. Dots: 4 FlowFISH technical replicates from each of 2 biological replicates (n=8). (**b**) Correlation of averaged variant frequencies across biological replicates for the PPIF enhancer and promoter Variant-FlowFISH experiments in THP-1 cells. Text at bottom right reports correlations for the variant frequencies from each of the FACS Bins (A-D). (**c**) Volcano plots for the PPIF enhancer and promoter highlight the effect size for each of the 5-bp edits on PPIF gene expression and the level of significance (*q* = Benjamin-Hochberg p-value, two-sided single-sample t-test). Dots represent each individual 5 bp variant. Red line: *q* = 0.05. Red dots: *q* < 0.001. Yellow dots: *q* < 0.05. Gray dots: Not significant.

**Supplementary Figure 6.**
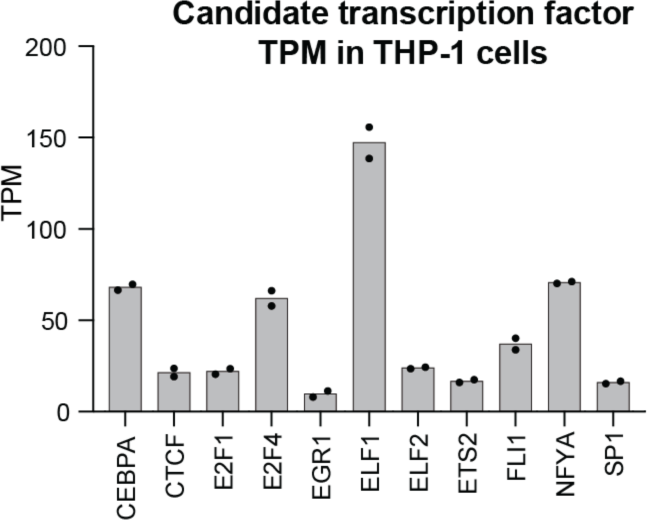
Expression levels of candidate transcription factors binding to the *PPIF* enhancer and promoter in THP-1 cells. RNA expression from bulk RNA-seq in transcripts per million (TPM). Candidate transcription factors binding to the PPIF enhancer or promoter were selected via a FIMO analysis (using a HOCOMOCO v11 database) of the sequences in these regulatory regions, with candidate factors having a TPM >8. Bars: mean. Dots: 2 technical replicates.

**Supplementary Figure 7.**
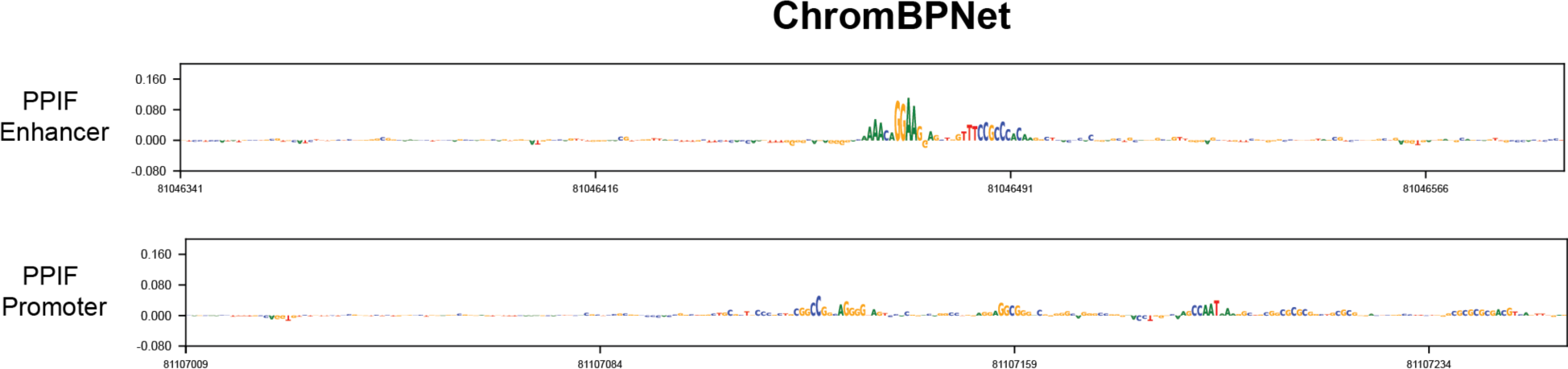
ChromBPNet^42^ predicts chromatin accessibility in THP-1 cells at the PPIF enhancer and promoter. ChromBPNet sequence interpretations (DeepSHAP) of the reference sequence at each of these loci show the predicted contribution of each nucleotide for chromatin accessibility signal.

**Supplementary Figure 8.**
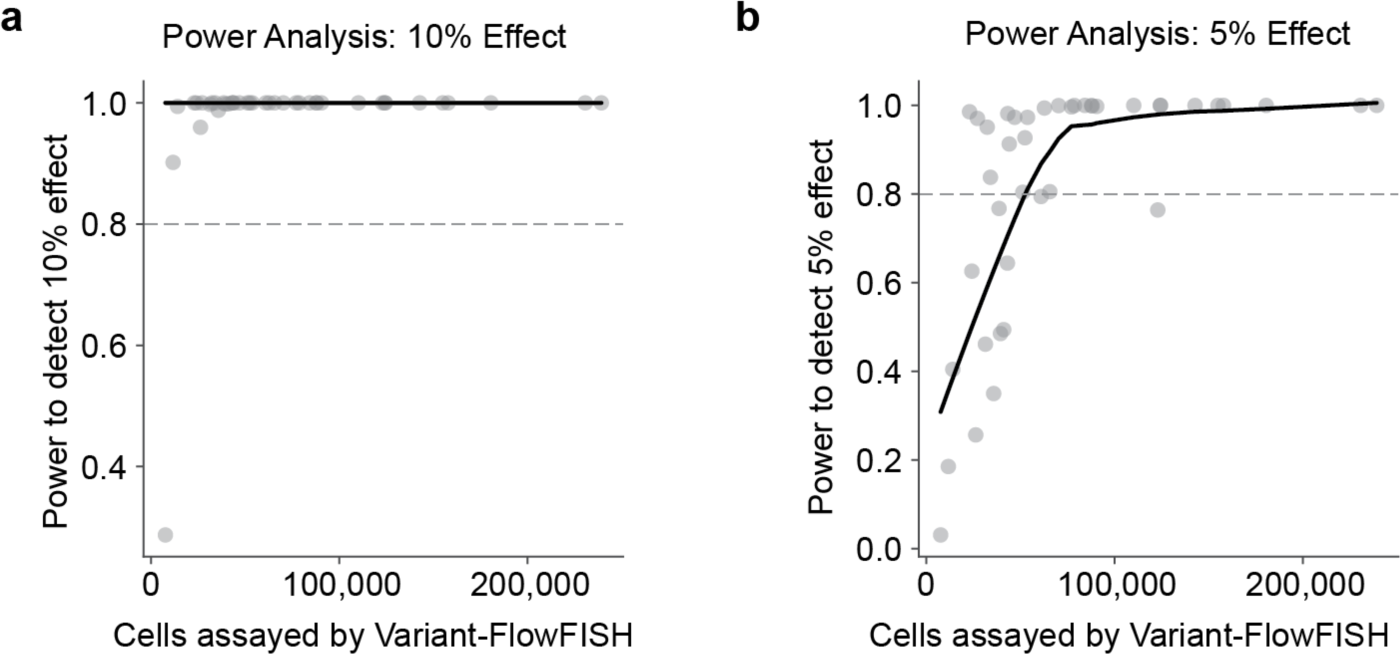
Power analysis of 8mer insertion screen in THP-1 cells. Relationship between power to detect a (**a**) 10% effect or (**b**) 5% effect on expression (*q <* 0.05) and total cells assayed by Variant-FlowFISH for each 8-mer insertion screened in THP-1 monocytes. Power at a given effect size was calculated by using the number of FlowFISH replicates and observed standard deviation in THP-1 cells for every 8mer insertion in the screen. Solid line represents nonparametric lowess regression of the power on number of cells assayed. Dotted line represents 80% power.

**Supplementary Figure 9.**
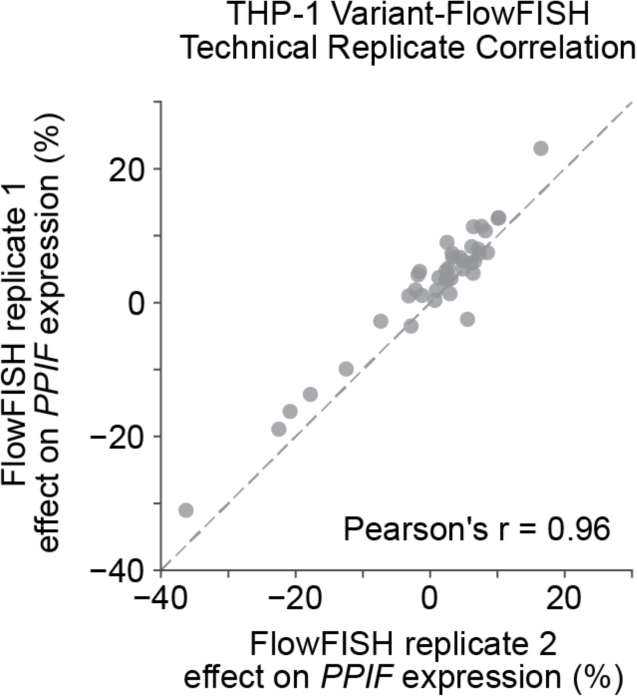
Reproducibility of measurements across technical Variant-FlowFISH replicates for 8mer insertion screen in THP-1 cells. Correlation of observed Variant-FlowFISH effect sizes across two technical replicates derived from the same biological replicate.

**Supplementary Figure 10.**
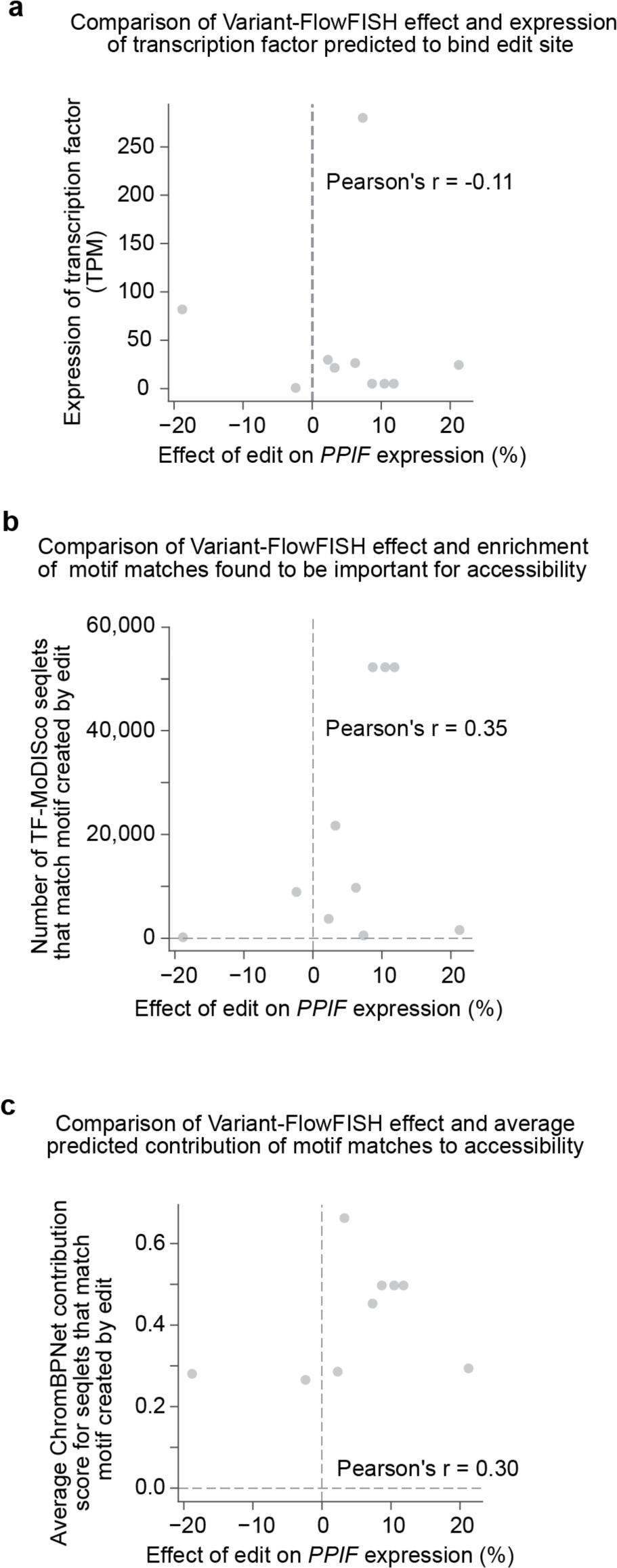
Correlation of 8mer insertion effets to common heuristics for the global activities of corresponding transcription factors in THP-1 cells. (**a**) Correlation between the effect size of transcription factor binding site insertion edits and expression of the transcription factors predicted to bind them for the subset of edits with strong *de novo* binding site predictions and non-zero TF expression in THP-1 cells. Edits were included if they created a transcription factor motif instance identified by comparing ChromBPNet DeepSHAP prediction to TF-MoDISco motifs with TOMTOM from MEME Suite. (**b**) Correlation between the effect size of transcription factor binding site insertion edits and frequency of motif matches for TF-MoDISco hits in THP-1 cells (a measure of the genome-wide importance of this transcription factor in influencing chromatin accessibility as estimated by ChromBPNet). (**c**) Correlation between effect size of transcription factor binding site insertion edits and average predicted importance (predicted effect on chromatin accessibility in a 1 kb window by ChromBPNet) of corresponding motif matches for TF-MoDISco hits in THP-1 cells.

**Supplementary Figure 11.**
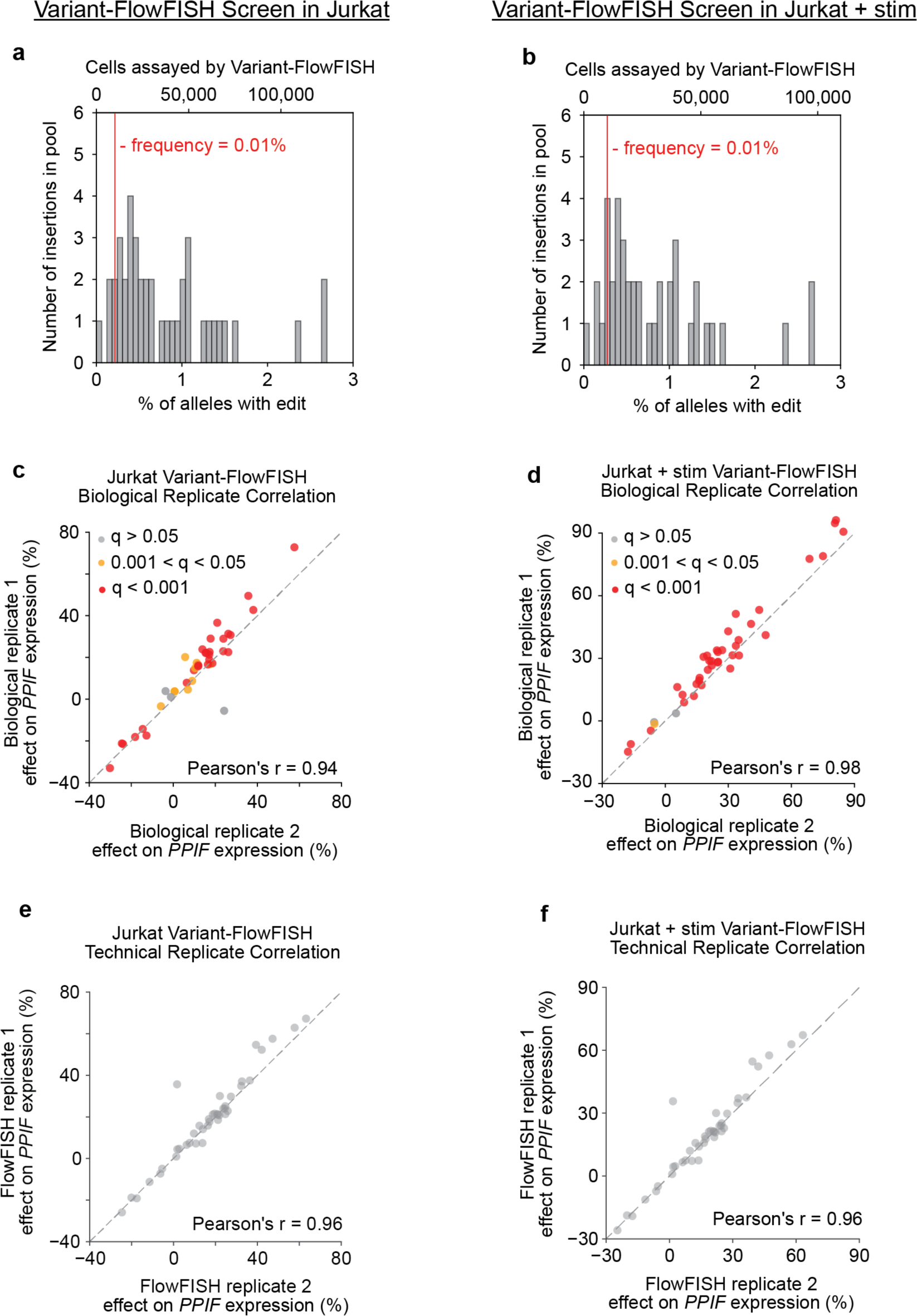
Edit frequencies, cell coverage and reproducibility of effects measured by Variant-FlowFISH for 8mer insertion screen edits in Jurkat cells with or without stimulation. (**a, b**) Histograms of frequencies of each 8-bp insertion in the edited pool of cells as a percentage of alleles (bottom x-axis) and the corresponding minimum number of cells assessed by Variant-FlowFISH (top x-axis) in (**a**) Jurkat cells and (**b**) Jurkat cells stimulated with PMA + anti-CD3 antibody. Bottom x-axis: frequency of edit in all sequencing reads. Top x-axis: Minimum number of cells assessed by Variant-FlowFISH. (**c, d**) Correlation of observed Variant-FlowFISH effect sizes across biological replicates in (**c**) Jurkat cells and (**d**) Jurkat cells stimulated with PMA + anti-CD3 antibody. Dots (*n*=41): all variants passing the frequency threshold. Red: *q* < 0.001. Yellow: *q* < 0.05, gray: *q* > 0.05 (Benjamini-Hochberg corrected *p*-value, one-sample t-test). (**e, f**) Correlation of observed Variant-FlowFISH effect sizes across two representative technical replicates in (**e**) Jurkat cells and (**f**) Jurkat cells stimulated with PMA + anti-CD3 antibody.

**Supplementary Figure 12.**
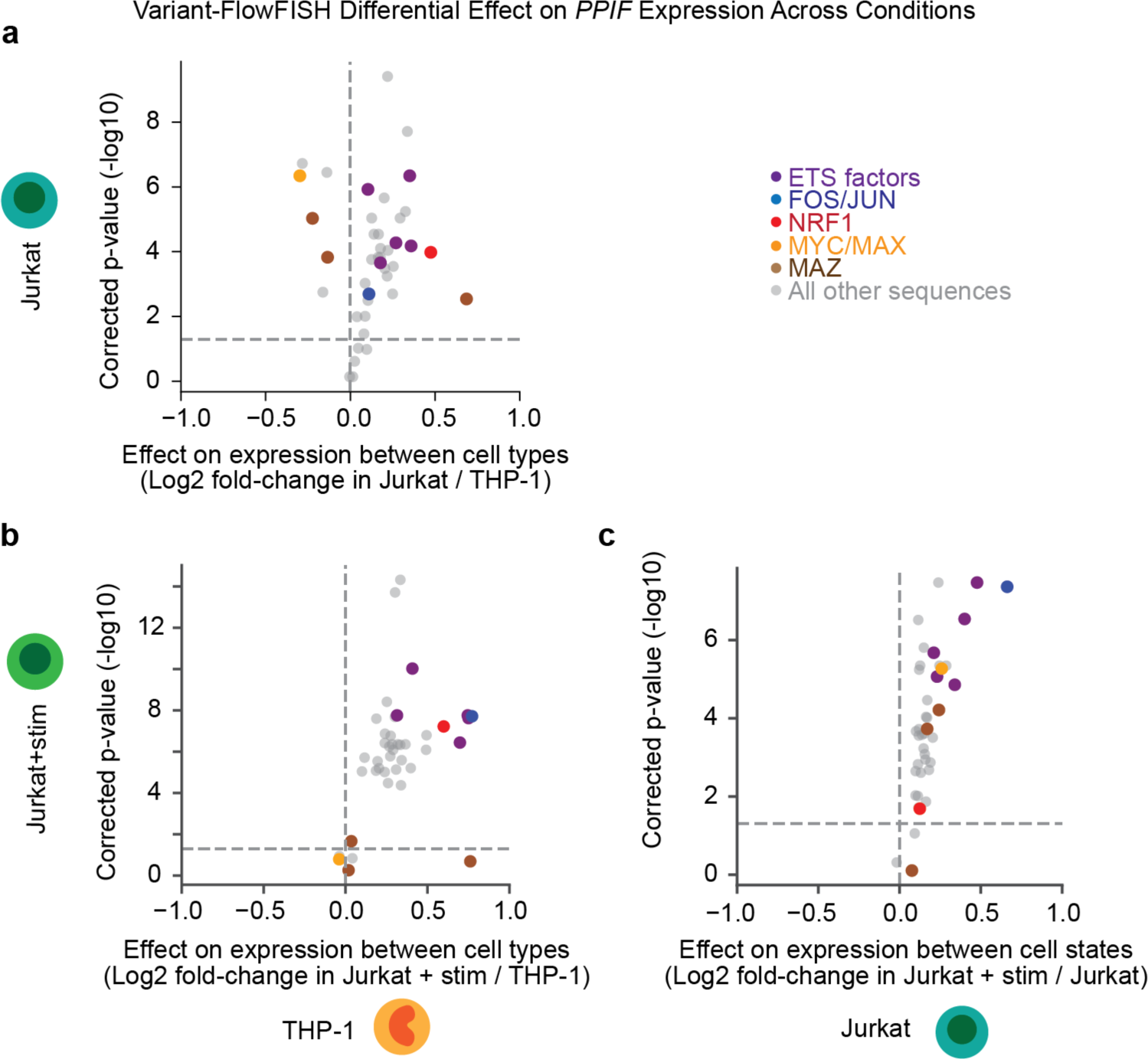
Differential effects of 8mer insertions on *PPIF* expression across conditions. Differential effects of 8-bp insertions on *PPIF* expression between (**a**) Jurkat cells and THP-1 cells, (**b**) Stimulated Jurkat cells and THP-1 cells, and (**c**) Stimulated and unstimulated Jurkat cells. Differential effects are the log2 fold changes of the effect sizes (second/first condition listed in a-c) and significantly different effects between conditions were determined using two-sided, two sample Welch’s t-test with a Benjamini-Hochberg multiple test correction. Horizontal dotted lines represent a corrected p-value of 0.05. Dots (*n*=41): all variants passing the frequency threshold. Dots are colored if the edit creates a predicted *de novo* motif instance of a transcription factor binding site (see legend).

**Supplementary Figure 13.**
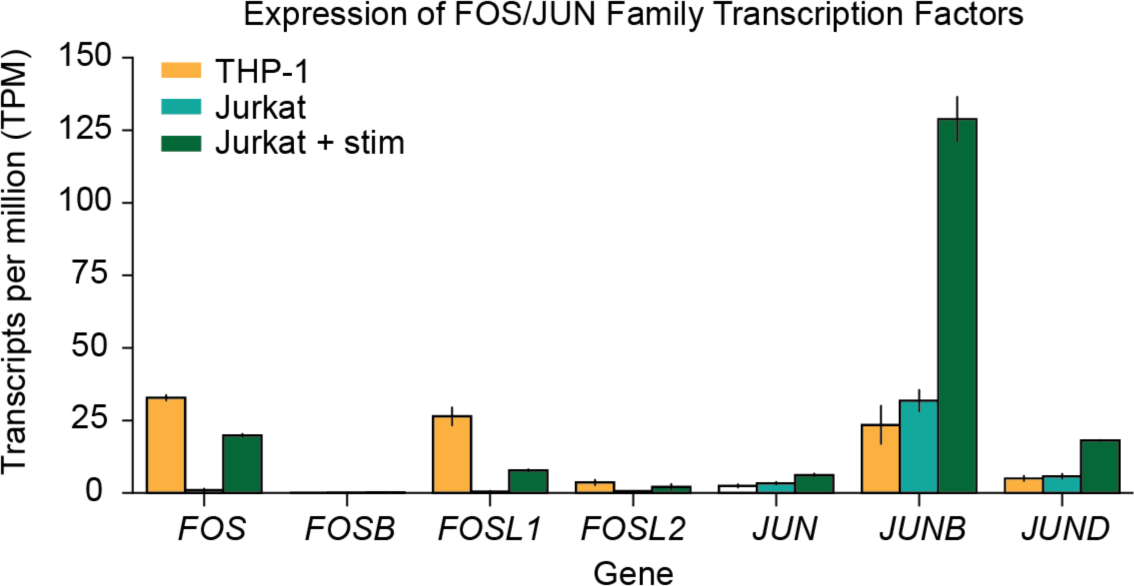
Gene expression of FOS and JUN family transcription factors. Expression of genes in the FOS and JUN transcription factor (TF) families across the three conditions as measured by RNA-seq in wild-type cells and represented as transcripts per million (TPM).

**Supplementary Figure 14.**
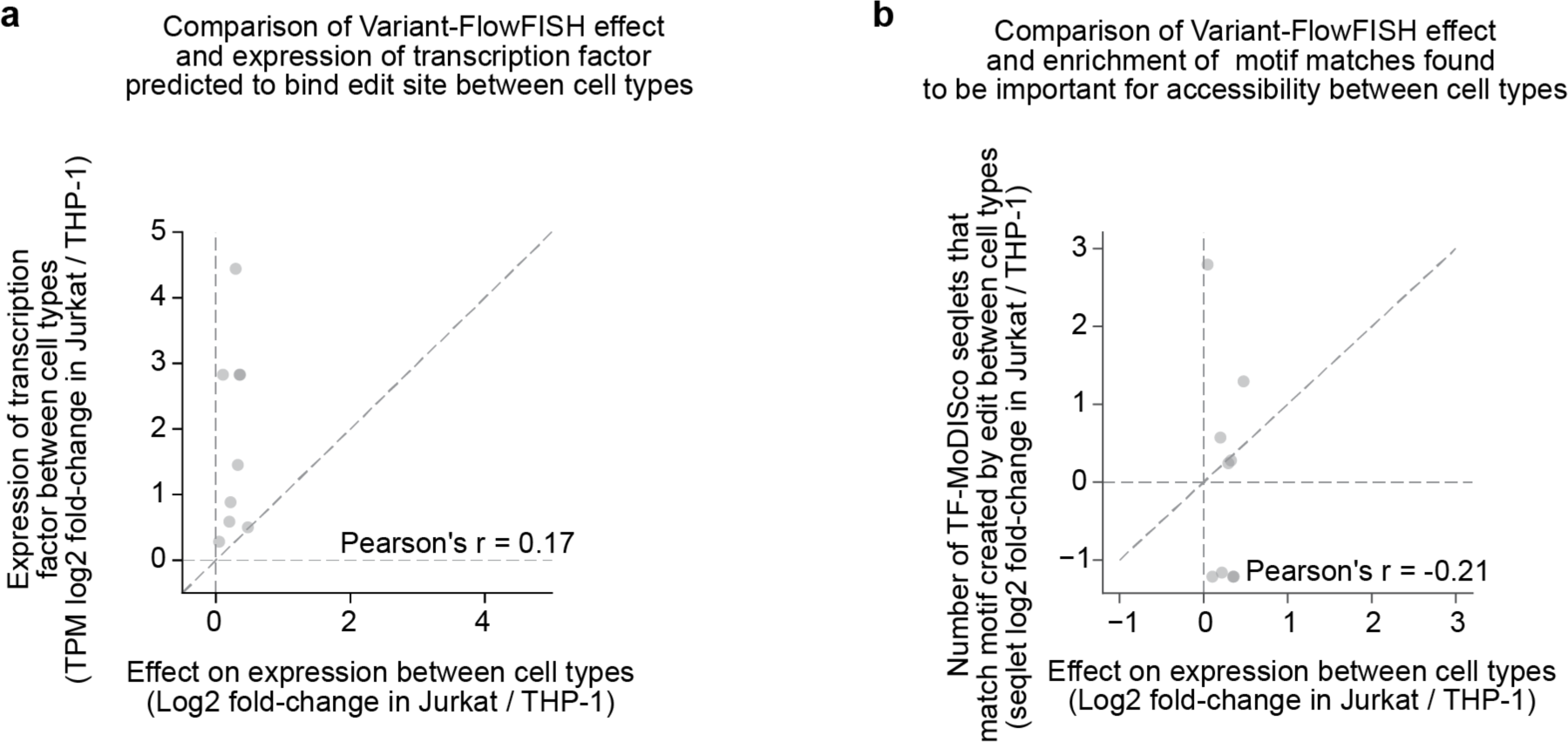
Correlation of change in 8mer insertion effects to change in common heuristics for global activities of corresponding transcription factors between cell types. (**a**) Correlation between the change in effect size of transcription factor binding site insertion edits across cell types and the change in expression of the transcription factors predicted to bind them for the subset of edits with strong *de novo* binding site predictions and non-zero transcription factor expression in both cell types. (**b**) Correlation between the effect size of transcription factor binding site insertion edits across cell types and change in frequency of motif matches for TF-MoDISco hits across cell types (for transcription factor motifs found by TF-MoDISco in both cell types).

**Supplementary Figure 15.**
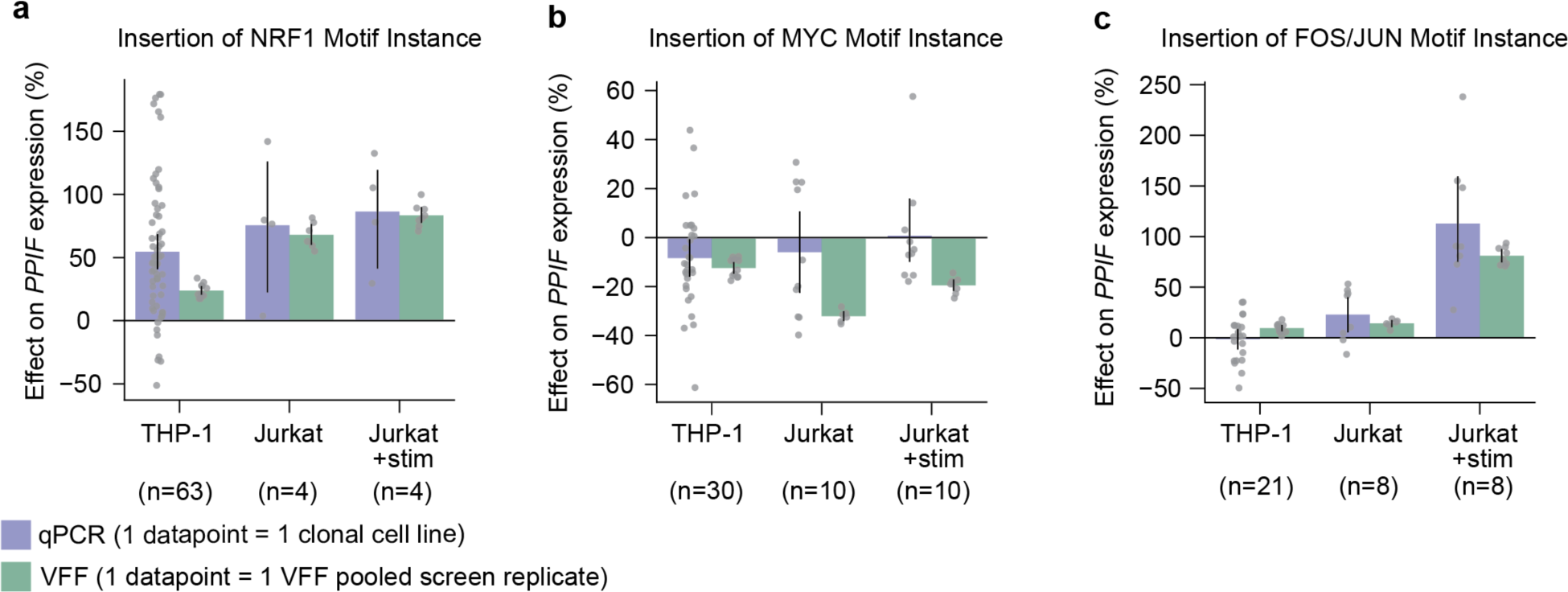
Validation of 8mer insertion effects with qPCR in clonally-derived cell lines. We derived homozygous edited clonal THP-1 and Jurkat cell lines for each of three 8-mer insertions, which created predicted binding sites for (**a**) NRF1, (**b**) MYC, or (**c**) FOS/JUN. Expression of *PPIF* was measured in each clonally derived cell line (gray dots) using qPCR. Number of edited clones are shown in blue below each condition. Effects are normalized to homozygous wild-type clones in the corresponding cell type (n = 162, 48, and 48 clones (edited + wild-type) for THP-1, Jurkat, and Jurkat+stim, respectively).

**Supplementary Figure 16.**
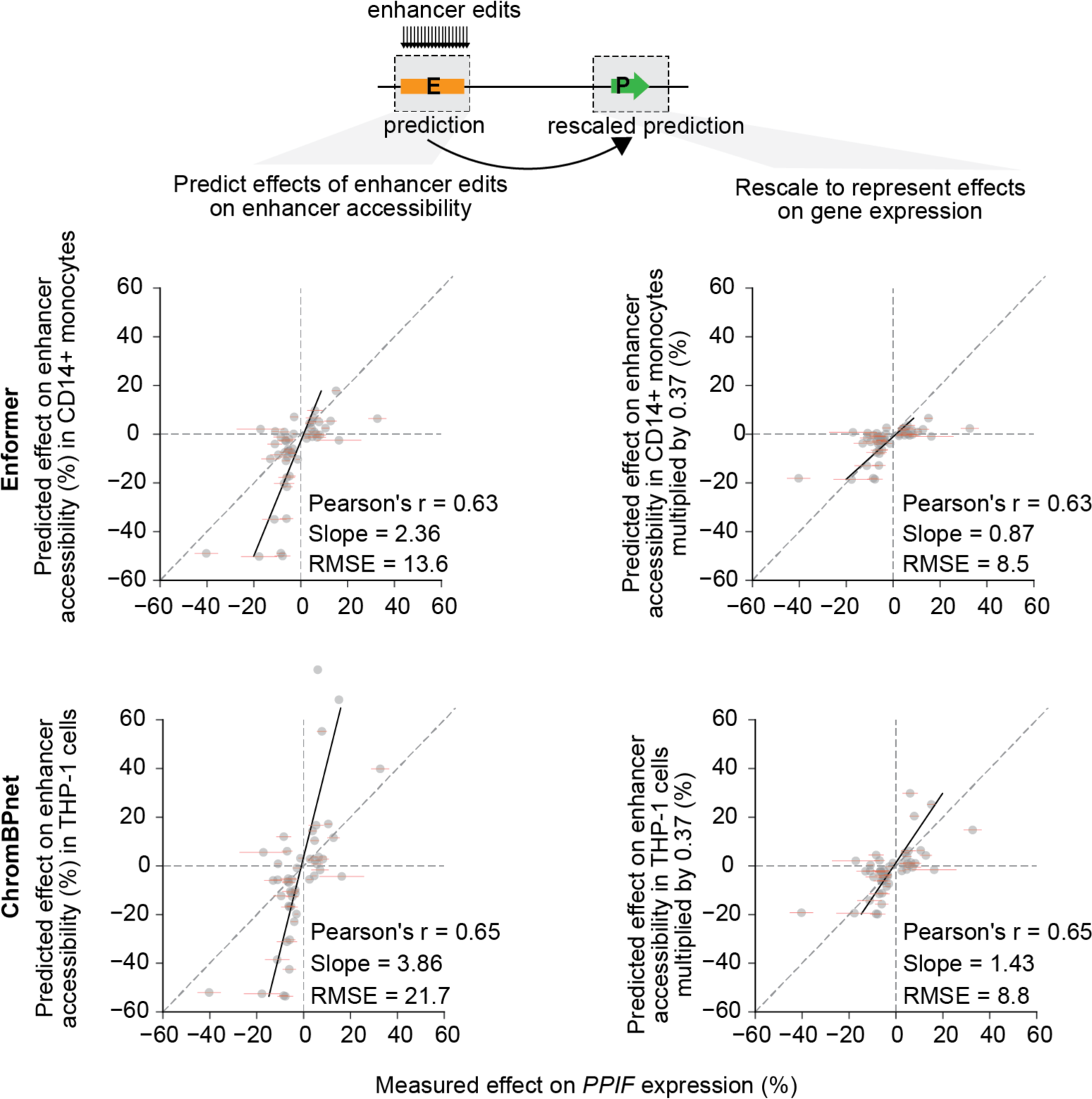
Rescaling predicted effects of variants on accessibility at the *PPIF* enhancer to predict effects on *PPIF* gene expression. We considered an alternative approach to assessing the effects of enhancer variants on gene expression, based on the notion that variants in a distal enhancer should locally affect the activity of that enhancer, and that altered enhancer activity should lead to a corresponding linear effect on gene expression. We computed the predicted effects of variants on accessibility at the *PPIF* enhancer and multiplied these effects by the measured effect of the enhancer on gene expression, which we previously quantified using CRISPRi-FlowFISH (37%). Each scatterplot shows the correlation of effects on PPIF expression measured by Variant-FlowFISH with predicted effects on PPIF enhancer accessibility signal (left) and correlation using enhancer edit effects on the enhancer multiplied by the experimental effect of enhancer CRISPRi perturbation (0.37) on *PPIF* expression (right). The bottom row is the same as Fig. 4d. Top row shows similar analyses for Enformer. Approaches in the right column perform better than directly predicting the effects of edits at the enhancer on gene expression using Enformer (shown in Fig. 4b). Black line: Linear regression line of best fit. Legend lists Pearson’s *r* correlation coefficient, slope from the linear regression, and root mean squared error (RMSE) of the predicted effects on expression (%).

**Supplementary Figure 17.**
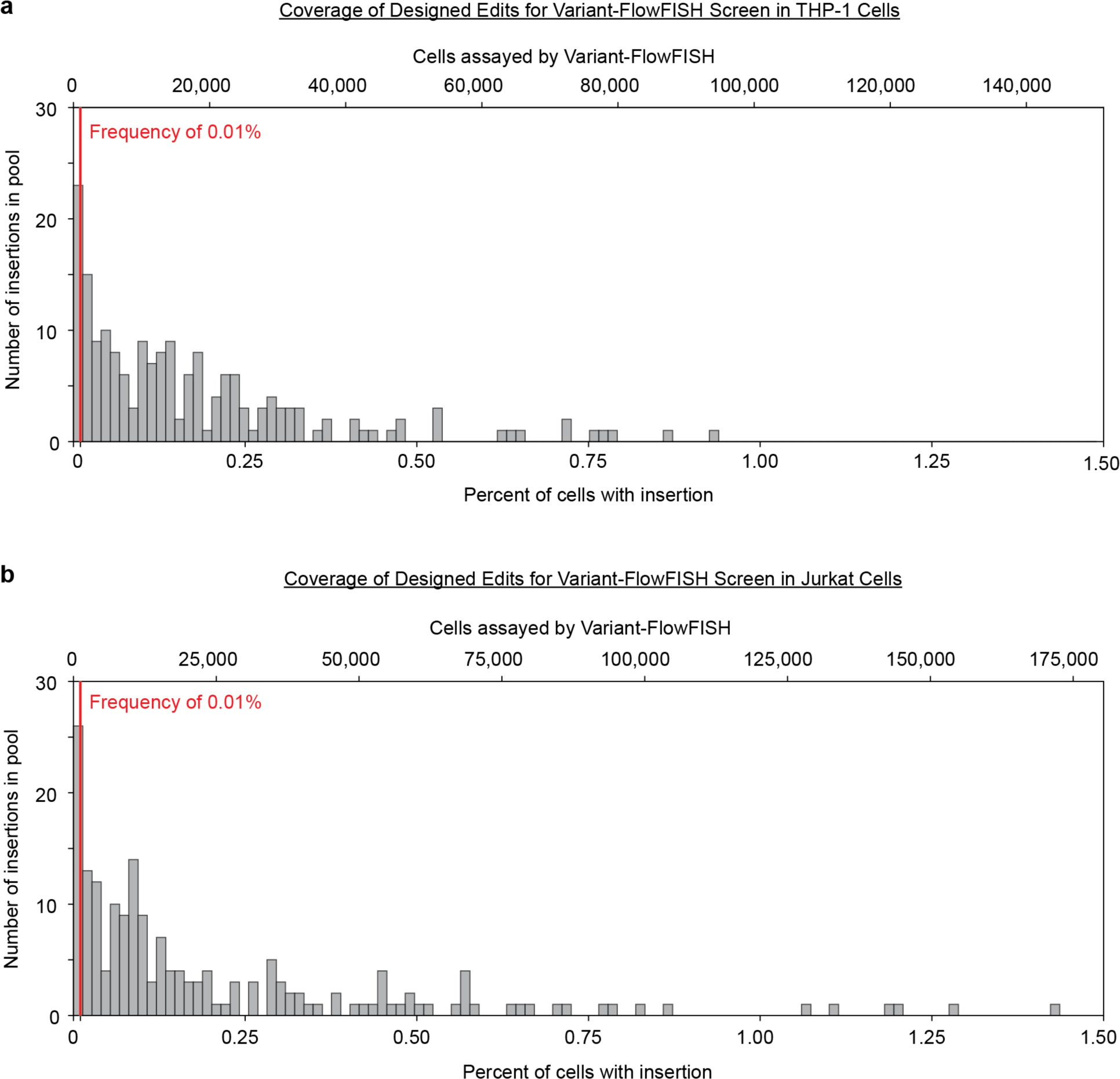
Edit frequencies and cell coverage for Variant-FlowFISH screens in THP-1 and Jurkat for rationally designed sequence edits. Histograms of frequencies of rationally designed sequence edits in the edited pool of cells as a percentage of alleles (bottom x-axis) and the corresponding minimum number of cells assessed by Variant-FlowFISH (top x-axis) in (**a**) THP-1 cells and (**b**) Jurkat cells.

**Supplementary Figure 18.**
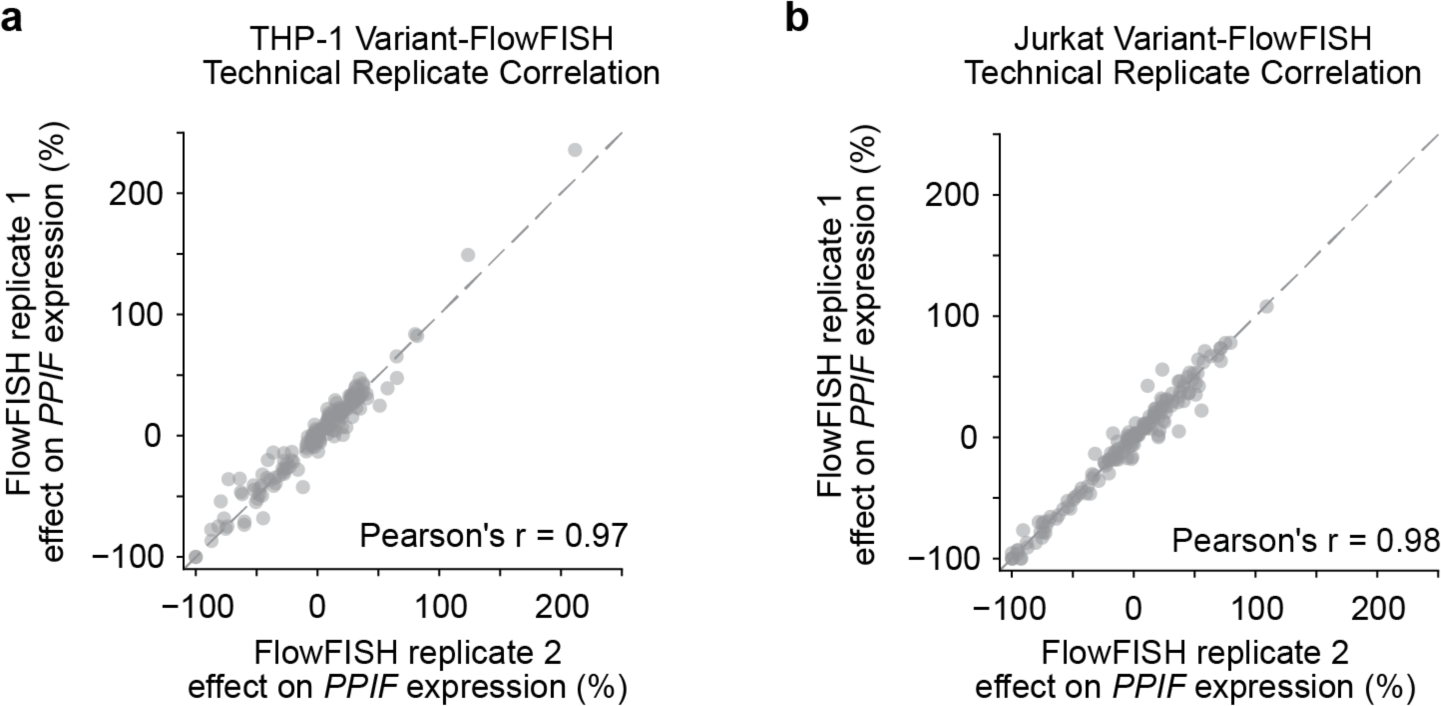
Technical replicate reproducibility of effects measured on *PPIF* expression for rationally designed sequence edits. Correlation of observed Variant-FlowFISH effect sizes across two representative technical replicates in (a) THP-1 cells and (**b**) Jurkat cells n = 164 edits).

**Supplementary Figure 19.**
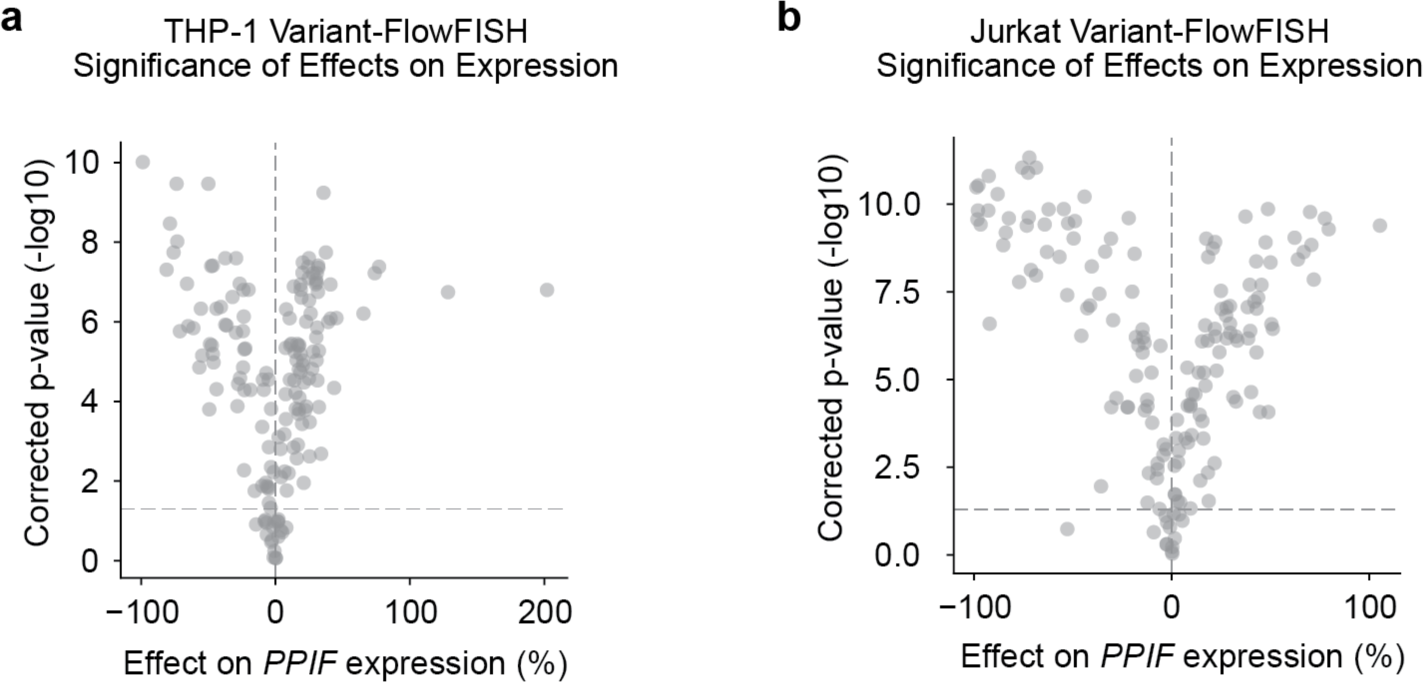
Effect sizes and *p*-values for rationally designed sequence edits in THP-1 and Jurkat. Volcano plots showing effects of designed sequence edit on *PPIF* expression measured by Variant-FlowFISH in (**a**) THP-1 cells and (**b**) Jurkat cells. *X*-axis: Measured effect in Variant-FlowFISH. *Y*-axis: Benjamini-Hochberg corrected *p*-values from two-sided, one-sample *t*-test. Each dot represents a different edit (*N* = 164). Horizontal dotted line: adjusted p-value = 0.05.

**Supplementary Figure 20.**
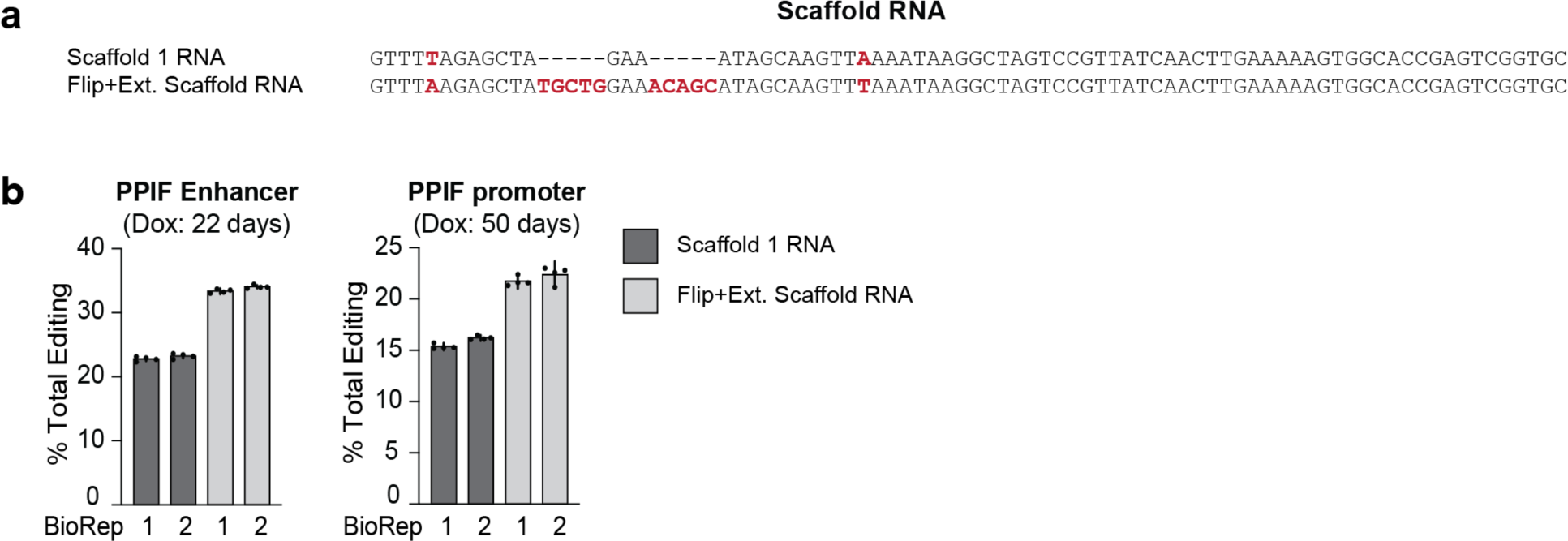
Comparing the editing rates between two different RNA scaffolds in THP-1 PE2 cells. (**a**) An alignment of the two scaffolds used in this experiment: Scaffold 1 RNA and the Flip and Extension Scaffold RNA ^96^. (**b**) A comparison between the two scaffold RNAs used during the pegRNA design, in terms of the percentage (%) of total editing seen at the PPIF enhancer and promoter in THP-1 PE2 cells, after 22-50 days in doxycycline treatment (as indicated). Shown is the mean +/-95% c.i for each of the bio-replicates. Dots represent PCR replicates (*n*=4).

**Supplementary Figure 21.**
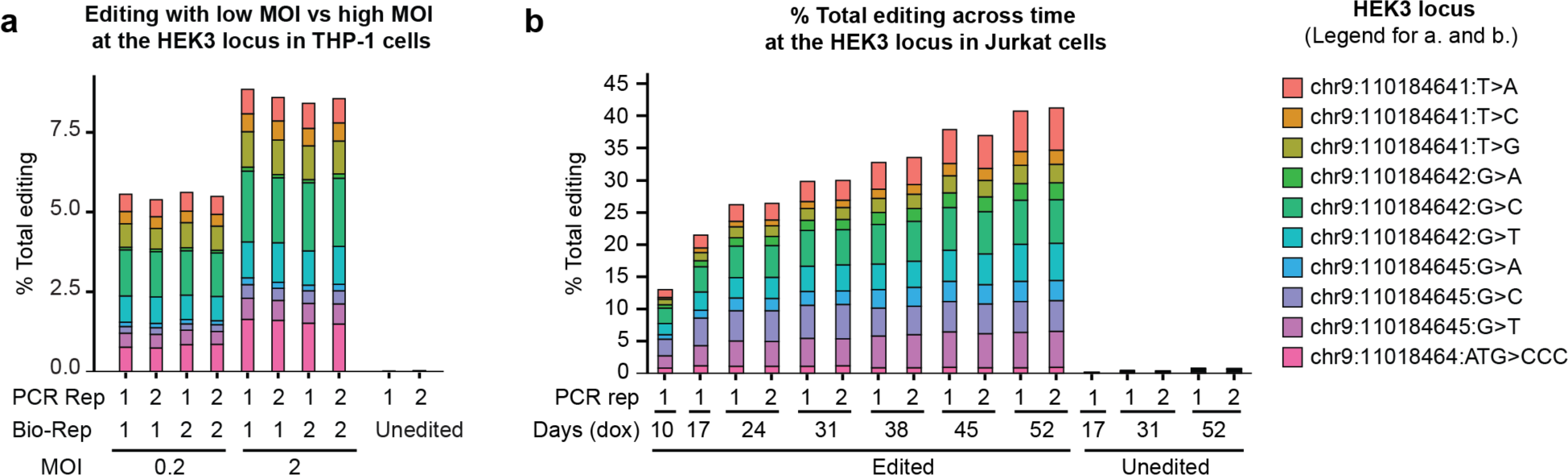
Optimizing prime editing in THP-1 PE2 and Jurkat PE2 cells. (**a**) The percentage of total editing at the HEK3 locus in THP-1 PE2 cells when testing two different multiplicity of infection (MOI) amounts (0.2 MOI vs. 2 MOI). Shown are two PCR replicates from two bio-replicates. (**b**) The percentage of total editing at the HEK3 locus in Jurkat cells, across time, as shown by the number of days of doxycycline treatment (which induces the PE2 editing machinery). 9 SNV edits and one MNV edit were introduced at the HEK3 locus, as shown in the legend. Coordinates for the HEK3 site are also shown.

**Supplementary Figure 22.**
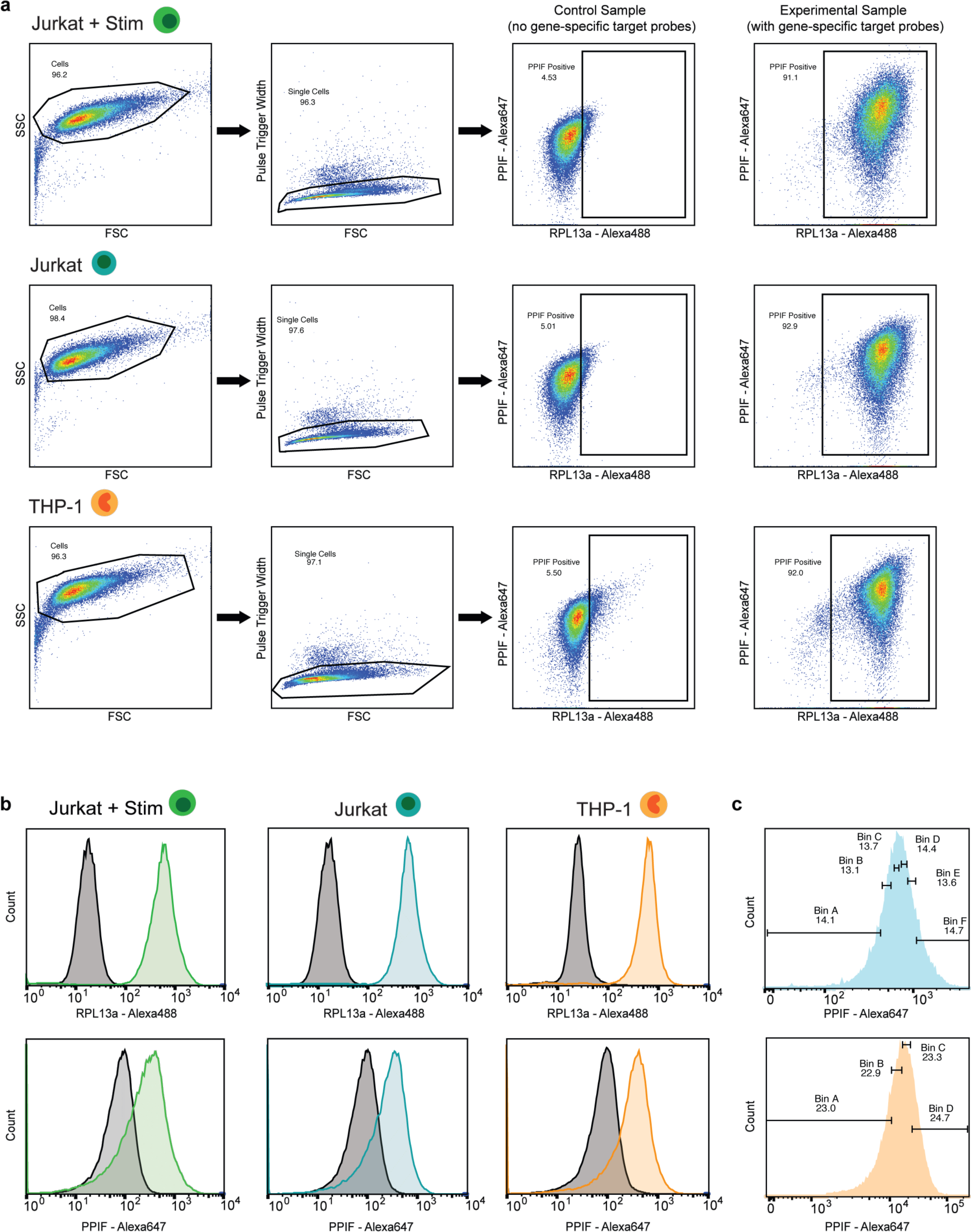
Representative FACS plots for Variant-FlowFISH screens. (**a**) Representative flow plots displaying the gating strategy implemented for each cell type studied with Variant-FlowFISH, using *PPIF* as the gene of interest and *RPL13A* as the control gene. First, cells are sorted from debris using forward scatter versus side scatter (first column). Then, we select for single cells (second column). We then use a control sample that has not been treated with gene-specific target probes (but has undergone the rest of the FlowFISH protocol), and set a gate to exclude all but the top ∼5% of control cells with the highest signal in both fluorescence channels (third column). Fourth column is a representative example of a successfully probed sample from each respective cell type. The gated region is the population of cells we sort. FSC: Forward scatter. SSC: Side scatter. Numbers on each plot represent the percentage (%) of cells included in the gate. (**b**) Representative plots of each cell type showing the distribution of fluorescence intensities from each cell. Gray: Control sample. Color: Sample treated with the indicated gene probeset. We reliably observe efficiently probed samples with both the RPL13a - Alexa488 and PPIF - Alexa647 probesets. (**c**) Sample plots of our sorting gates in either the 6 bin or 4 bin setting. When using 4 bins we placed the sorting gates next to each other and so that they cover the entire distribution (∼25% of the population captured in each gate). When using the 6-bin strategy we placed the sort gates equidistant from each leaving small gaps in between gates, and each gate captured 10-15% of the population.

## References

1. Claussnitzer, M., Cho, J.H., Collins, R., Cox, N.J., Dermitzakis, E.T., Hurles, M.E., Kathiresan, S., Kenny, E.E., Lindgren, C.M., MacArthur, D.G., et al. (2020). A brief history of human disease genetics. Nature 577. 10.1038/s41586-019-1879-7.

2. Nasser, J., Bergman, D.T., Fulco, C.P., Guckelberger, P., Doughty, B.R., Patwardhan, T.A., Jones, T.R., Nguyen, T.H., Ulirsch, J.C., Lekschas, F., et al. (2021). Genome-wide enhancer maps link risk variants to disease genes. Nature 593, 238–243.

3. Maurano, M.T., Humbert, R., Rynes, E., Thurman, R.E., Haugen, E., Wang, H., Reynolds, A.P., Sandstrom, R., Qu, H., Brody, J., et al. (2012). Systematic localization of common disease-associated variation in regulatory DNA. Science 337, 1190–1195.

4. Frangoul, H., Altshuler, D., Domenica Cappellini, M., Chen, Y.-S., Domm, J., Eustace, B.K., Foell, J., de la Fuente, J., Grupp, S., Handgretinger, R., et al. (2020). CRISPR-Cas9 Gene Editing for Sickle Cell Disease and β-Thalassemia. N. Engl. J. Med. 10.1056/NEJMoa2031054.

5. Canver, M.C., Smith, E.C., Sher, F., Pinello, L., Sanjana, N.E., Shalem, O., Chen, D.D., Schupp, P.G., Vinjamur, D.S., Garcia, S.P., et al. (2015). BCL11A enhancer dissection by Cas9-mediated in situ saturating mutagenesis. Nature 527. 10.1038/nature15521.

6. Avsec, Ž., Agarwal, V., Visentin, D., Ledsam, J.R., Grabska-Barwinska, A., Taylor, K.R., Assael, Y., Jumper, J., Kohli, P., and Kelley, D.R. (2021). Effective gene expression prediction from sequence by integrating long-range interactions. Nat. Methods 18, 1196–1203.

7. Linder, J., Srivastava, D., Yuan, H., Agarwal, V., and Kelley, D.R. (2023). Predicting RNA-seq coverage from DNA sequence as a unifying model of gene regulation. bioRxiv, 2023.08.30.555582. 10.1101/2023.08.30.555582.

8. Avsec, Ž., Weilert, M., Shrikumar, A., Krueger, S., Alexandari, A., Dalal, K., Fropf, R., McAnany, C., Gagneur, J., Kundaje, A., et al. (2021). Base-resolution models of transcription-factor binding reveal soft motif syntax. Nat. Genet. 53, 354–366.

9. Chen, K.M., Wong, A.K., Troyanskaya, O.G., and Zhou, J. (2022). A sequence-based global map of regulatory activity for deciphering human genetics. Nat. Genet. 54, 940–949.

10. Dudnyk, K., Shi, C., and Zhou, J. (2023). Sequence basis of transcription initiation in human genome. bioRxiv. 10.1101/2023.06.27.546584.

11. Agarwal, V., and Shendure, J. (2020). Predicting mRNA Abundance Directly from Genomic Sequence Using Deep Convolutional Neural Networks. Cell Rep. 31, 107663.

12. Engreitz, J.M., Haines, J.E., Perez, E.M., Munson, G., Chen, J., Kane, M., McDonel, P.E., Guttman, M., and Lander, E.S. (2016). Local regulation of gene expression by lncRNA promoters, transcription and splicing. Nature 539, 452–455.

13. Ulirsch, J.C., Nandakumar, S.K., Wang, L., Giani, F.C., Zhang, X., Rogov, P., Melnikov, A., McDonel, P., Do, R., Mikkelsen, T.S., et al. (2016). Systematic Functional Dissection of Common Genetic Variation Affecting Red Blood Cell Traits. Cell 165, 1530–1545.

14. Martyn, G.E., Wienert, B., Yang, L., Shah, M., Norton, L.J., Burdach, J., Kurita, R., Nakamura, Y., Pearson, R.C.M., Funnell, A.P.W., et al. (2018). Natural regulatory mutations elevate the fetal globin gene via disruption of BCL11A or ZBTB7A binding. Nat. Genet. 50, 498–503.

15. Melnikov, A., Murugan, A., Zhang, X., Tesileanu, T., Wang, L., Rogov, P., Feizi, S., Gnirke, A., Callan, C.G., Kinney, J.B., et al. (2012). Systematic dissection and optimization of inducible enhancers in human cells using a massively parallel reporter assay. Nat. Biotechnol. 30. 10.1038/nbt.2137.

16. Kircher, M., Xiong, C., Martin, B., Schubach, M., Inoue, F., Bell, R.J.A., Costello, J.F., Shendure, J., and Ahituv, N. (2019). Saturation mutagenesis of twenty disease-associated regulatory elements at single base-pair resolution. Nat. Commun. 10, 1–15.

17. Grossman, S.R., Zhang, X., Wang, L., Engreitz, J., Melnikov, A., Rogov, P., Tewhey, R., Isakova, A., Deplancke, B., Bernstein, B.E., et al. (2017). Systematic dissection of genomic features determining transcription factor binding and enhancer function. Proc. Natl. Acad. Sci. U. S. A. 114, E1291–E1300.

18. Kwasnieski, J.C., Mogno, I., Myers, C.A., Corbo, J.C., and Cohen, B.A. (2012). Complex effects of nucleotide variants in a mammalian cis-regulatory element. Proc. Natl. Acad. Sci. U. S. A. 109, 19498–19503.

19. White, M.A., Kwasnieski, J.C., Myers, C.A., Shen, S.Q., Corbo, J.C., and Cohen, B.A. (2016). A Simple Grammar Defines Activating and Repressing cis-Regulatory Elements in Photoreceptors. Cell Rep. 17, 1247–1254.

20. Gamma-Globin Gene Promoter Elements Required for Interaction With Globin Enhancers (1998). Blood 91, 309–318.

21. The enhanceosome (2008). Curr. Opin. Struct. Biol. 18, 236–242.

22. King, D.M., Hong, C.K.Y., Shepherdson, J.L., Granas, D.M., Maricque, B.B., and Cohen, B.A. (2020). Synthetic and genomic regulatory elements reveal aspects of cis-regulatory grammar in mouse embryonic stem cells. 10.7554/eLife.41279.

23. Sanjana, N.E., Wright, J., Zheng, K., Shalem, O., Fontanillas, P., Joung, J., Cheng, C., Regev, A., and Zhang, F. (2016). High-resolution interrogation of functional elements in the noncoding genome. Science 353. 10.1126/science.aaf7613.

24. Martin-Rufino, J.D., Castano, N., Pang, M., Grody, E.I., Joubran, S., Caulier, A., Wahlster, L., Li, T., Qiu, X., Riera-Escandell, A.M., et al. (2023). Massively parallel base editing to map variant effects in human hematopoiesis. Cell 186, 2456–2474.e24.

25. Chen, Z., Javed, N., Moore, M., Wu, J., Sun, G., Vinyard, M., Collins, A., Pinello, L., Najm, F.J., and Bernstein, B.E. (2023). Integrative dissection of gene regulatory elements at base resolution. Cell Genom 3, 100318.

26. Morris, J.A., Caragine, C., Daniloski, Z., Domingo, J., Barry, T., Lu, L., Davis, K., Ziosi, M., Glinos, D.A., Hao, S., et al. (2023). Discovery of target genes and pathways at GWAS loci by pooled single-cell CRISPR screens. Science 380, eadh7699.

27. Cuella-Martin, R., Hayward, S.B., Fan, X., Chen, X., Huang, J.-W., Taglialatela, A., Leuzzi, G., Zhao, J., Rabadan, R., Lu, C., et al. (2021). Functional interrogation of DNA damage response variants with base editing screens. Cell 184, 1081–1097.e19.

28. Cheng, L., Li, Y., Qi, Q., Xu, P., Feng, R., Palmer, L., Chen, J., Wu, R., Yee, T., Zhang, J., et al. (2021). Single-nucleotide-level mapping of DNA regulatory elements that control fetal hemoglobin expression. Nat. Genet. 53, 869–880.

29. Diao, Y., Li, B., Meng, Z., Jung, I., Lee, A.Y., Dixon, J., Maliskova, L., Guan, K.-L., Shen, Y., and Ren, B. (2016). A new class of temporarily phenotypic enhancers identified by CRISPR/Cas9-mediated genetic screening. Genome Res. 26, 397–405.

30. Rajagopal, N., Srinivasan, S., Kooshesh, K., Guo, Y., Edwards, M.D., Banerjee, B., Syed, T., Emons, B.J.M., Gifford, D.K., and Sherwood, R.I. (2016). High-throughput mapping of regulatory DNA. Nat. Biotechnol. 34, 167–174.

31. Korkmaz, G., Lopes, R., Ugalde, A.P., Nevedomskaya, E., Han, R., Myacheva, K., Zwart, W., Elkon, R., and Agami, R. (2016). Functional genetic screens for enhancer elements in the human genome using CRISPR-Cas9. Nat. Biotechnol. 34, 192–198.

32. Ryu, J., Barkal, S., Yu, T., Jankowiak, M., Zhou, Y., Francoeur, M., Phan, Q.V., Li, Z., Tognon, M., Brown, L., et al. (2023). Joint genotypic and phenotypic outcome modeling improves base editing variant effect quantification. medRxiv, 2023.09.08.23295253. 10.1101/2023.09.08.23295253.

33. Anzalone, A.V., Randolph, P.B., Davis, J.R., Sousa, A.A., Koblan, L.W., Levy, J.M., Chen, P.J., Wilson, C., Newby, G.A., Raguram, A., et al. (2019). Search-and-replace genome editing without double-strand breaks or donor DNA. Nature 576, 149–157.

34. Ren, X., Yang, H., Nierenberg, J.L., Sun, Y., Chen, J., Beaman, C., Pham, T., Nobuhara, M., Takagi, M.A., Narayan, V., et al. (2023). High throughput PRIME editing screens identify functional DNA variants in the human genome. bioRxiv, 2023.07.12.548736. 10.1101/2023.07.12.548736.

35. Chardon, F.M., Suiter, C.C., Daza, R.M., Smith, N.T., Parrish, P., McDiarmid, T., Lalanne, J.-B., Martin, B., Calderon, D., Ellison, A., et al. (2023). A multiplex, prime editing framework for identifying drug resistance variants at scale. bioRxiv, 2023.07.27.550902. 10.1101/2023.07.27.550902.

36. Erwood, S., Bily, T.M.I., Lequyer, J., Yan, J., Gulati, N., Brewer, R.A., Zhou, L., Pelletier, L., Ivakine, E.A., and Cohn, R.D. (2022). Saturation variant interpretation using CRISPR prime editing. Nat. Biotechnol. 40, 885–895.

37. Fulco, C.P., Nasser, J., Jones, T.R., Munson, G., Bergman, D.T., Subramanian, V., Grossman, S.R., Anyoha, R., Doughty, B.R., Patwardhan, T.A., et al. (2019). Activity-by-contact model of enhancer-promoter regulation from thousands of CRISPR perturbations. Nat. Genet. 51, 1664–1669.

38. ORFs, Splicing & Coding (2016). Gene Infinity.Org. http://www.geneinfinity.org/sp/sp_coding.html.

39. Nelson, J.W., Randolph, P.B., Shen, S.P., Everette, K.A., Chen, P.J., Anzalone, A.V., An, M., Newby, G.A., Chen, J.C., Hsu, A., et al. (2021). Engineered pegRNAs improve prime editing efficiency. Nat. Biotechnol., 1–9.

40. Kim, H.K., Yu, G., Park, J., Min, S., Lee, S., Yoon, S., and Kim, H.H. (2021). Predicting the efficiency of prime editing guide RNAs in human cells. Nat. Biotechnol. 39, 198–206.

41. Li, X., Chen, W., Martin, B.K., Calderon, D., Lee, C., Choi, J., Chardon, F.M., McDiarmid, T., Kim, H., Lalanne, J.-B., et al. (2023). Chromatin context-dependent regulation and epigenetic manipulation of prime editing. bioRxiv, 2023.04.12.536587. 10.1101/2023.04.12.536587.

42. Pampari, A., Shcherbina, A., Nair, S., Schreiber, J., Patel, A., Wang, A., Kundu, S., Shrikumar, A., and Kundaje, A. Bias factorized, base-resolution deep learning models of chromatin accessibility reveal cis-regulatory sequence syntax, transcription factor footprints and regulatory variants. In preparation.

43. Bernardi, P., and Di Lisa, F. (2015). The mitochondrial permeability transition pore: Molecular nature and role as a target in cardioprotection. J. Mol. Cell. Cardiol. 78, 100.

44. Shrikumar, A., Greenside, P., and Kundaje, A. (06--11 Aug 2017). Learning Important Features Through Propagating Activation Differences. In Proceedings of the 34th International Conference on Machine Learning Proceedings of Machine Learning Research., D. Precup and Y. W. Teh, eds. (PMLR), pp. 3145–3153.

45. Lundberg, S.M., and Lee, S.-I. (2017). A Unified Approach to Interpreting Model Predictions. Adv. Neural Inf. Process. Syst. 30.

46. Kvon, E.Z., Zhu, Y., Kelman, G., Novak, C.S., Plajzer-Frick, I., Kato, M., Garvin, T.H., Pham, Q., Harrington, A.N., Hunter, R.D., et al. (2020). Comprehensive In Vivo Interrogation Reveals Phenotypic Impact of Human Enhancer Variants. Cell 180, 1262–1271.e15.

47. Farley, E.K., Olson, K.M., Zhang, W., Brandt, A.J., Rokhsar, D.S., and Levine, M.S. (2015). Suboptimization of developmental enhancers. Science 350. 10.1126/science.aac6948.

48. Computational Models for Neurogenic Gene Expression in the Drosophila Embryo (2006). Curr. Biol. 16, 1358–1365.

49. Erceg, J., Saunders, T.E., Girardot, C., Devos, D.P., Hufnagel, L., and Furlong, E.E.M. (2014). Subtle Changes in Motif Positioning Cause Tissue-Specific Effects on Robustness of an Enhancer’s Activity. PLoS Genet. 10, e1004060.

50. Levo, M., and Segal, E. (2014). In pursuit of design principles of regulatory sequences. Nat. Rev. Genet. 15, 453–468.

51. Low Affinity Binding Site Clusters Confer Hox Specificity and Regulatory Robustness (2015). Cell 160, 191–203.

52. Structural Rules and Complex Regulatory Circuitry Constrain Expression of a Notch- and EGFR-Regulated Eye Enhancer (2010). Dev. Cell 18, 359–370.

53. Snetkova, V., Ypsilanti, A.R., Akiyama, J.A., Mannion, B.J., Plajzer-Frick, I., Novak, C.S., Harrington, A.N., Pham, Q.T., Kato, M., Zhu, Y., et al. (2021). Ultraconserved enhancer function does not require perfect sequence conservation. Nat. Genet. 53, 521–528.

54. Galupa, R., Alvarez-Canales, G., Borst, N.O., Fuqua, T., Gandara, L., Misunou, N., Richter, K., Alves, M.R.P., Karumbi, E., Perkins, M.L., et al. (2023). Enhancer architecture and chromatin accessibility constrain phenotypic space during Drosophila development. Dev. Cell 58, 51–62.e4.

55. Sahu, B., Hartonen, T., Pihlajamaa, P., Wei, B., Dave, K., Zhu, F., Kaasinen, E., Lidschreiber, K., Lidschreiber, M., Daub, C.O., et al. (2022). Sequence determinants of human gene regulatory elements. Nat. Genet. 54, 283–294.

56. Rauluseviciute, I., Riudavets-Puig, R., Blanc-Mathieu, R., Castro-Mondragon, J.A., Ferenc, K., Kumar, V., Lemma, R.B., Lucas, J., Chèneby, J., Baranasic, D., et al. (2023). JASPAR 2024: 20th anniversary of the open-access database of transcription factor binding profiles. Nucleic Acids Res. 10.1093/nar/gkad1059.

57. Bailey, T.L., Johnson, J., Grant, C.E., and Noble, W.S. (2015). The MEME Suite. Nucleic Acids Res. 43, W39–W49.

58. Shrikumar, A., Tian, K., Avsec, Ž., Shcherbina, A., Banerjee, A., Sharmin, M., Nair, S., and Kundaje, A. (2018). Technical Note on Transcription Factor Motif Discovery from Importance Scores (TF-MoDISco) version 0.5.6.5. arXiv [cs.LG].

59. Wang, T., Birsoy, K., Hughes, N.W., Krupczak, K.M., Post, Y., Wei, J.J., Lander, E.S., and Sabatini, D.M. (2015). Identification and characterization of essential genes in the human genome. Science 350, 1096–1101.

60. Bergman, D.T., Jones, T.R., Liu, V., Ray, J., Jagoda, E., Siraj, L., Kang, H.Y., Nasser, J., Kane, M., Rios, A., et al. (2022). Compatibility rules of human enhancer and promoter sequences. Nature 607, 176–184.

61. Weingarten-Gabbay, S., Nir, R., Lubliner, S., Sharon, E., Kalma, Y., Weinberger, A., and Segal, E. (2019). Systematic interrogation of human promoters. Genome Research 29, 171–183. 10.1101/gr.236075.118.

62. Yu, M., Yang, X.Y., Schmidt, T., Chinenov, Y., Wang, R., and Martin, M.E. (1997). GA-binding protein-dependent transcription initiator elements. Effect of helical spacing between polyomavirus enhancer a factor 3(PEA3)/Ets-binding sites on initiator activity. J. Biol. Chem. 272, 29060–29067.

63. Deen, D., Butter, F., Daniels, D.E., Ferrer-Vicens, I., Ferguson, D.C.J., Holland, M.L., Samara, V., Sloane-Stanley, J.A., Ayyub, H., Mann, M., et al. (2021). Identification of the transcription factor MAZ as a regulator of erythropoiesis. Blood Adv 5, 3002–3015.

64. Her, S., Claycomb, R., Tai, T.C., and Wong, D.L. (2003). Regulation of the rat phenylethanolamine N-methyltransferase gene by transcription factors Sp1 and MAZ. Mol. Pharmacol. 64, 1180–1188.

65. Kurisaki, K., Kurisaki, A., Valcourt, U., Terentiev, A.A., Pardali, K., ten Dijke, P., Heldin, C.-H., Ericsson, J., and Moustakas, A. (2003). Nuclear Factor YY1 Inhibits Transforming Growth Factor β- and Bone Morphogenetic Protein-Induced Cell Differentiation. Mol. Cell. Biol. 23, 4494–4510.

66. Yao, Y.L., Yang, W.M., and Seto, E. (2001). Regulation of transcription factor YY1 by acetylation and deacetylation. Mol. Cell. Biol. 21, 5979–5991.

67. Walz, S., Lorenzin, F., Morton, J., Wiese, K.E., von Eyss, B., Herold, S., Rycak, L., Dumay-Odelot, H., Karim, S., Bartkuhn, M., et al. (2014). Activation and repression by oncogenic MYC shape tumour-specific gene expression profiles. Nature 511, 483–487.

68. Staller, P., Peukert, K., Kiermaier, A., Seoane, J., Lukas, J., Karsunky, H., Möröy, T., Bartek, J., Massagué, J., Hänel, F., et al. (2001). Repression of p15INK4b expression by Myc through association with Miz-1. Nat. Cell Biol. 3, 392–399.

69. Xiao, T., Li, X., and Felsenfeld, G. (2021). The Myc-associated zinc finger protein (MAZ) works together with CTCF to control cohesin positioning and genome organization. Proc. Natl. Acad. Sci. U. S. A. 118. 10.1073/pnas.2023127118.

70. Izumi, H., Molander, C., Penn, L.Z., Ishisaki, A., Kohno, K., and Funa, K. (2001). Mechanism for the transcriptional repression by c-Myc on PDGF beta-receptor. J. Cell Sci. 114, 1533–1544.

71. Bennett, M.K., Ngo, T.T., Athanikar, J.N., Rosenfeld, J.M., and Osborne, T.F. (1999). Co-stimulation of promoter for low density lipoprotein receptor gene by sterol regulatory element-binding protein and Sp1 is specifically disrupted by the yin yang 1 protein. J. Biol. Chem. 274, 13025–13032.

72. Yukawa, M., Jagannathan, S., Vallabh, S., Kartashov, A.V., Chen, X., Weirauch, M.T., and Barski, A. (2020). AP-1 activity induced by co-stimulation is required for chromatin opening during T cell activation. J. Exp. Med. 217. 10.1084/jem.20182009.

73. Cochran, K., Schreiber, J., Yin, M., Mantripragada, A., Marinov, G., Yu, H., Lis, J., and Kundaje, A. ProCapNet: Dissecting the cis-regulatory syntax of transcription initiation with deep learning. In preparation.

74. Soufi, A., Donahue, G., and Zaret, K.S. (2012). Facilitators and impediments of the pluripotency reprogramming factors’ initial engagement with the genome. Cell 151. 10.1016/j.cell.2012.09.045.

75. Nair, S., Ameen, M., Sundaram, L., Pampari, A., Schreiber, J., Balsubramani, A., Wang, Y.X., Burns, D., Blau, H.M., Karakikes, I., et al. (2023). Transcription factor stoichiometry, motif affinity and syntax regulate single-cell chromatin dynamics during fibroblast reprogramming to pluripotency. bioRxiv. 10.1101/2023.10.04.560808.

76. Linder, J., and Seelig, G. (2021). Fast activation maximization for molecular sequence design. BMC Bioinformatics 22, 510.

77. Anishchenko, I., Pellock, S.J., Chidyausiku, T.M., Ramelot, T.A., Ovchinnikov, S., Hao, J., Bafna, K., Norn, C., Kang, A., Bera, A.K., et al. (2021). De novo protein design by deep network hallucination. Nature 600, 547–552.

78. de Almeida, B.P., Schaub, C., Pagani, M., Secchia, S., Furlong, E.E.M., and Stark, A. (2023). Targeted design of synthetic enhancers for selected tissues in the Drosophila embryo. Nature. 10.1038/s41586-023-06905-9.

79. Taskiran, I.I., Spanier, K.I., Dickmänken, H., Kempynck, N., Pančíková, A., Ekşi, E.C., Hulselmans, G., Ismail, J.N., Theunis, K., Vandepoel, R., et al. (2023). Cell type directed design of synthetic enhancers. Nature, 1–3.

80. Gosai, S.J., Castro, R.I., Fuentes, N., Butts, J.C., Kales, S., Noche, R.R., Mouri, K., Sabeti, P.C., Reilly, S.K., and Tewhey, R. (2023). Machine-guided design of synthetic cell type-specific cis-regulatory elements. bioRxiv. 10.1101/2023.08.08.552077.

81. de Almeida, B.P., Reiter, F., Pagani, M., and Stark, A. (2022). DeepSTARR predicts enhancer activity from DNA sequence and enables the de novo design of synthetic enhancers. Nat. Genet. 54, 613–624.

82. Kryshtafovych, A., Schwede, T., Topf, M., Fidelis, K., and Moult, J. (2021). Critical assessment of methods of protein structure prediction (CASP)-Round XIV. Proteins 89, 1607–1617.

83. Gschwind, A.R., Mualim, K.S., Karbalayghareh, A., Sheth, M.U., Dey, K.K., Jagoda, E., Nurtdinov, R.N., Xi, W., Tan, A.S., Jones, H., et al. (2023). An encyclopedia of enhancer-gene regulatory interactions in the human genome. bioRxiv, 2023.11.09.563812. 10.1101/2023.11.09.563812.

84. Karollus, A., Mauermeier, T., and Gagneur, J. (2023). Current sequence-based models capture gene expression determinants in promoters but mostly ignore distal enhancers. Genome Biol. 24, 56.

85. Priber, J., Fonai, F., Jakus, P.B., Racz, B., Chinopoulos, C., Tretter, L., Gallyas, F., Jr, Sumegi, B., and Veres, B. (2015). Cyclophilin D disruption attenuates lipopolysaccharide-induced inflammatory response in primary mouse macrophages. Biochem. Cell Biol. 93, 241–250.

86. Piot, C., Croisille, P., Staat, P., Thibault, H., Rioufol, G., Mewton, N., Elbelghiti, R., Cung, T.T., Bonnefoy, E., Angoulvant, D., et al. (2008). Effect of Cyclosporine on Reperfusion Injury in Acute Myocardial Infarction. 10.1056/NEJMoa071142.

87. Panel, M., Ruiz, I., Brillet, R., Lafdil, F., Teixeira-Clerc, F., Nguyen, C.T., Calderaro, J., Gelin, M., Allemand, F., Guichou, J.-F., et al. (2019). Small-Molecule Inhibitors of Cyclophilins Block Opening of the Mitochondrial Permeability Transition Pore and Protect Mice From Hepatic Ischemia/Reperfusion Injury. Gastroenterology 157, 1368–1382.

88. Matharu, N., Rattanasopha, S., Tamura, S., Maliskova, L., Wang, Y., Bernard, A., Hardin, A., Eckalbar, W.L., Vaisse, C., and Ahituv, N. (2019). CRISPR-mediated activation of a promoter or enhancer rescues obesity caused by haploinsufficiency. Science 363. 10.1126/science.aau0629.

89. Tamura, S., Nelson, A.D., Spratt, P.W.E., Kyoung, H., Zhou, X., Li, Z., Zhao, J., Holden, S.S., Sahagun, A., Keeshen, C.M., et al. (2022). CRISPR activation rescues abnormalities in SCN2A haploinsufficiency-associated autism spectrum disorder. bioRxiv, 2022.03.30.486483. 10.1101/2022.03.30.486483.

90. Koeppel, J., Weller, J., Peets, E.M., Pallaseni, A., Kuzmin, I., Raudvere, U., Peterson, H., Liberante, F.G., and Parts, L. (2023). Prediction of prime editing insertion efficiencies using sequence features and DNA repair determinants. Nat. Biotechnol. 41, 1446–1456.

91. Reilly, S.K., Gosai, S.J., Gutierrez, A., Mackay-Smith, A., Ulirsch, J.C., Kanai, M., Mouri, K., Berenzy, D., Kales, S., Butler, G.M., et al. (2021). Direct characterization of cis-regulatory elements and functional dissection of complex genetic associations using HCR–FlowFISH. Nat. Genet. 53, 1166– 1176.

92. Hsu, J.Y., Grünewald, J., Szalay, R., Shih, J., Anzalone, A.V., Lam, K.C., Shen, M.W., Petri, K., Liu, D.R., Joung, J.K., et al. (2021). PrimeDesign software for rapid and simplified design of prime editing guide RNAs. Nat. Commun. 12, 1034.

93. O’Leary, N.A., Wright, M.W., Brister, J.R., Ciufo, S., Haddad, D., McVeigh, R., Rajput, B., Robbertse, B., Smith-White, B., Ako-Adjei, D., et al. (2016). Reference sequence (RefSeq) database at NCBI: current status, taxonomic expansion, and functional annotation. Nucleic Acids Res. 44, D733–D745.

94. Khan, A., Fornes, O., Stigliani, A., Gheorghe, M., Castro-Mondragon, J.A., van der Lee, R., Bessy, A., Chèneby, J., Kulkarni, S.R., Tan, G., et al. (2018). JASPAR 2018: update of the open-access database of transcription factor binding profiles and its web framework. Nucleic Acids Res. 46, D1284.

95. Hsu, P.D., Scott, D.A., Weinstein, J.A., Ran, F.A., Konermann, S., Agarwala, V., Li, Y., Fine, E.J., Wu, X., Shalem, O., et al. (2013). DNA targeting specificity of RNA-guided Cas9 nucleases. Nat. Biotechnol. 31, 827–832.

96. Chen, B., Gilbert, L.A., Cimini, B.A., Schnitzbauer, J., Zhang, W., Li, G.-W., Park, J., Blackburn, E.H., Weissman, J.S., Qi, L.S., et al. (2013). Dynamic imaging of genomic loci in living human cells by an optimized CRISPR/Cas system. Cell 155, 1479–1491.

97. Clement, K., Rees, H., Canver, M.C., Gehrke, J.M., Farouni, R., Hsu, J.Y., Cole, M.A., Liu, D.R., Joung, J.K., Bauer, D.E., et al. (2019). CRISPResso2 provides accurate and rapid genome editing sequence analysis. Nat. Biotechnol. 37, 224–226.

98. Karimzadeh, M., Ernst, C., Kundaje, A., and Hoffman, M.M. (2018). Umap and Bismap: quantifying genome and methylome mappability. Nucleic Acids Res. 46, e120.

99. Chardon FM, Suiter CC, Daza RM, Smith NT, Parrish P, McDiarmid T, Lalanne JB, Martin B, Calderon D, Ellison A, Berger AH, Shendure J, Starita LM (2023). A multiplex, prime editing framework for identifying drug resistance variants at scale. bioRxiv.

